# FuzzAletheia: Mapping dynamic interaction landscapes of intrinsically disordered protein-ligand complexes to accelerate ligand optimization

**DOI:** 10.1101/2025.11.26.690418

**Authors:** Hanping Wang, Fanhao Wang, Ruoyao Xiong, Qi Sun, Luhua Lai

## Abstract

Intrinsically disordered proteins (IDPs), representing more than 30% of human proteome, present great challenges in rational drug design due to their highly flexible conformations and dynamic interactions. Though their malfunction is linked to major diseases, IDPs are difficult to target. A critical roadblock lies in hit-to-lead optimization due to their dynamic interactions with hit compounds. Here we developed FuzzAletheia, an IDP-ligand dynamic binding pattern analyzer, to accelerate hit optimization. It analyzes the molecular dynamics simulation trajectories of IDP-ligand to cluster interactions into distinct spatiotemporal stable binding features, which are then visualized and quantified to provide insights into the dynamic binding orientation, strength, and stability. This analysis pinpoints modification sites on hit compounds for optimization. We applied FuzzAletheia to the disordered p53 transactivation domain 1 and its hit compound **1050.** Guided by FuzzAletheia analysis, we designed and synthesized six derivatives of **1050** and the most potent compound achieved a 10-fold enhancement in activity, demonstrating its power to accelerate IDP hit optimization.

## Introduction

Intrinsically disordered proteins (IDPs) are proteins that lack well-defined structures under physiological conditions. The structural dynamics enables IDPs to function as protein-protein interaction network hubs (*1, 2*) and to form membraneless organelles (*3, 4*). Comprising over 30% of the human proteome (*5, 6*), IDPs play crucial roles in processes such as signal transduction (*7, 8*) and transcription regulation (*9, 10*). Due to their significant role in regulating cell fate and their involvement in major diseases, IDPs have been regarded as the “most wanted” targets for drug discovery (*11, 12*). A number of compounds that regulate IDP functions via directly binding to the disordered regions have been identified (*12*). Phase I clinical trial results of Masofaniten (*13*) and OMO-103 (*14*) demonstrate that directly targeting disordered regions is a safe and potentially effective strategy for cancer treatment, highlighting the promise of IDP-targeted drugs as a novel therapeutic avenue. However, the lengthy journey to bring IDP ligands to the clinic (*15, 16*), along with their scarcity (*12*), highlights IDP ligand optimization as a critical and urgent challenge in IDP drug discovery.

This challenge arises from the unique binding characteristics of IDP-ligand complexes. Unlike ordered complexes with well-defined binding patterns, IDP-ligand complexes exhibit “fuzziness”, a dynamic behaviour characterized by continuous transitions between multiple conformations, making experimental studies difficult. The fuzzy IDP-ligand binding ensemble comprises multiple binding patterns defined by distinct binding features, including binding orientation, interaction types, and spatiotemporal stability. Lack of a well-defined binding pattern renders the conventional structure-based lead optimization method unsuitable for IDP ligands (*17, 18*). Furthermore, the binding of small molecules can induce IDP conformational ensemble shift (*17, 19*). Chemical structure modifications of small-molecule ligands often lead to altered binding patterns, making structure-activity relationship (SAR)-based optimization inefficient or even ineffective (*20, 21*). Fortunately, it has been demonstrated that small molecules bind to IDPs mainly through certain key residues (*22, 23*). This suggests that not all binding patterns contribute equally to the overall interaction, with some playing more critical roles. Elucidating the characteristics of these predominant binding patterns will offer valuable insights for IDP ligand optimization.

The development of molecular force fields and simulation methods has made molecular dynamics (MD) simulation a powerful tool to obtain atomic-level structural details of heterogeneous IDP-ligand interaction ensembles, the information that is beyond the reach of current experimental approaches (*24-27*). Robustelli et al. (*18*) simulated the α-synuclein-fasudil system and found that tyrosine-glutamate residue pairs orient fasudil through aromatic stacking and charge-charge interactions to induce more residues to form additional interactions. Their findings imply the importance of binding orientation and anchor interactions in IDP-ligand complexes. Zhu et al. (*17*) simulated androgen receptor Tau5 domain (AR-Tau5) with EPI-series ligands using replica exchange with solute tempering (REST2) (*28*), compared their overall binding features, and found that ligands with better activity could induce AR-Tau5 to form more helical structures. These studies suggest that MD simulations can be used to guide IDP ligand optimization. However, currently available MD-based optimization methods require extensive MD simulations for each IDP-ligand complex to calculate the relative affinities between compounds, which is extremely time-consuming. Furthermore, MD-based binding mechanism analyses usually average binding features across trajectories and give statistical pictures for the overall prominent interactions, which obscures the features of distinct binding patterns and makes it difficult to assess the contribution of each pattern, thereby limiting its effectiveness in guiding ligand optimization.

The transcription factor p53 is a critical tumor suppressor that prevents tumor initiation and progression through various mechanisms (*29, 30*) and has been considered as a promising target for cancer therapy. Current strategies for p53-focused drug discovery primarily involve two types of mechanisms (*31-33*): restoring the wild-type function of mutant p53 and activating p53 by inhibiting its negative regulator MDM2. However, clinical studies have revealed that MDM2 inhibitors often exhibit low tumor-suppressive efficacy and cause significant dose-dependent side effects, underscoring the urgent need for alternative approaches to activate p53 (*32*). We have previously developed a hierarchical computational strategy for IDP drug virtual screening (IDPDVS) and applied it to discover two p53 activating molecules, **1047** and **1050**, that bind to the disordered p53 transactivation domain 1 (p53TAD1) (*34*), presenting a new pharmacological strategy for p53 activation. Despite being effective, the potency of these compounds needs to be improved.

In the present study, we developed a novel approach, FuzzAletheia, to extract IDP-ligand binding patterns and specify key binding features of each pattern in the fuzzy complex to guide ligand modification site selection, thereby accelerating IDP ligand optimization. FuzzAletheia quantifies binding features to effectively cluster complex structures into distinct binding patterns. Then, these binding patterns are visualized as point clouds to offer an easy way to understand binding orientation with spatial stability. For each binding pattern, its ligand substructure-residue interaction probabilities are calculated to quantify both interaction strength and temporal stability. Based on the utilization of these detailed binding features and the assumption that stabilizing unstable interactions in existing patterns can enhance the affinity, it is possible to realize the rapid selection of IDP ligand modification sites to design better derivatives. We have used FuzzAletheia to reveal p53TAD1-**1050** predominant binding patterns and identify potential modification sites. Guided by these insights, we only synthesized six **1050** derivatives to achieve up to 10-fold activity enhancement. Further analysis revealed that the derivative can introduce new interactions while maintaining the original binding patterns to strengthen its binding ability. We applied the same analysis to analyze the REST2 trajectories of AR-Tau5 with compounds **EPI-002** and **iodoEPI-002** to verify the general applicability of FuzzAletheia. This work provides a powerful tool for accelerating IDP ligand optimization and unraveling the binding mechanisms of fuzzy complexes.

## Computational Methods and Evaluation

### Ensemble Generation for the p53TAD1-1050 Complex

The conformational ensembles of the systems studied were obtained through MD simulations. We performed a total of 1.2 µs unbiased all-atom MD simulations of apo p53TAD1 (residues 1-39) using the Charmm36m (*35*) force field with the recommended TIP3P water model and 1.2 µs MD simulations of the p53TAD1-**1050** complex with the ligand parameterized by CGenFF (*36*). Four previously selected druggable conformations (*34*) (Figure S1) were used as the initial structures for the complex simulations. The simulation snapshots were output every 10 ps, leading to about 120,000 structures for each system.

### Overview of the FuzzAletheia Method

The overall binding feature analysis, which averages residue-ligand interaction probability and orientation of residues across the ensemble studied, cannot give a specific description required for guiding ligand optimization. To address this issue, we developed the FuzzAletheia method to extract and quantify predominant IDP-ligand binding patterns. FuzzAletheia originates from the observation that in a stable binding pattern, the ligand and its binding residues constrain mutual positions for a period of time. Methodologically, FuzzAletheia defines a stable binding pattern, also termed the predominant binding pattern, as a binding pattern in which multiple residues (at least four here) interact simultaneously with the ligand for a period of time (at least 0.8 ns here). This definition allows FuzzAletheia to extract and quantify binding patterns in fuzzy complexes by well-defined binding features, including binding orientation, interaction types, energy distribution, and spatiotemporal stability.

As illustrated in Figure 1A-C, FuzzAletheia includes five steps: 1) **Conformation alignment based on the ligand.** The conformations in the ensemble are aligned by superposing the ligand structures. 2) **Binding residue spatial distribution capturing.** First, each residue in the aligned conformation ensemble is represented by its geometric center. Then binding residue spatial distribution can be captured by keeping center points from residues engaged in stable binding (those aggregated into point clouds) and discarding those in weak or negligible binding (dispersed points). 3) **Point cloud clustering.** The point clouds of each residue are clustered using DBSCAN (Density-Based Spatial Clustering of Applications with Noise) (*37*). The clustered point clouds indicate that the corresponding residue is spatially constrained in these structures by binding to the ligand, either directly or through neighboring residues. 4) **Binding pattern extraction.** The predominant binding patterns are extracted based on point clouds and residue-ligand contact probability. The residue-ligand contact probability determines whether the point cloud results from direct residue-ligand interactions. For the p53TAD1-**1050** system, complex structures in which at least four residues continuously bind the ligand for at least 0.8 ns were collected as a predominant binding pattern. 5) **Binding pattern quantification and visualization.** In an IDP-ligand predominant binding pattern, the binding features of a residue and the ligand evolve over time, making it difficult to generalize using a single conformation. We therefore represent the predominant binding patterns through conformational ensembles rather than single snapshots. Figure 1C illustrates these patterns using: i) point cloud plots depicting orientation information with spatial stability, and ii) interaction probability plots quantifying four kinds of interaction types (hydrophobic, polar, H-bond, and π-π) with temporal stability. For better visualization of binding orientation, FuzzAletheia allows users to define a new coordinate system based on the understanding of the ligand structure.

**Figure 1.**
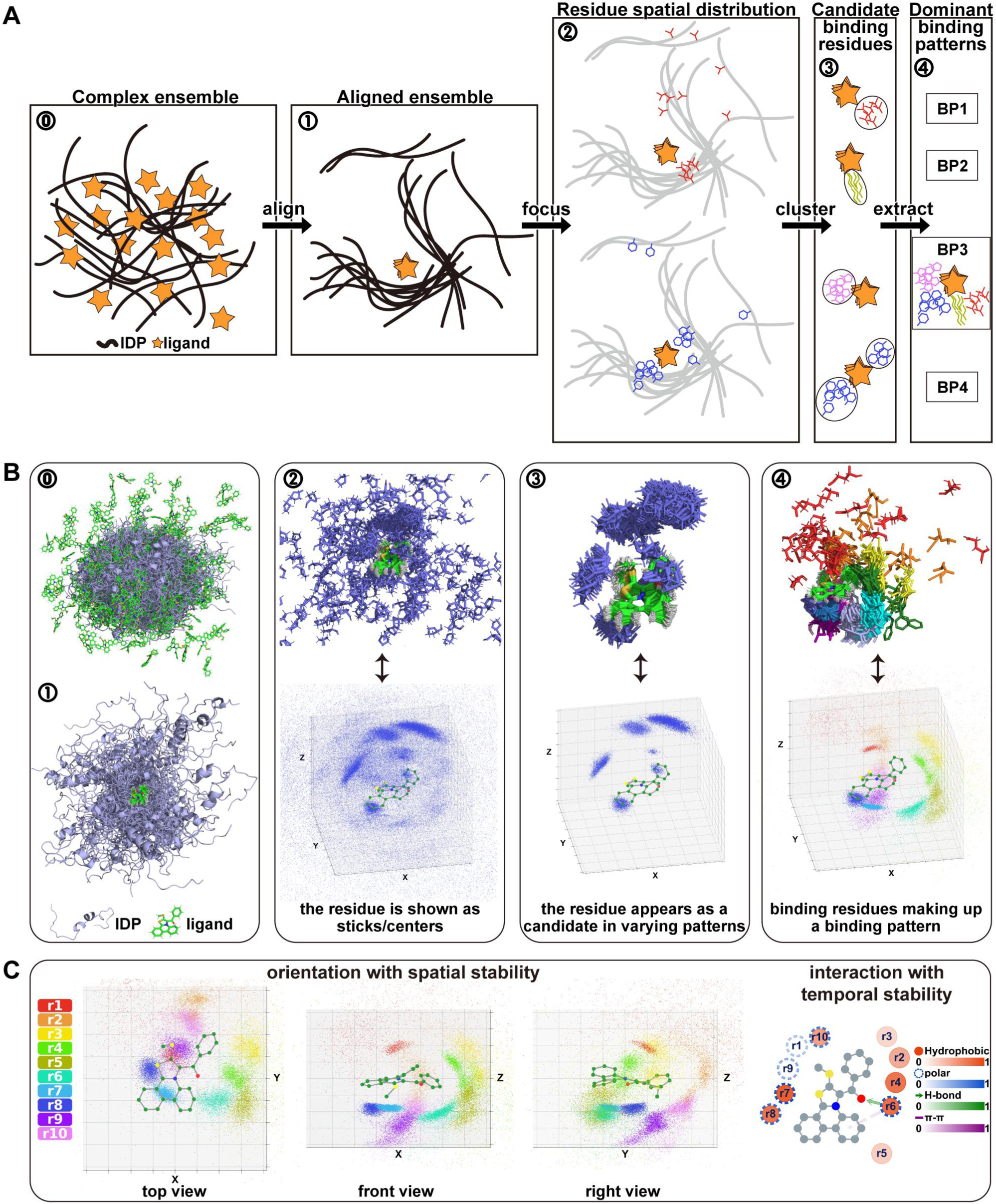
Illustration of the process that FuzzAletheia extracts and describes disordered protein-ligand predominant binding patterns. **A**. Schematic illustration of the FuzzAletheia method. MD trajectories are 1) aligned based on the ligand to 2) capture residue spatial distribution, then 3) candidate binding residues are clustered out and 4) their binding features are used to extract predominant binding patterns. **B**. The example of p53TAD1 interacting with compound **1050** in steps (**A**). **C**. The predominant binding patterns are shown as ensembles. The binding orientation is visualized as three views with residues as colored centers and the ligand as a ball-and-stick model. The binding strength is quantified using interaction probability of hydrophobic interaction, polar interaction, hydrogen bond, and π-π interaction between residues and the ligand. For hydrogen bonds, the arrow is defined from donor to acceptor.

### Implementation and parameter evaluation of the FuzzAletheia method

The parameters of FuzzAletheia were adjusted using the p53TAD1-**1050** complex trajectories. In step 3, FuzzAletheia uses DBSCAN to cluster point clouds. DBSCAN identifies a point as a core point if at least *minPts* points are within its *ε* radius. It then traverses all reachable points through the neighboring core points to determine which point belongs to the cluster. Decreasing *minPts* can cluster out smaller point clouds (Figure S2A) and is therefore suitable for obtaining short-duration binding patterns. However, it may lead to some point clouds being mistakenly assigned to the same cluster. Given that the predominant binding patterns are wanted, a higher *minPts* threshold was chosen here.

In step 4, as illustrated in Figure 2A, binding pattern extraction uses information on the evolution of conformations over time. For MD simulation, the trajectory is labeled with both the residue-ligand contact and the clustered point clouds to present the evolution of the binding features over time. FuzzAletheia processes the trajectory sequentially in overlapping sliding windows that comprise a specified number of frames along the temporal dimension and extracts binding patterns through the following steps (see details in Materials and Methods): 1) Chronologically identify a core region—a representative window with the most binding features of a binding pattern, including the number of residues binding to the ligand (defined by contact probability), the consistency of the residues’ binding orientation (point clouds) and the binding stability (proportion of point clouds). 2) Align windows in the trajectory to the core region, and 3) assign windows highly similar to the core region to this binding pattern. 4) Remove the assigned windows from the trajectory and repeat the above steps until no new core is found.

**Figure 2.**
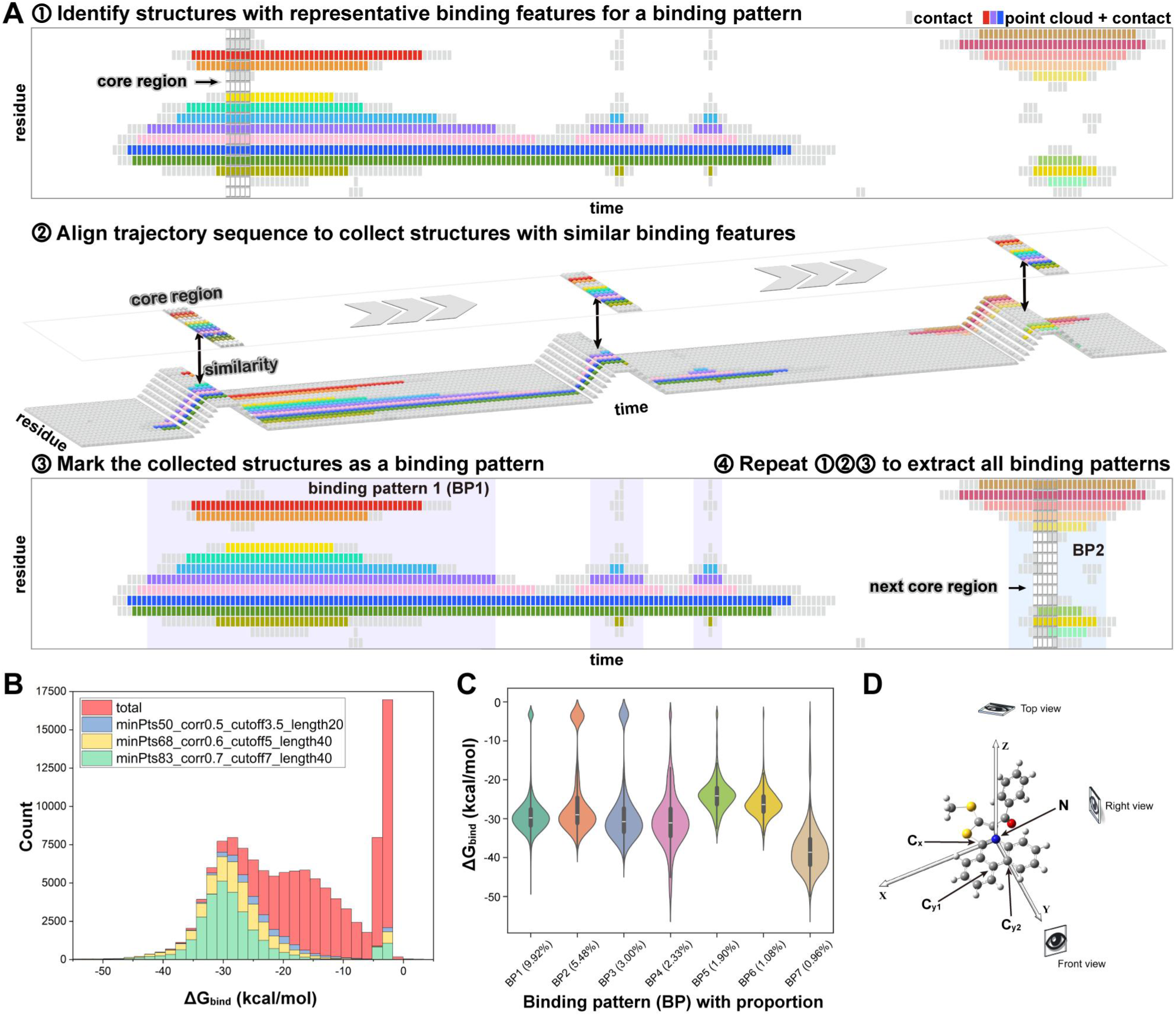
Implementation and parameter evaluation of FuzzAletheia extraction using the p53TAD1-1050 trajectory as an example. **A**. The protocol that FuzzAletheia used to extract binding patterns. The MD trajectory is labeled with point cloud clustering results (colored grid), residue-ligand contact (grey grid), and non-interacting (white grid). Each color represents a cluster of point clouds. **B**. The overall binding free energy (*ΔG_bind_*) distribution of extracted binding patterns shows that FuzzAletheia performs well in extracting low-energy structures. **C**. The binding free energy (*ΔG_bind_*) distributions of the seven p53TAD1-**1050** predominant binding patterns indicate that FuzzAletheia has outstanding clustering ability. **D**. The new coordinate system is defined based on compound **1050** for better visualization of its binding patterns. The N atom was selected as the new origin. The C_x_ was used to define the x-axis. The C_y1_ and C_y2_ were used to define the y-axis.

The extraction process contains four parameters: *L*, the length for defining window size; *P_contact_*, the contact probability cutoff to define whether a residue binds to the ligand in a window; *S_core_*, the cutoff calculated using the number of binding residues for defining a core region; and *S_query_*, the similarity of the queried window to the core region for window assignment. Increasing these parameters represents a stricter definition of the binding pattern. Increasing *L* means that the core region of a binding pattern should persist longer, and increasing *S_core_* means that more binding residues should be involved in a binding pattern, both leading to a reduction in the total number of binding patterns (Figure S2B). Increasing *S_query_* requires a window to more closely resemble the core region to be assigned to the binding pattern, resulting in fewer frames per pattern (Figure S2C) and an increased total number of binding patterns. A larger *P_contact_* represents a stricter requirement for the continuity of the binding in a window. For ensembles with poor temporal continuity due to the sampling method used, such as the trajectories produced by REST2, reducing *P_contact_* can extract more binding patterns (Figure S3).

Next, we evaluated the extraction performance by analyzing the distribution of binding free energies of the extracted binding patterns of the p53TAD1-**1050** system. The extraction process is energy-independent, but as shown in Figure 2B, even the most relaxed extraction conditions accurately extract low-energy structures, indicating that FuzzAletheia captures the nature of IDP-ligand binding. Furthermore, the energy distribution of each binding pattern (Figure 2C and Figure S2D-E) shows a concentrated distribution, validating the rationality of our binding feature-based clustering method. A moderate parameter was selected for the subsequent binding pattern analysis because long-duration patterns are of interest (Figure S2C, black box: *minPts*=83, *S_core_*=7, *L*=40, *S_query_*=0.7, *P_contact_* =0.5).

## Results

### Overall Binding Analysis for the p53TAD1-1050 Complex

Since we attempt to design derivatives by computationally revealing interaction patterns between p53TAD1 and **1050**, careful analysis of the complex MD trajectory is necessary. Figure 3B shows that the maximum residue-ligand contact probability is between **1050** and p53TAD1 residue W23, which is consistent with the previous NMR study (*34*) that the binding of **1050** caused significant chemical shift perturbations in p53TAD1 W23, confirming that our simulations align well with the experimental data. Thus, we further analyzed the p53TAD1-**1050** binding features based on these trajectories.

**Figure 3.**
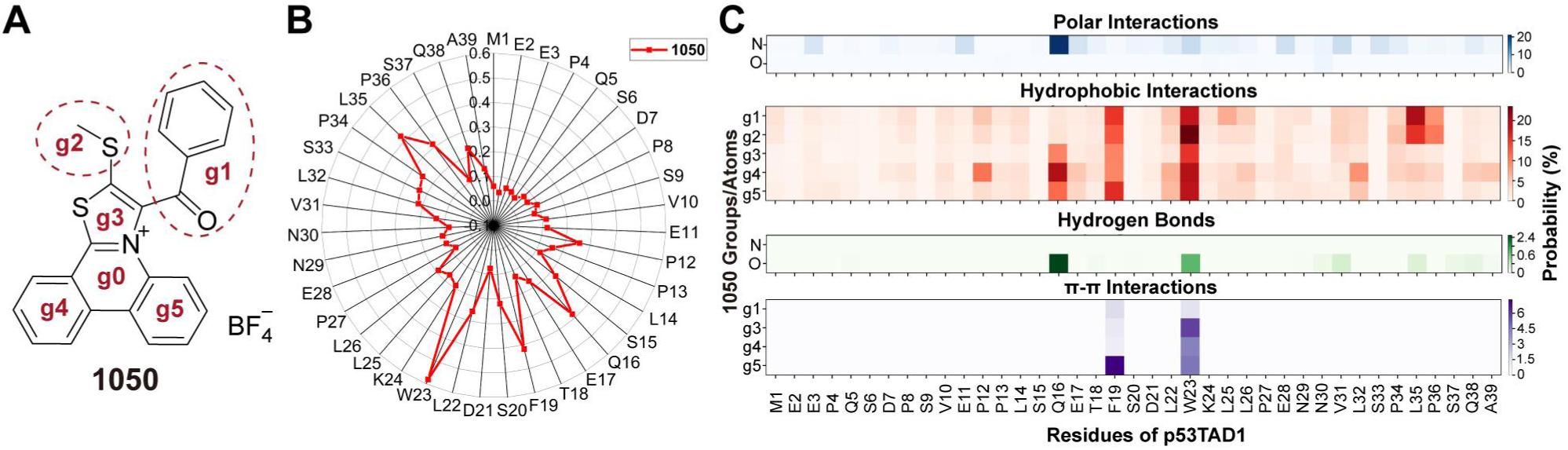
Overall analysis of the p53TAD1-1050 complex MD trajectory. **A**. Definition of groups in compound **1050**. **B**. Contact probabilities between residues and compound **1050**. **C**. Interaction probabilities between p53TAD1 residues and **1050** groups. The maximum probabilities of polar, hydrophobic, hydrogen bond, and π-π interactions are 21.1% (Q16, N), 23.6% (W23, g2), 2.6% (Q16, O), and 7.3% (F19, g5), respectively.

Quantification of individual chemical group contributions to binding is critical for elucidating IDP-small molecule interaction mechanisms and enabling rational ligand optimization. Therefore, we divided **1050** into six groups (Figure 3A) and analyzed their four types of non-bonded interaction probabilities with residues, including polar, hydrophobic, hydrogen bond, and π-π interactions. Before analyzing predominant binding patterns, we conducted an overall analysis of the p53TAD1-**1050** interaction. As shown in Figure 3C, **1050** predominantly binds to p53TAD1 via hydrophobic interactions with F19, W23, and P34–P36, as well as polar interactions with Q16. For hydrophobic interaction, groups g1–g5 all exhibit high probability with W23; groups g1–g2 are preferred by P34–P36 and group g5 is preferred by F19. The considerable polar interactions between the N atom in g0/g3 and Q16 play a key role in binding. The π-π interactions basically occur between g3–g5 and F19, W23, in which g5 is strongly preferred by F19. The carbonyl O atom in g1 forms a few hydrogen bonds with Q16 and W23. These interactions collectively maintain the fuzzy p53TAD1-**1050** complex, though the detailed binding features need further revealing.

### Dynamic Binding Pattern Analysis of the p53TAD1-1050 Complex Using FuzzAletheia

We used FuzzAletheia to extract predominant binding patterns of the p53TAD1-**1050** complex for ligand optimization. All seven predominant binding patterns extracted using the moderate parameter are presented in Figure 4 and Figure S4, in which the coordinate system of point cloud plots was defined as in Figure 2D. Together, the seven patterns account for 24.67% of the total trajectory (Figure 4A). Their binding energies range from -20 to -40 kcal/mol (Figure 2C). BP1 is the most predominant binding pattern (BP), accounting for 9.92%. In BP1 (Figure 4C), Q16 anchors **1050** within the hydrophobic pocket formed by residues including P12, F19, W23, and L35–A39 through polar interaction with the N atom in g0 and 19% hydrogen bonds with the O atom in g1. The proper orientation allows g5 to form a 64.71% π-π interaction with F19. Most hydrophobic interactions involve the g4-5 groups, and g1 only interacts with W23 and L35. BP7 is the lowest-energy binding pattern, accounting for 0.96%. In BP7, E28 anchors **1050** with polar interactions, and other residues wrap **1050** well with hydrophobic interactions, leaving only groups g1 and g2 solvent accessible. In all the binding patterns, p53TAD1 anchors **1050** in a hydrophobic pocket through polar interactions by a residue above or below the g0 ring. From BP2 to BP6, the anchor residues are Q16, V31, S33, Q16 and K24/L25, respectively. The ligand groups participating in hydrophobic interactions are as follows: in BP2, groups g1 and g2; in BP3, groups g1 and g5; in BP4, groups g1 and g5; in BP5, group g1; and in BP6, groups g1 and g2 (Table S1). For hydrogen bonds, in addition to those formed by the anchor residues (Q16 in BP1 and V31 in BP3), W23 is also accessible for g1 when it is near g5 (BP4). The combined analysis using point cloud plots and interaction probability plots reveals how **1050** groups contribute to forming the existing binding patterns.

**Figure 4.**
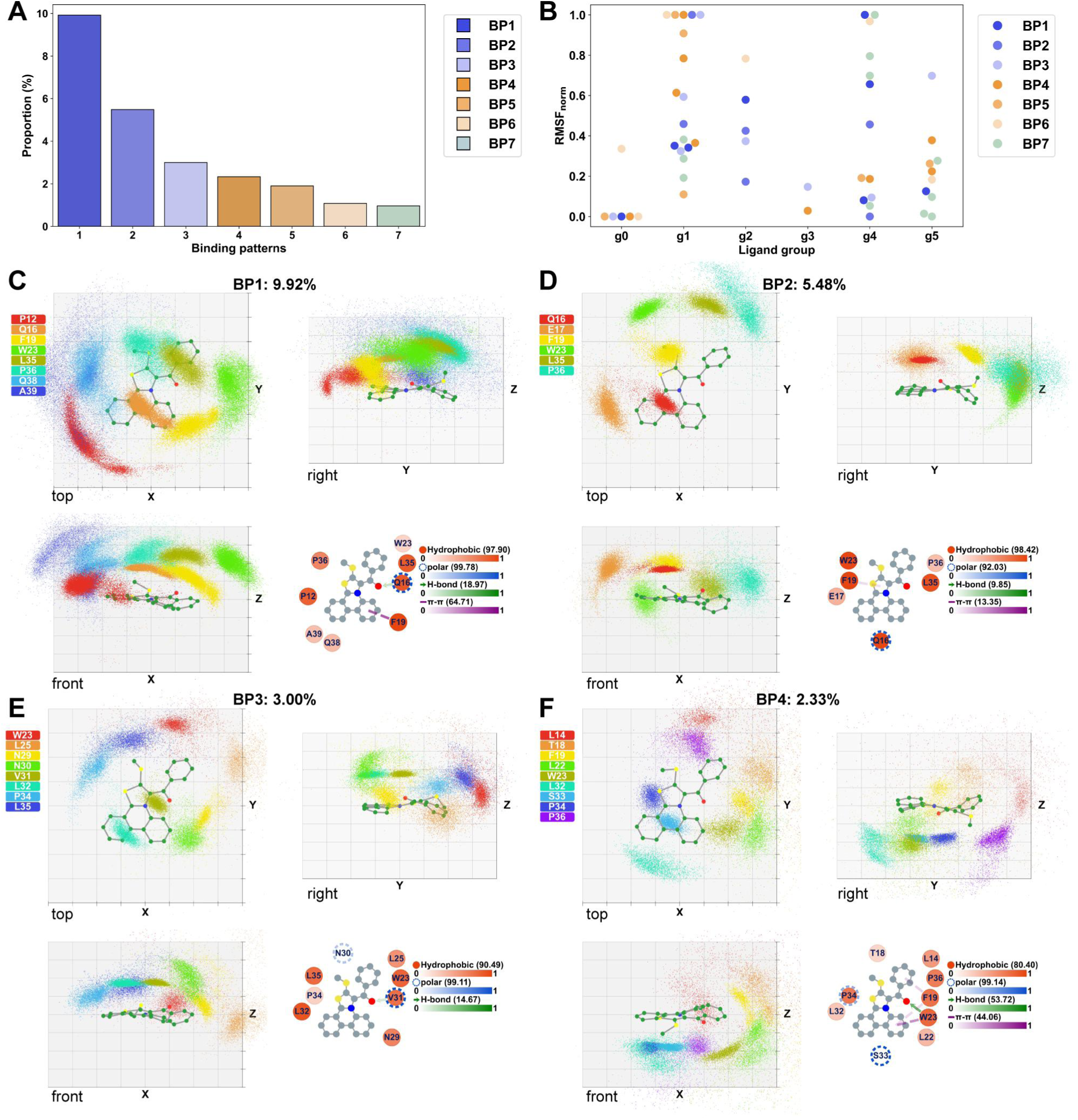
Predominant binding pattern analysis of the p53TAD1-1050 complex. **A**. The proportion of predominant binding patterns in the trajectory. The proportion of each pattern is calculated by dividing the number of frames in the binding pattern by the total number of frames. **B**. The normalized RMSF of distances between the six groups of **1050** and interacting residues. High RMSF_norm_ values indicate unstable interactions and vice versa. **C**-**F**. The orientation and interaction of each binding pattern are visualized using FuzzAletheia. For the orientation plot, each 3D binding pattern is presented through three views, where the compound is shown in ball-and-stick and residues are shown in colored geometric centers. The grid in the background has a spacing length of 2.5 Å. For the interaction plot, the hydrophobic, polar, π-π, and hydrogen bond interactions are shown as red circles, blue segmented circles, purple dashed lines, and green arrows (pointing from donor to acceptor), respectively. The number of each interaction refers to its maximum probability.

The binding patterns formed by IDPs and ligands are highly dynamic, with residues in constant motion, leading to unstable interactions. Stabilizing these unstable interactions may improve the ligand binding ability. Specifically, comparative analysis of residues’ binding stability within binding patterns can discriminate between ligand groups indispensable for binding and warranting modification. We propose a root-mean-square distance fluctuation (RMSF)-based metric, defined by the distance between residues and ligand groups of maximal interaction probability, to evaluate the binding stability of residue-ligand group pairs. As shown in Figure 4B and Table S1, group g0 is involved in the largest number of stable interactions, while group g1 is largely involved in unstable interactions. The residues forming polar interactions with the N atom of g0 are essentially the most stable ones in all binding patterns, indicating their crucial role in the formation of these binding patterns: on one hand, the N and O atoms of compound **1050** can form a synergistic polar interaction and hydrogen bond with the amide group (Figure 4C, E and Figure S5); on the other hand, the formation of these polar interactions may be attributed to the space limitation to the residues caused by the bulky ring (g3–g5), implying indirect contributions of groups g4–g5 to the binding. Groups g1–g2 mainly play their roles via hydrophobic interactions. As shown in Figure 4, the residues interacting with g2 are more stable due to their proximity to the hydrophobic core formed by p53TAD1 and **1050**. In contrast, the residues interacting with g1 are less stable and have low interaction probabilities. Their interactions are less stable as they are distanced from the core, and the interaction with g1 does not constrain them sufficiently, highlighting that group g1 has the potential for further optimization.

As analyzed above, g1 is involved in several hydrophobic interactions and minimal π-π interactions through the phenyl ring, suggesting that its substitution may stabilize the existing interactions. The hydrogen bonds formed by the carbonyl in group g1 are important parts of the anchor interactions that we would like to preserve, so the benzoyl ortho-substitutions are excluded for their possible interference. From the binding orientation in BPs 1-7 (Figure 4C-F and Figure S4), both the meta- and para-positions of the benzoyl group can accommodate substituent modifications. Thus, we chose the para-position in g1 to design derivatives for experimental testing.

Exploiting the stability of residue-ligand interactions simplifies the complicated task of analyzing a large number of heterogeneous IDP-ligand binding patterns. We determine the modification sites according to the following rules: 1) Identify the ligand groups to be retained, that is, the groups engaged in anchor interactions, through orientation, interaction types, and binding stability analysis. 2) Recognize the ligand groups involved in the unstable interactions through the binding stability analysis. 3) Pinpoint modification sites that are involved in unstable interactions and design synthetically feasible compounds that can stabilize the unstable interactions.

### Optimization of 1050 Guided by FuzzAletheia Analysis

As there are hydrophobic and polar residues near the para-position of the g1 benzoyl group, we designed and synthesized six derivatives (Figure 5G) with both polar and hydrophobic substituents to explore the SAR. The synthetic route of the derivatives is shown in Figure S6. We first confirmed their binding to p53TAD1 using surface plasmon resonance (SPR) assay (Figure 5A, G and Figure S7). The p53TAD1 protein was immobilized on a CM5 chip through amine coupling, and the compounds flowed through the chip as a mobile phase to measure their binding ability. All modifications led to an increase in binding affinity, especially the bulky substituents. These results validate the importance of the modification site selected by FuzzAletheia. We then checked whether these compounds can break p53-MDM2 binding using the AlphaScreen assay. In the AlphaScreen assay, the interacting p53 and MDM2 bring the donor and acceptor close through their respective tags, enabling the quantification of the p53-MDM2 binding based on the emission intensity of the acceptor. Figure 5B demonstrated that derivatives with bulky hydrophobic substituents exhibited enhanced inhibitory capacity, while those with polar substituents displayed reduced inhibition. This observation aligns with the binding mechanism of p53TAD1 to MDM2, in which residues 19–26 in p53TAD1 insert into the hydrophobic cleft of MDM2. The hydrophobic substituents likely strengthen interactions of the ligands to the key hydrophobic residues (F19 and W23), thereby enhancing p53-MDM2 inhibition. In contrast, polar substituents may induce p53TAD1 conformations that weakly constrain residues 19–26, leading to decreased inhibition.

**Figure 5.**
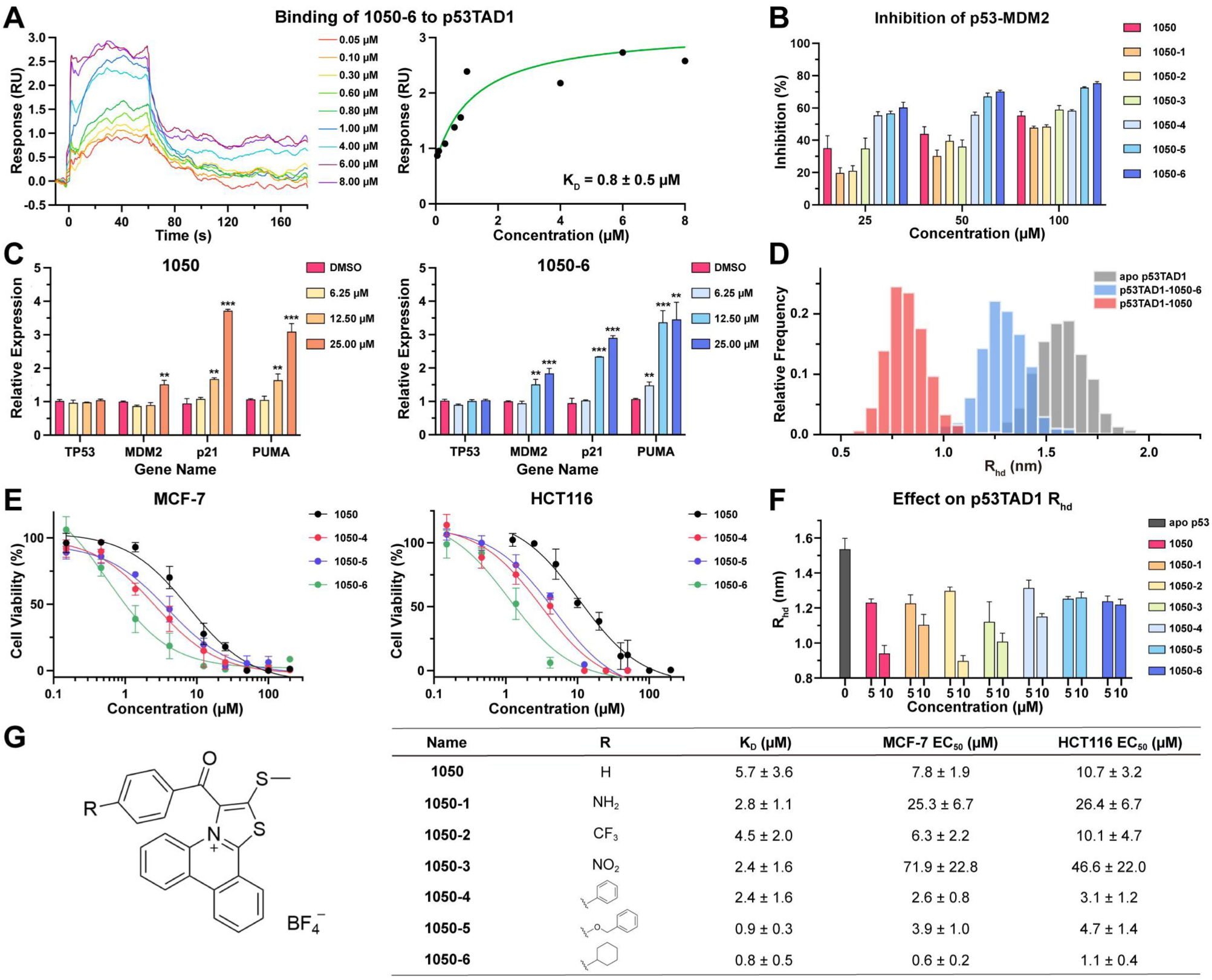
The activity of 1050 derivatives. **A**. SPR sensorgrams and binding affinity of **1050-6** to p53TAD1 on CM5 chip. **B**. AlphaScreen assay for p53-MDM2 interaction inhibition by **1050** and derivatives. **C**. Quantitative PCR analysis of the expression levels of typical p53 target genes. Data presented as mean ± SD (n = 3); \*\**P* < 0.01, \*\*\**P* < 0.001; unpaired t-test. **D**. *R_hd_* from FCS of p53TAD1 in apo, **1050**-bound, and **1050-6**-bound states. The compound concentration used in the FCS assay was 10 μM. **E**. Antiproliferative activity of **1050**-series compounds in MCF-7 and HCT116 cell lines. **F**. FCS analysis of p53TAD1 *R_hd_* changes upon compound binding. **G**. Binding affinity and antiproliferative activity of **1050**-series compounds.

We selected **1050-6** as an example to compare how the binding of compounds sharing a common scaffold influences the p53TAD1 conformational ensemble (Figure 5D) and the similarities and differences in their binding patterns (next section). We used fluorescence correlation spectroscopy (FCS) to infer the hydration radius (*R_hd_*) of the labeled p53TAD1 by examining its diffusion coefficient. As shown in Figure 5D, FCS results demonstrated that after adding **1050** or **1050-6**, the p53TAD1 *R_hd_* becomes smaller, indicating that both ligands induce p53TAD1 to form more compact conformations. Compared to **1050**, **1050-6** resulted in a smaller shift, probably because the bulky substituent extends the p53-bound state conformations. The *R_g_* distributions calculated from MD simulations also indicated that the binding of both ligands leads to a more compact p53TAD1 conformational ensemble (Figure S8). This consistency again confirms the reliability of our simulations. We further measured the effects of all the **1050**-series compounds on the p53TAD1 conformational ensemble (Figure 5F). Although all ligands can induce p53TAD1 to become more compact, the shrinking extents are different, suggesting that while sharing common binding features, they also have unique ones.

To explore the cellular activity of these ligands, we investigated their effect on the expression of p53-regulated genes and antiproliferative activity in MCF-7 and HCT116 cell lines. Figure 5C shows that the p53 target genes *p21*, *puma*, and *MDM2* were transcriptionally activated upon treatment with both **1050** and **1050-6** in a dose-dependent manner without influencing the expression of the *p53* gene. **1050-6** exhibited superior activity to **1050**, as it achieved activation at a lower concentration. As shown in Figure 5E, G and Figure S9, the derivatives with bulky hydrophobic substituents (**1050-4**, **1050-5**, **1050-6**) were more effective in reducing cancer cell viability than **1050**. The derivatives with polar substituents showed reduced potency. Considering that disrupting the p53-MDM2 interaction results in the activation of p53, which promotes cancer cell death, the SAR observed in binding affinity, p53-MDM2 interaction inhibition, p53-regulated genes activation, and cell growth inhibition align consistently. These findings underscore the value of the binding patterns identified using FuzzAletheia in guiding ligand optimization.

As analyzed above, the interactions formed by g1 are basically hydrophobic; thus, the introduction of bulky hydrophobic substituents may stabilize existing hydrophobic interactions and enhance binding strength between p53TAD1 and ligands. Together, the activity of the **1050**-series compounds observed in the experiment is consistent with the results of the predominant binding pattern analysis.

### FuzzAletheia Reveals that 1050-6 Enhances Its Binding Affinity for p53TAD1 by Retaining the Existing Binding Pattern of 1050 and Inducing New Interactions

To check whether our design principles work, we ran 1.2 µs MD simulations of the p53TAD1-**1050-6** complex and conducted overall binding analysis and binding patterns analysis. The overall binding analysis (Figure 6B) showed that **1050-6** largely retains its preference for Q16, F19, and W23, shifts and enhances its preference from P34–P36 to N30–L35, and shows new preferences for E2–Q5 and P8–V10. Comparing the interaction heatmap of **1050-6** (Figure 6C) to that of **1050** (Figure 3C), introducing group g6 makes polar interactions and hydrogen bonds shift to the main chain of V31. The maximum probability of polar interactions shows a significant increase, while that of hydrogen bonds jumps from 2.6% to 11.0%, suggesting the emergence of more stable binding patterns. For hydrophobic interactions, group g6 replaces g3–g5 to interact with F19 and W23.

**Figure 6.**
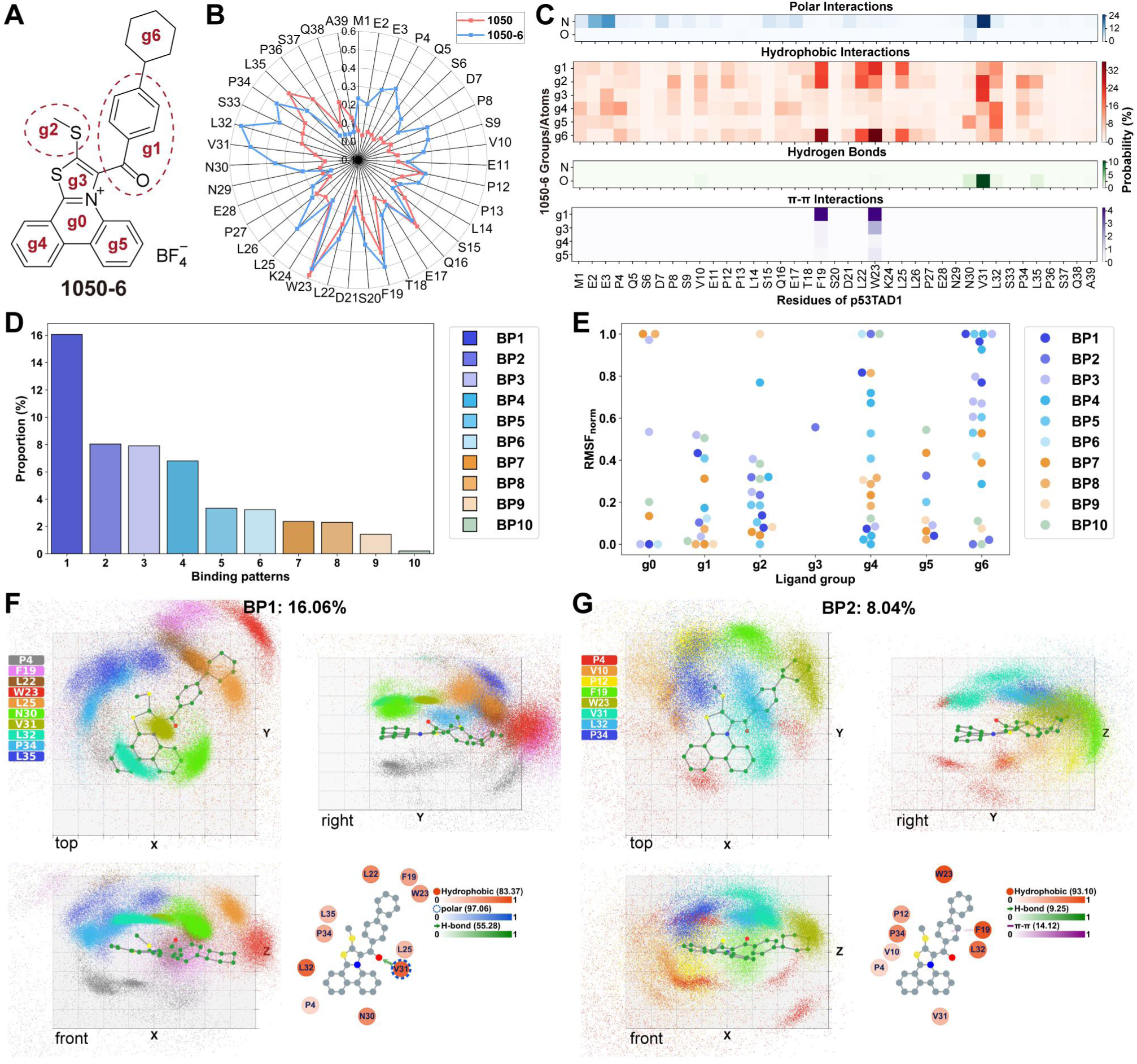
Analysis of the p53TAD1-1050-6 complex MD trajectory. **A**. Definition of groups in compound **1050-6**. **B**. The residue contact probability of compounds **1050** and **1050-6**. **C**. The interaction heatmap between p53TAD1 residues and **1050-6** groups. The maximum probabilities of polar, hydrophobic, hydrogen bond, and π-π interactions are 25.1% (V31, N), 35.7% (W23, g6), 11.0% (V31, O), and 4.3% (W23, g1), respectively. **D**. The proportion of predominant binding patterns in the trajectory. **E**. The normalized RMSF of distances between the seven groups of **1050-6** and interacting residues. High RMSF_norm_ values indicate unstable interactions and vice versa. **F**-**G**. Binding patterns of the p53TAD1-**1050-6** complex are visualized using FuzzAletheia. For the orientation plot, each 3D binding pattern is presented through three views, where the compound is shown in ball-and-stick and residues are shown in colored geometric centers. The grid in the background has a spacing length of 2.5 Å. For the interaction plot, the hydrophobic, polar, π-π, and hydrogen bond interactions are shown as red circles, blue segmented circles, purple dashed lines, and green arrows (pointing from donor to acceptor), respectively. The number of each interaction refers to its maximum probability.

From the interaction heatmap analysis, it seems that the introduction of g6 significantly changed the binding patterns between p53TAD1 and ligands. However, when we extracted the p53TAD1-**1050-6** binding patterns (Figure 6F-G and Figure S10-11) using FuzzAletheia and compared them to those of the p53TAD1-**1050** complex, it is interesting to find that the p53TAD1-**1050** pattern 3 becomes the largest pattern 1 of the p53TAD1-**1050-6** complex (Figure S12A). The slight difference lies in the shift of the position of W23 and the engagement of P4 below g3–g5. This difference is probably because introducing a bulky group like g6 forces p53TAD1 to use more N-terminal residues to wrap the ligand when the C-terminal residues remain unchanged. More importantly, due to a stricter constraint of F19, L22, and W23, the N-terminal residue P4 is more easily accessible to group g4, which even forms a stable interaction, thus making the pattern more advantageous.

The scenario where the **1050-6** scaffold preserves the original interactions of **1050**, while the newly introduced group forms additional interactions that stabilize the binding pattern, is similar to selecting an appropriate binding pattern from that induced by the lead compound. This resembles the conformational selection model, but instead of starting from the unbound conformation, it begins with the bound conformation of the lead compound. This indicates that even for IDP ligands, derivatives can be designed to stabilize the established binding patterns of the lead compound to improve activity.

FuzzAletheia extracted a total of ten p53TAD1-**1050-6** binding patterns, accounting for 51.69% of all the conformations in the ensemble (Figure 6D), which is significantly higher than that of p53TAD1-**1050** (24.67%), and their binding energies range from -30 to -50 kcal/mol (Figure S13). A new type of pattern (Figure 6G, BP2) is identified, which is dominated by the hydrophobic interactions of F19, W23, and L32. The third largest binding pattern (Figure S10, BP3) is maintained by V31 and L32, which is accompanied by the polar interactions of N-terminal residues E2 and D7 from the other side of the g3–g5 ring. The binding pattern where Q16 serves as an anchor remains (Figure S10, BP4), though it differs from the p53TAD1-**1050** BP1, 2, or 5 but resembles p53TAD1-**1050** BP7 (Figure S12B). Other binding patterns distinct from patterns of the p53TAD1-**1050** complex are shown in Figure S10-11. Given the high flexibility of the p53TAD1 structure, it was not surprising that **1050-6** induces new p53TAD1 bound states. Compared to p53TAD1-**1050** binding patterns, the patterns of the p53TAD1-**1050-6** complex generally have lower energy, longer duration, and more residues involved, indicating a stronger binding ability of **1050-6** for p53TAD1, consistent with experimental results. The binding stability analysis (Figure 6E) of **1050-6** groups demonstrated that the introduction of g6 does stabilize the interactions g1 engages. It is worth mentioning that the introduction of g6 makes g2–g5 groups participate in more interactions and has little influence on the distribution of their interaction stability but significantly introduces additional weak interactions for the g0 group. In BPs 3, 7, and 8, the newly introduced weak interactions arise from additional polar interactions with residues other than the anchor residues (Table S2). These binding patterns provide a prototype for the more stable binding patterns that can be formed by p53TAD1 and the **1050** scaffold.

### General applicability of FuzzAletheia

To examine the general applicability of FuzzAletheia, we analyzed the binding patterns of disordered AR-Tau5 and EPI-series ligands, including **EPI-002** (*38*) and **iodoEPI-002** (*39*), based on the freely available trajectories that were generated using REST2 (*17*). The impact of the poor continuity of the trajectory caused by replica exchange on binding pattern extraction was reduced by lowering *P_contact_* (Figure S3). Other parameters used for extraction are listed in Table S3. Six predominant binding patterns that totally account for 4.64% were extracted for the AR-Tau5-**EPI-002** system (Figure S14-15). The binding stability analysis revealed that group g2 has the largest number of unstable interactions and is therefore the most suitable place for modification (Figure S14D). Another candidate is the other phenyl ring, group g4. If the proportion of these interactions is taken into account, the weighted binding stability analysis also shows that group g2 is worth modifying (Figure S14E). Next, we analyzed the binding patterns formed by AR-Tau5 and **iodoEPI-002**, an analog with iodination on group g2. In previous studies (*17, 39*), **iodoEPI-002** has shown a 10-fold increase in experimental activity and an enhanced binding affinity in MD simulation (Figure S16). Thirteen predominant binding patterns that totally account for 28.69% of the trajectory were extracted (Figure S17-20), which is significantly higher than that of EPI-002 (4.64%). Figure S17D shows that the interactions of group g2 are stabilized after substitution, and group g5 of iodoEPI-002 is worth further modification. In BPs 1, 4, 5, and 9, AR-Tau5 embraced iodoEPI-002 in a crescent shape from the side of the iodine substituent, forming a stable hydrophobic core with group g2, accompanied by modest hydrogen bonds and polar interactions with group g1 or g5. These novel BPs explain that the iodine substituent enhances the binding ability by providing a center to form more stable hydrophobic cores. The analysis results of AR-Tau5 and EPI-series ligands are in line with the IDP ligand modification site selection strategy proposed here, proving the universality of FuzzAletheia.

## Discussion

While an increasing number of ligands that bind directly to IDPs have been discovered, their optimization for progressing to clinical use remains challenging due to the lack of effective optimization methods. In this study, we have developed FuzzAletheia, a novel approach to deciphering binding patterns between IDPs and ligands that can guide ligand optimization. By integrating binding orientation, interaction types, and binding stability analysis, we revealed binding patterns of the p53TAD1-**1050** complex and employed this knowledge for rational optimization that led to the identification of three more potent compounds. This study develops a method to define, extract, visualize, and quantify dynamic binding patterns of fuzzy IDP-ligand complexes to guide ligand optimization, providing a framework for structure-based ligand optimization targeting IDPs.

FuzzAletheia presents residues as geometric centers, balancing binding dynamics and easy visualization. By providing orientation, interaction, and stability insights, FuzzAletheia can be used to pinpoint the contribution of each group in the ligand and guide the selection of important modification sites for optimization. Additionally, FuzzAletheia also provides an excellent clustering method for protein conformations with inter-residue dynamic differences. For example, with the alignment based on a key residue prone to intramolecular interactions, FuzzAletheia can cluster apo IDP conformations based on the interactions of that key residue.

While FuzzAletheia offers valuable insights, it is not without limitations. Currently, it remains difficult to quantify the binding strength of new compounds without the support of extensive MD simulations. An alternative strategy involves using FuzzAletheia for clustering, followed by the “multi-conformational-affinity” strategy (*34*) to rank derivatives, although this process might overlook binding dynamics and is probably time-consuming. Therefore, direct prediction of compound affinities via point cloud binding patterns needs to be further investigated. Another limitation may arise when dealing with highly flexible compounds, which could disperse the point clouds and reduce both binding pattern extraction accuracy and visualization clarity. In such cases, clustering compound conformations and then analyzing the binding patterns of each cluster could be effective.

This study is based on two basic hypotheses: first, predominant binding patterns exist within IDP-ligand complexes, and second, compounds with a common scaffold share similar binding patterns. Therefore, the extracted binding patterns can assist structure-based lead optimization. For the first assumption, many studies have demonstrated the presence of key residues for binding and specificity (*40-42*). These residues are likely to form predominant binding patterns. Our analysis of **1050** and **1050-6** binding patterns also provides evidence for this hypothesis. As for the second assumption, our results show that **1050-6** can retain binding patterns from the patterns of the common scaffold. An NMR study also indicated that compound Givinostat and its derivative GSD-16-24 can bind α-synuclein in a similar manner (*42*).

We found that different residues can form similar binding patterns (such as BP2 and BP3 of **1050**) and one residue can be involved in different binding patterns along the trajectory. Notably, most p53TAD1-**1050**/**1050-6** binding patterns typically involve the two key residues (F19, W23) that overlap across different combinations of binding residues, which is probably crucial to ensuring the dynamics of p53TAD1. When different residues can form similar binding patterns, their energy levels are often closely matched, allowing for perturbations that prompt transitions between these patterns. When a residue combination can form multiple binding patterns of similar energy with a ligand, the IDP conformation is not fixed but keeps transitioning between these patterns. Considering the scenario where two binding patterns with similar energy do not share common residues: once a ligand has formed a binding pattern, it is challenging for the residues from the other pattern to compete. Conversely, if residues overlap between two binding patterns, proximity allows these shared residues to sway between patterns, effectively creating a dynamic competition for the ligand. This competition of binding patterns resembles Frustration (*43*), where IDP-protein complexes never collapse to single conformations due to competition between suboptimal interactions during binding.

We also observed some less frequent binding patterns (those extracted using smaller *minPts*), which encompass a diverse array of residue combinations and typically include key residues. It is possible that some of these patterns may act as transition states, facilitating the transition of ligands between more stable binding patterns. For example, even when two predominant binding patterns do not overlap in residues, their overlap with a transition state allows interconversion via the transition state. These less frequent binding patterns could account for the extensive influence of ligands on residues as detected by NMR. Such a mechanism might also shed light on the promiscuity observed in some IDP ligands, where many transition binding patterns sustain the binding.

The binding patterns described here are derived from residue-level point clouds, which suggests that even when a point cloud is formed, the residue atoms may remain dynamic, accompanied by the fluctuation of interactions. Therefore, when the cooperative interactions within a pattern have not been formed and are squeezed out by competition from another binding pattern, it may be attributed to dynamic shuttling (*18*). Based on the binding patterns represented by point clouds, the “Fuzziness” of IDP-ligand binding can be considered as a result of matching ligand pharmacophore and residue combinations with specific orientations, which leads to uniform and diverse binding patterns. We name this mechanism “pattern coupling”. Similar binding patterns of different residue combinations, different predominant binding patterns of the same residues, and transition states generated by pattern coupling together lead to fuzzy IDP-ligand complexes. This binding mechanism ensures that IDP is not fixed in a single conformation unless a binding pattern with the ligand becomes optimal.

## Acknowledgments

The authors thank Dr. Beiming Cheng, Dr. Limin Chen, Dr. Yibo Li and Dr. Shiwei Wang for helpful discussions, Liying Wang, Yujie Yang, Junxi Mu and Zhixian Huang for manuscript improvement, Dr. Hantian You and Dr. Weijie Bian for experimental help and Zhaoyang Li for coding skills.

## Funding

National Natural Science Foundation of China (22237002)

National Natural Science Foundation of China (T2321001)

National Natural Science Foundation of China (22307002)

The CAMS Innovation Fund for Medical Sciences (2021-I2M-5-014)

## Notes

### Competing Interest Statement

The authors have declared no competing interest.

