## Supplemental Figures S1-49 and Tables S1-3 for "FuzzAletheia: Mapping dynamic interaction landscapes of intrinsically disordered protein-ligand complexes to accelerate ligand optimization"

Hanping Wang *et al.*

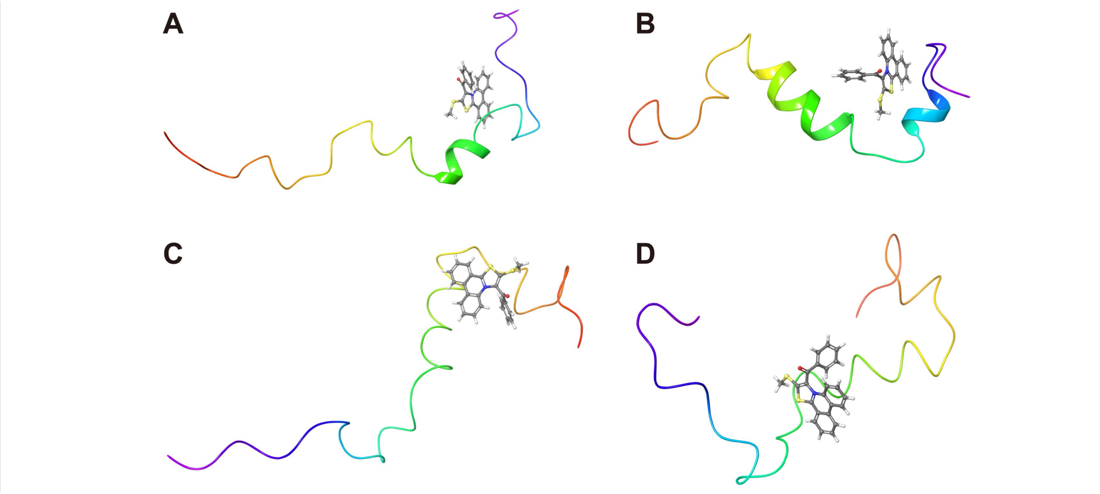


**Figure S1**. Initial structures for p53TAD1-1050 MD simulations, where the p53TAD1 structures were also used to run apo MD simulations. p53TAD1 is shown in the rainbow-colored cartoon with the N-terminus in blue and the C-terminus in red. Compound 1050 is shown in ball-and-stick.

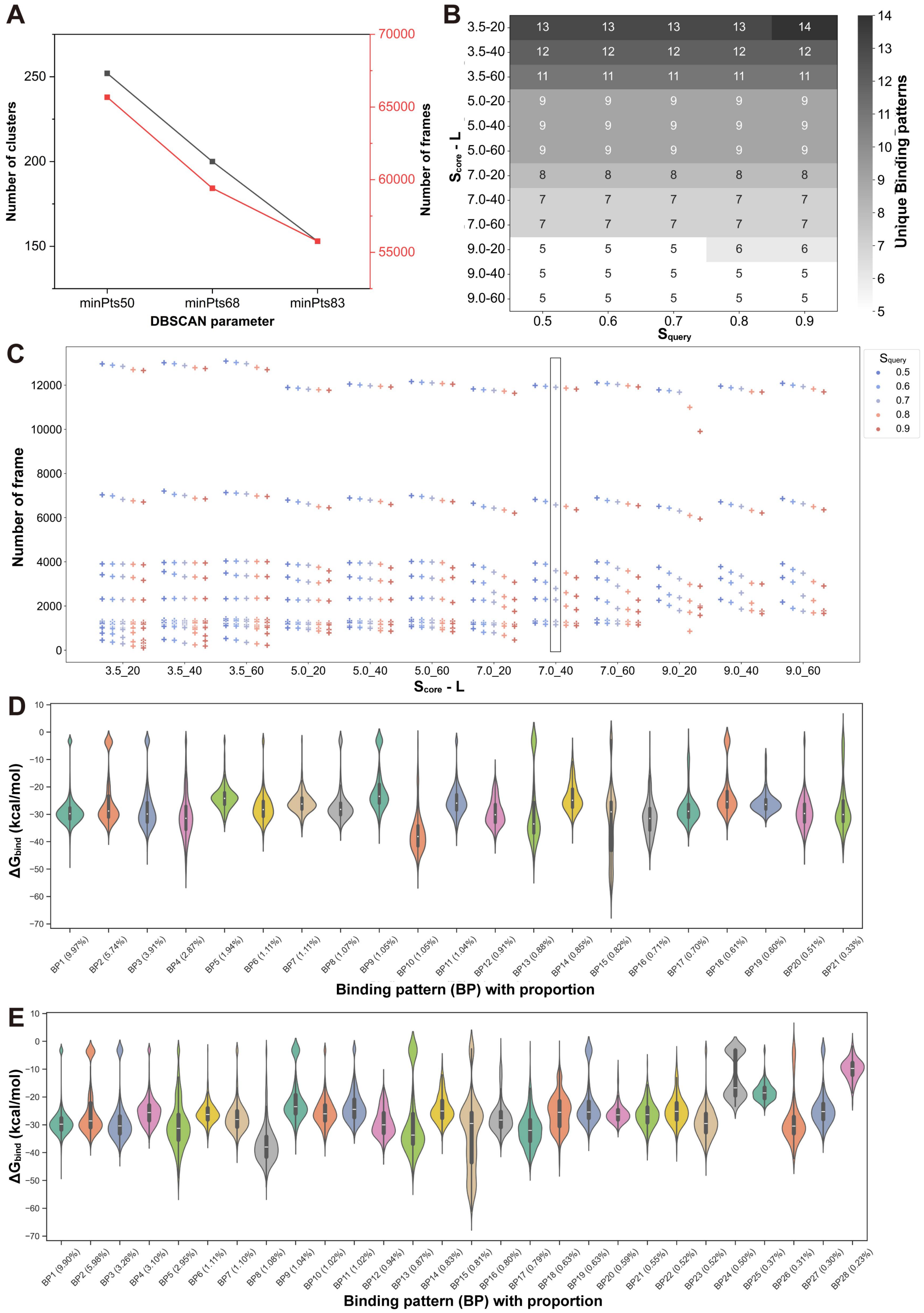


**Figure S2**. Evaluation of the influence of parameters on the performance of extraction. (A) The influence of *minPts* of DBSCAN on the total number of point clouds and extracted structures. (B) The influence of extraction parameters on the total number of extracted binding patterns. $S_{core}$ and $L$ determine the lower bound of the core region selection. $S_{query}$ determines the lower bound of the region least like the binding pattern. (C) Increasing $S_{query}$ decreases the total number of frames per pattern. The point clouds were clustered using *minPts*=83. The adopted parameters are circled in the black box. (D) The binding free energy ($\Delta G_{bind}$) distributions of the p53TAD1-1050 binding patterns extracted using the following parameters: *minPts*=68, $S_{core}$=5, $L$=40, $S_{query}$=0.6. (E) The binding free energy ($\Delta G_{bind}$) distributions of the p53TAD1-1050 binding patterns extracted using the following parameters: $minPts$=50, $S_{core}$=3.5, $L$=20, $S_{query}$=0.5.

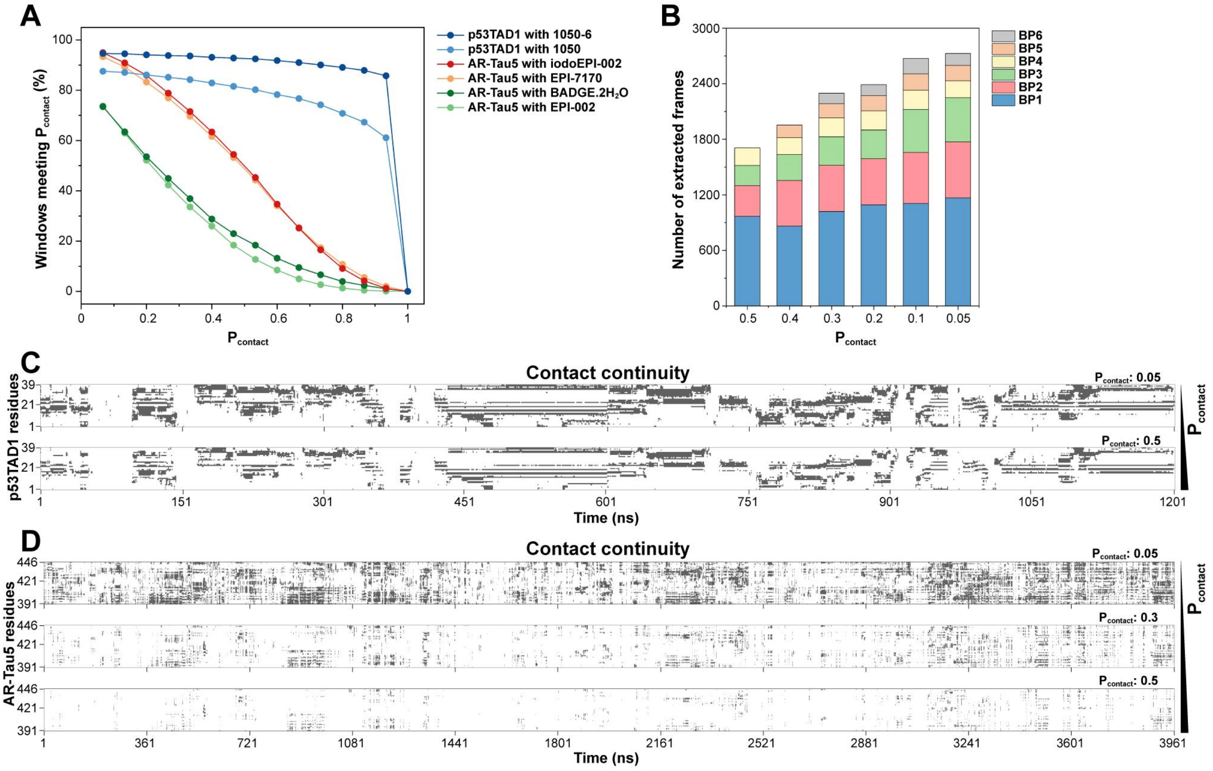


**Figure S3**. Evaluation of the influence of $P_{contact}$ on the performance of FuzzAletheia extraction. (A) The proportion of windows that have at least four residues binding to the ligand with a contact probability > $P_{contact}$ in trajectories produced by classical MD simulation (p53TAD1 systems) and REST2 (AR-Tau5 systems). Only these windows possibly contain predominant binding patterns. (B) The number of frames of the predominant binding patterns formed by AR-Tau5 and EPI-002 that are extracted using varying $P_{contact}$. (C)-(D) The temporal continuity of contacts formed by the residues and the ligand. Residues with a contact probability greater than the cutoff value with the ligand within the specified time period (0.8 ns for p53TAD1-1050 and 1.2 ns for AR-Tau5-EPI-002) are marked black; otherwise, they are marked white. The p53TAD1-1050 system (C), simulated using classical MD, shows good continuity. The system of AR-Tau5 and EPI-002 (D), simulated using REST2, exhibits poor continuity where the contact between the residue and the ligand is often interrupted.

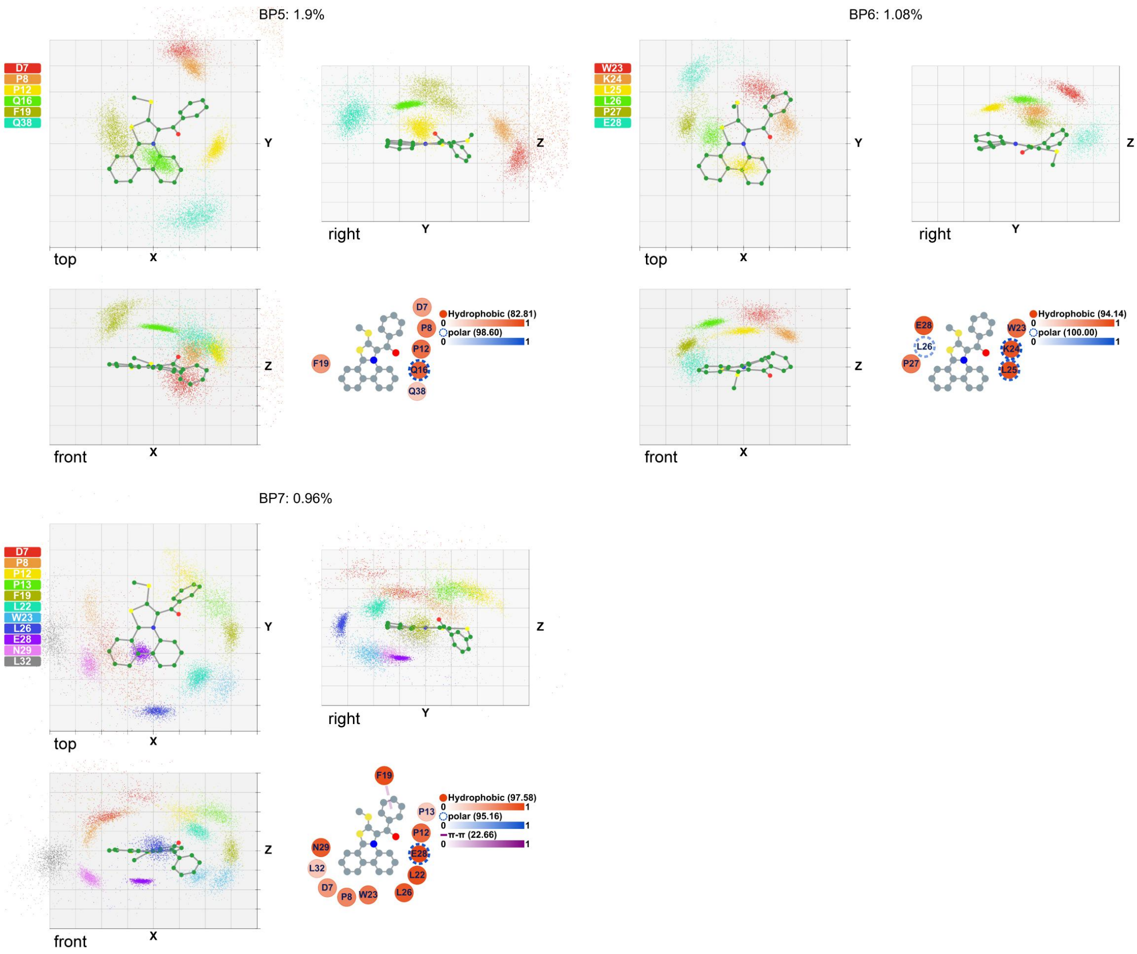


**Figure S4**. Binding patterns 5-7 of the p53TAD1-1050 complex. For the orientation plot, each 3D binding pattern is presented through three views, where the compound is shown in ball-and-stick and residues are shown in colored geometric centers. The grid in the background has a spacing length of 2.5 Å. For the interaction plot, the hydrophobic, polar, π-π, and hydrogen bond interactions are shown as red circles, blue segmented circles, purple dashed lines, and green arrows (pointing from donor to acceptor), respectively. The number of each interaction refers to its maximum probability.

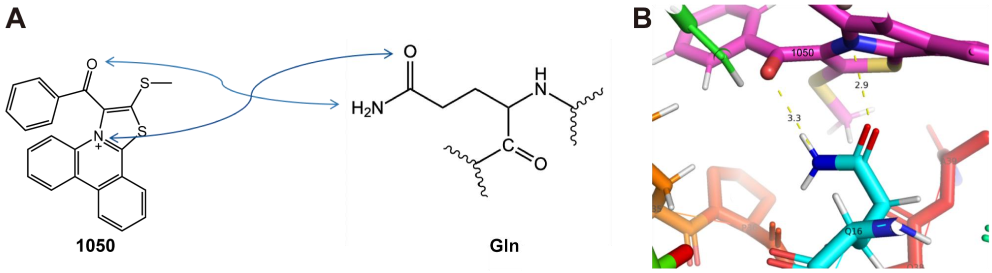


**Figure S5**. The synergistic polar interaction and hydrogen bond formed by the N and O atoms of compound 1050 with the amide group. (A) The illustration of the synergistic interaction. (B) A representative conformation of the synergistic interaction.

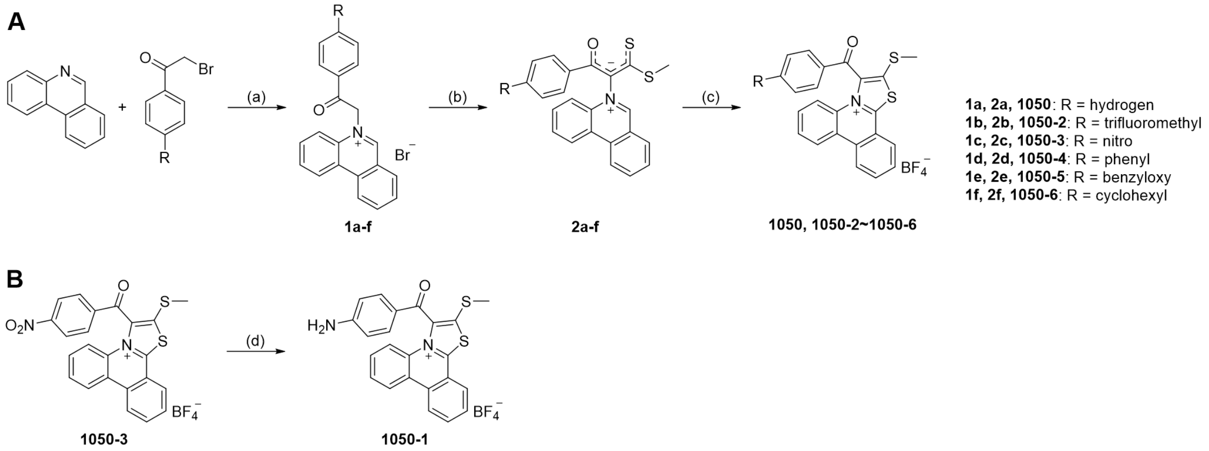


**Figure S6**. Synthetic route of 1050-series compounds. (A) Reagents and conditions: (a) MeCN, refluxed, 48 h; (b) $CS_2$, MeI, 43% $K_2CO_3$, 20-24 h; (c) $HBF_4$, MeOH, 80-83 °C, 72-96 h. (B) Reagents and conditions: (d) Fe, $NH_4Cl$, 2.5 h.

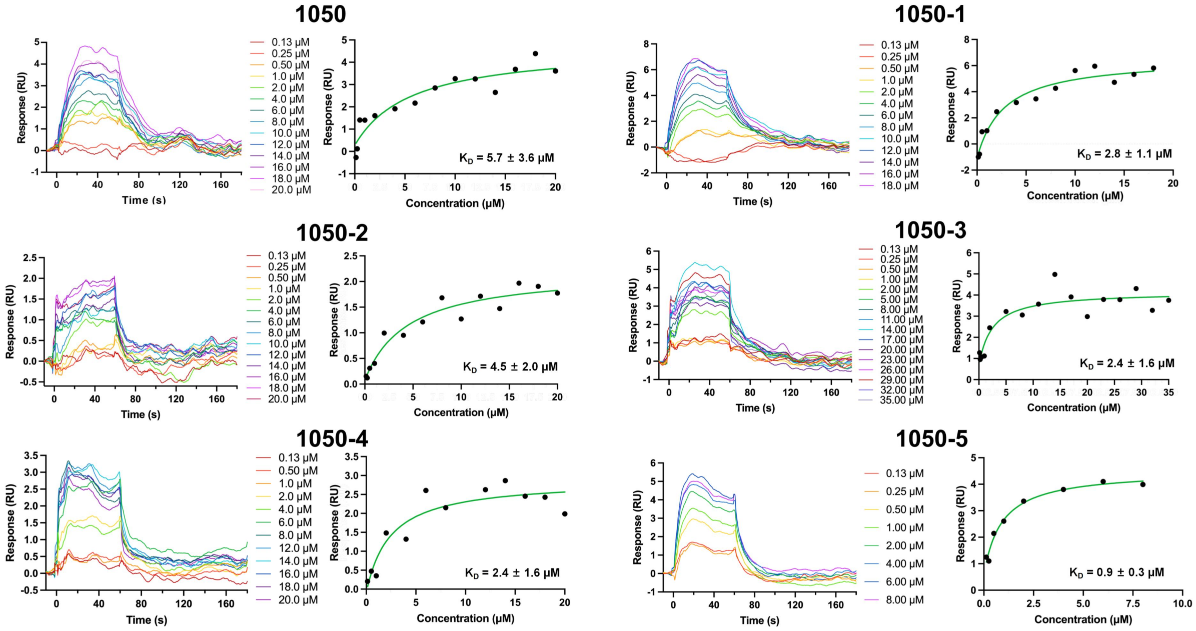


**Figure S7**. SPR binding curves and affinities of 1050-series compounds.

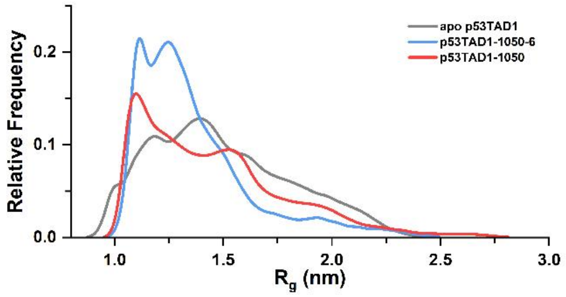


**Figure S8**. $R_g$ from MD simulation of p53TAD1 in apo, 1050-bound, and 1050-6-bound states.

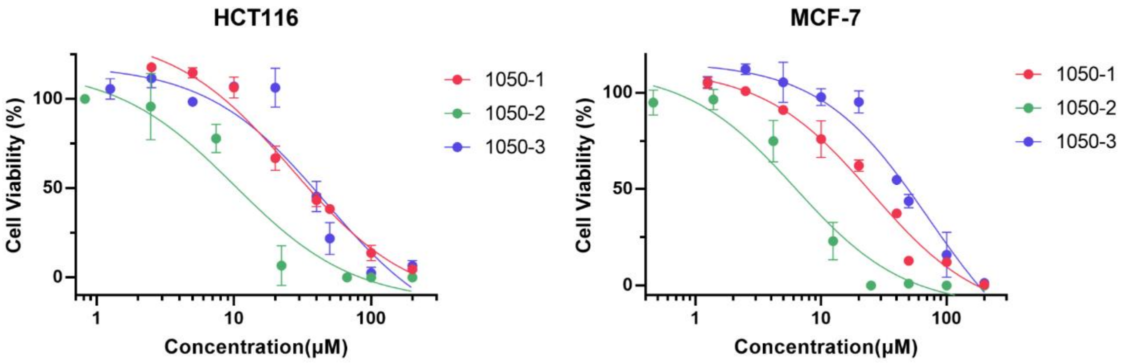


**Figure S9**. Cell viability inhibition activity of 1050-1, 1050-2, and 1050-3 in HCT116 and MCF-7 cell lines.

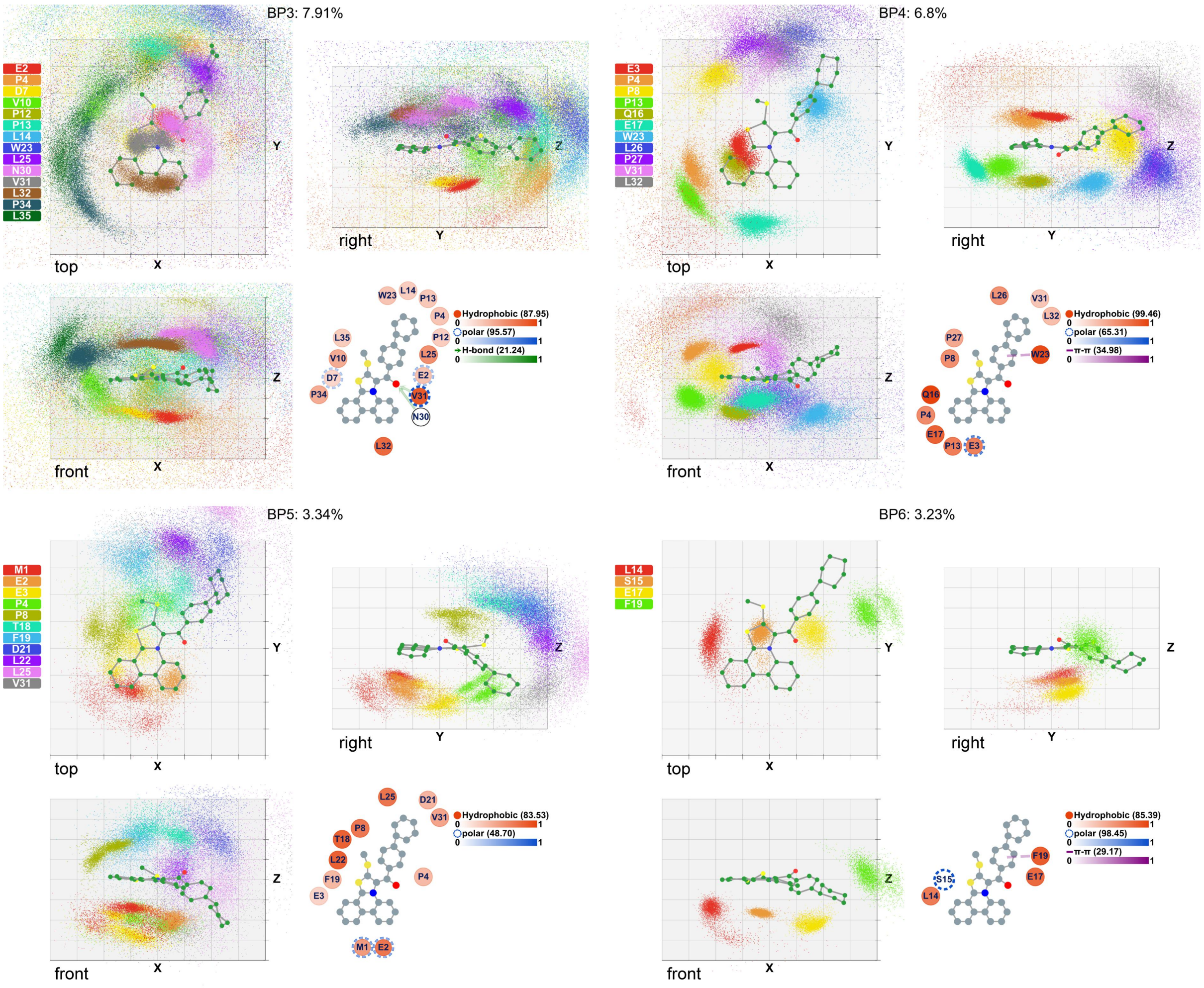


**Figure S10**. Binding patterns 3-6 of the p53TAD1-1050-6 complex. For the orientation plot, each 3D binding pattern is presented through three views, where the compound is shown in ball-and-stick and residues are shown in colored geometric centers. The grid in the background has a spacing length of 2.5 Å. For the interaction plot, the hydrophobic, polar, π-π, and hydrogen bond interactions are shown as red circles, blue **segmented circles, purple dashed lines,** and green arrows (pointing from donor to acceptor), respectively. The number of each interaction refers to its maximum probability.

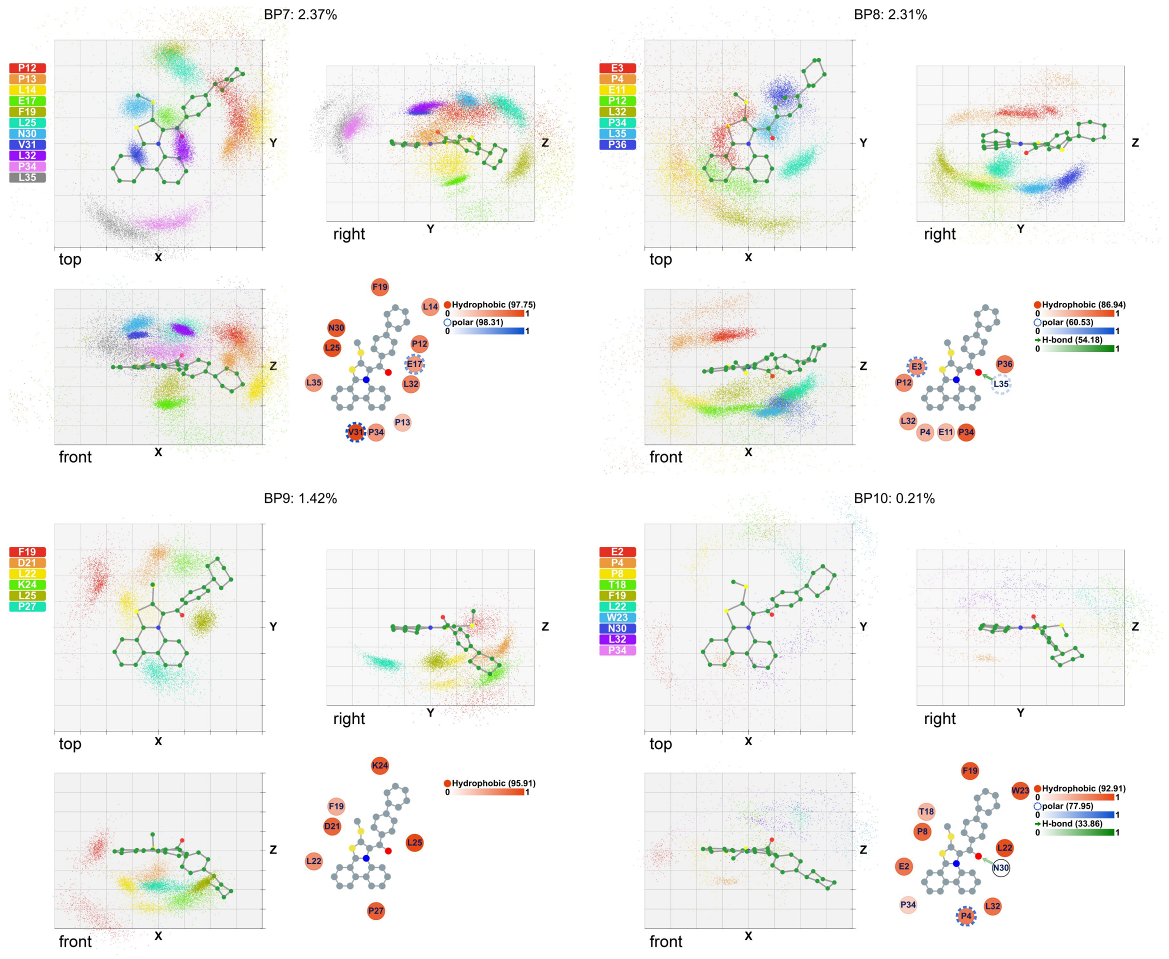


**Figure S11**. Binding patterns 7-10 of the p53TAD1-1050-6 complex. For the orientation plot, each 3D binding pattern is presented through three views, where the compound is shown in ball-and-stick and residues are shown in colored geometric centers. The grid in the background has a spacing length of 2.5 Å. For the interaction plot, the hydrophobic, polar, π-π, and hydrogen bond interactions are shown as red circles, blue segmented circles, purple dashed lines, and green arrows (pointing from donor to acceptor), respectively. The number of each interaction refers to its maximum probability.

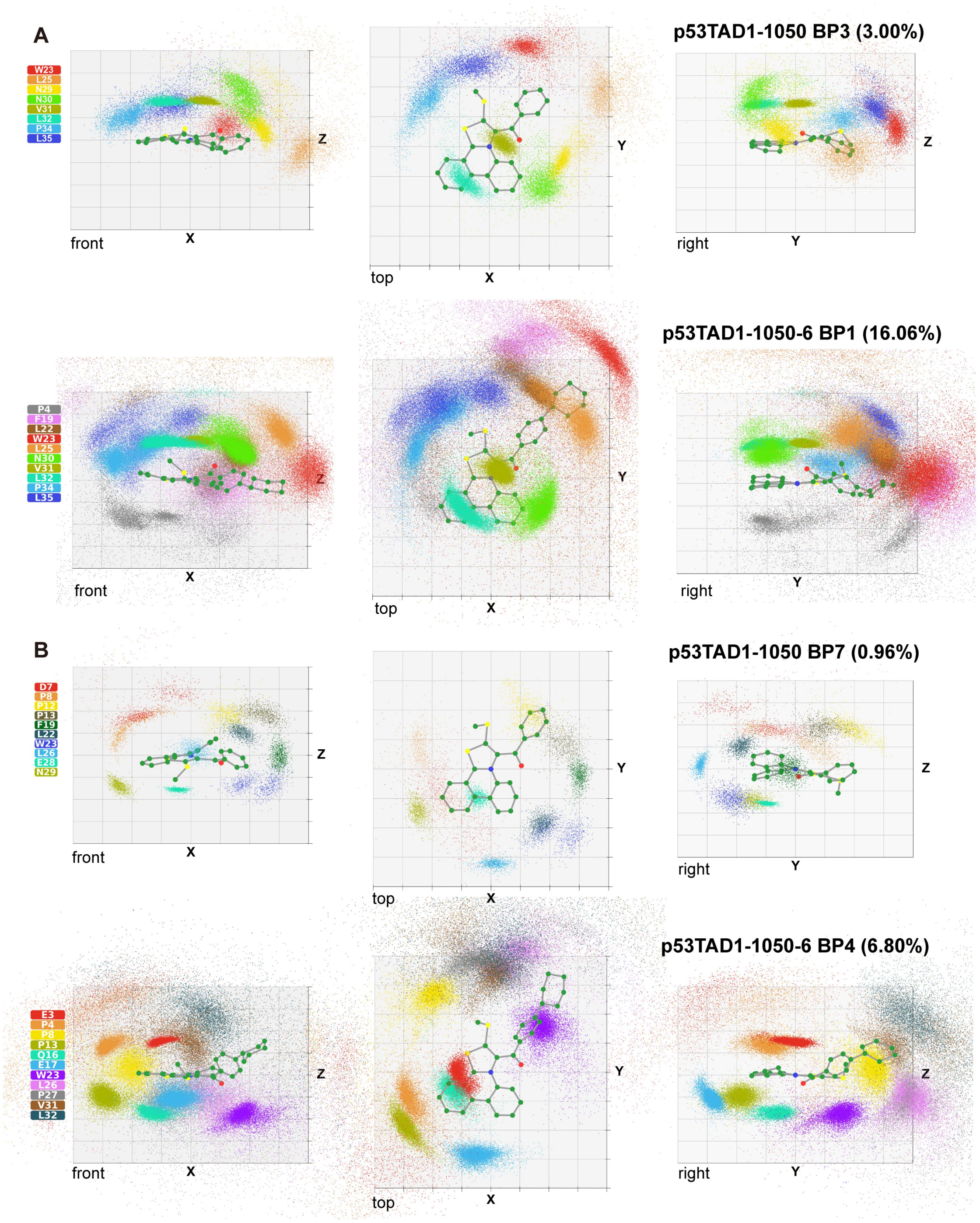


**Figure S12**. Similar binding patterns between p53TAD1-1050 and p53TAD1-1050-6. (A) p53TAD1-1050 BP3 (top) and p53TAD1-1050-6 BP1 (bottom). (B) p53TAD1-1050 BP6 (top) and p53TAD1-1050-6 BP4 (bottom). The following residues have close orientation: P8 (p53TAD1-1050 BP6) to P4 (p53TAD1-1050-6), L26 to E17, E28 to Q16, N29 to P13.

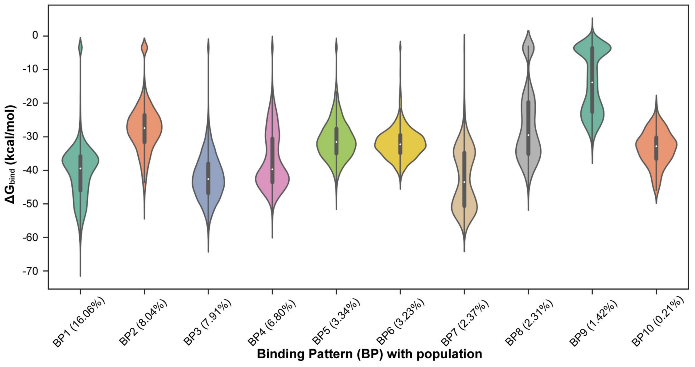


**Figure S13**. The binding free energy (*ΔG*$_{bind}$) distributions of ten p53TAD1-1050-6 predominant binding patterns.

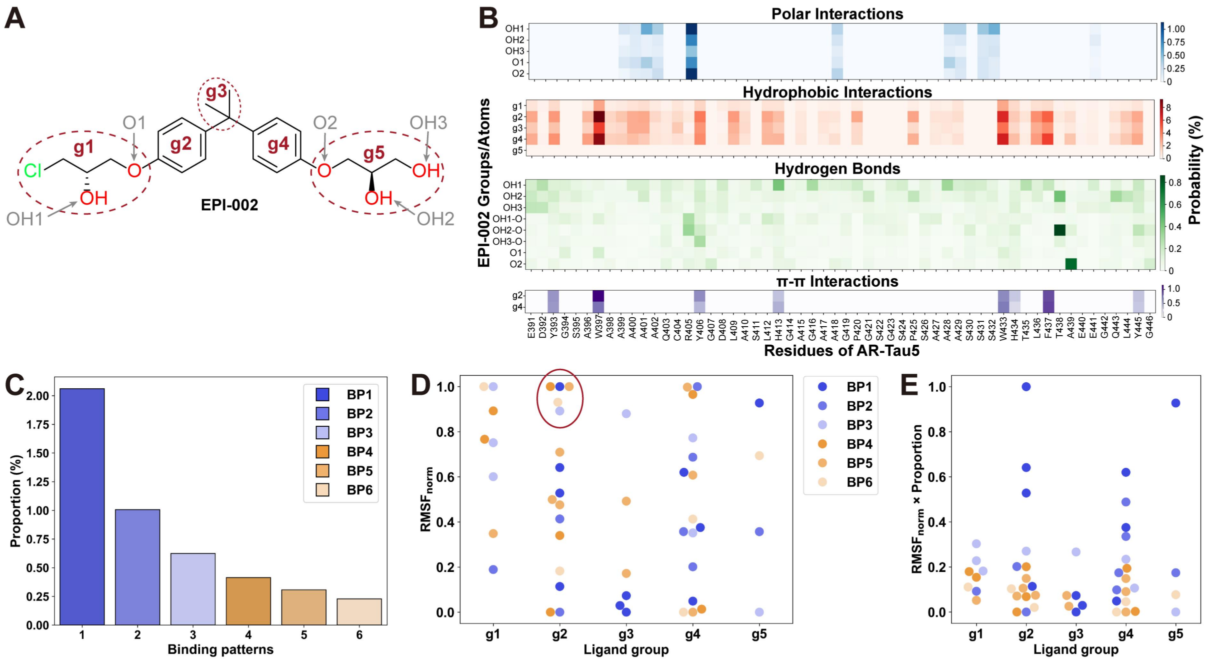


**Figure S14**. Overall and binding stability analysis of AR-Tau5 with EPI-002. (A) Definition of groups in compound EPI-002. (B) The overall interaction probabilities between AR-Tau5 and EPI-002 groups. The OH represents the OH group as the donor group, and OH-O represents the OH group as the acceptor group. (C) The proportion of predominant binding patterns in the trajectory. (D) The normalized RMSF of distances between the ligand groups and interacting residues in the predominant binding patterns. (E) The normalized RMSF is multiplied by the proportion of corresponding binding patterns. The proportion is normalized by the largest BP proportion.

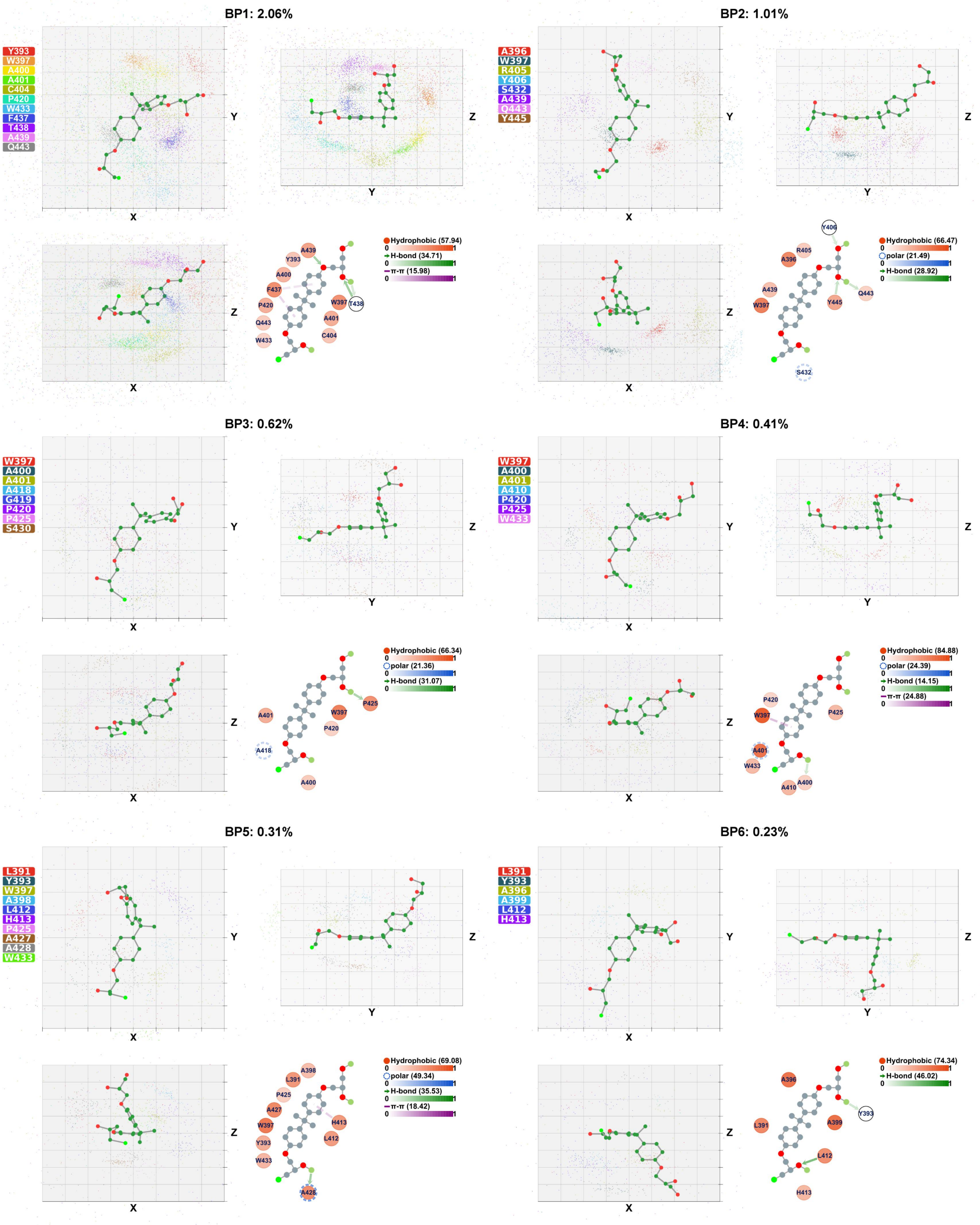


**Figure S15**. Binding patterns that are formed by AR-Tau5 and EPI-002. For the orientation plot, each 3D binding pattern is presented through three views, where the compound is shown in ball-and-stick and residues are shown in colored geometric centers. The grid in the background has a spacing length of 2.5 Å. For the interaction plot, the hydrophobic, polar, π-π, and hydrogen bond interactions are shown as red circles, blue segmented circles, purple dashed lines, and green arrows (pointing from donor to acceptor), respectively. The number of each interaction refers to its maximum probability.

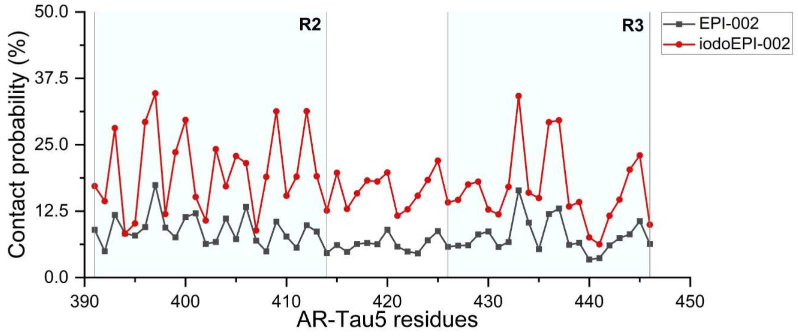


**Figure S16**. The residue contact probability of compounds EPI-002 and iodoEPI-002.

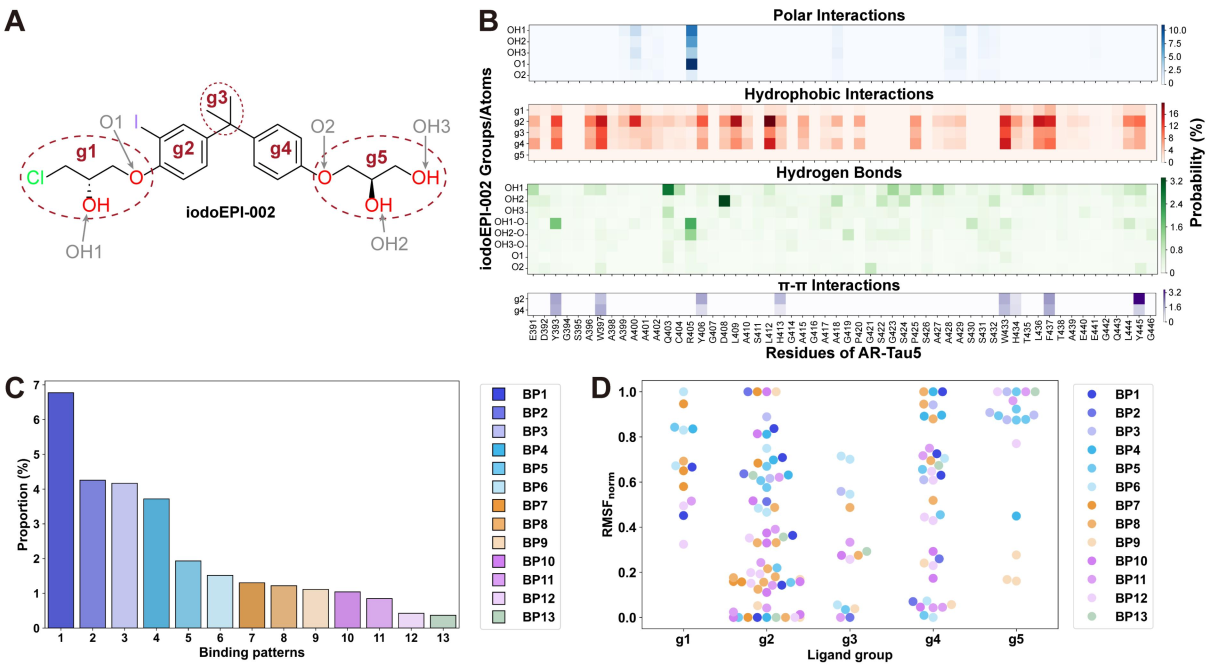


**Figure S17**. Overall and binding stability analysis of AR-Tau5 with iodoEPI-002. (A) Definition of groups in compound iodoEPI-002. (B) The overall interaction probabilities between AR-Tau5 and iodoEPI-002 groups. The OH represents the OH group as the donor group, and OH-O represents the OH group as the acceptor group. (C) The proportion of predominant binding patterns in the trajectory. (D) The normalized RMSF of distances between the ligand groups and interacting residues in the predominant binding patterns.

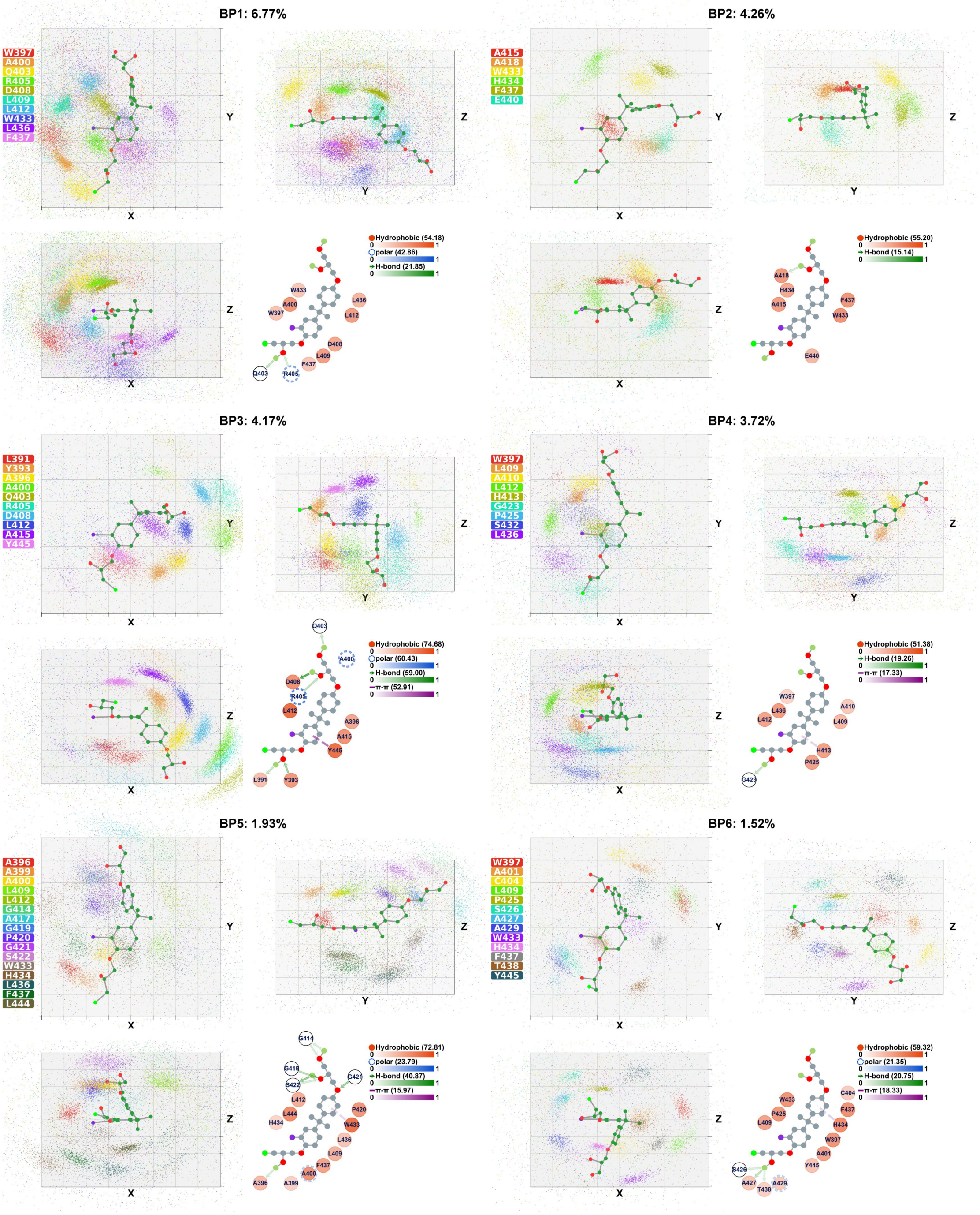

**Figure S18**. Binding patterns 1-6 that are formed by AR-Tau5 and iodoEPI-002. For the orientation plot, each 3D binding pattern is presented through three views, where the compound is shown in ball-and-stick and residues are shown in colored geometric centers. The grid in the background has a spacing length of 2.5 Å. For the interaction plot, the hydrophobic, polar, π-π, and hydrogen bond interactions are shown as red circles, blue segmented circles, purple dashed lines, and green arrows (pointing from donor to acceptor), respectively. The number of each interaction refers to its maximum probability.

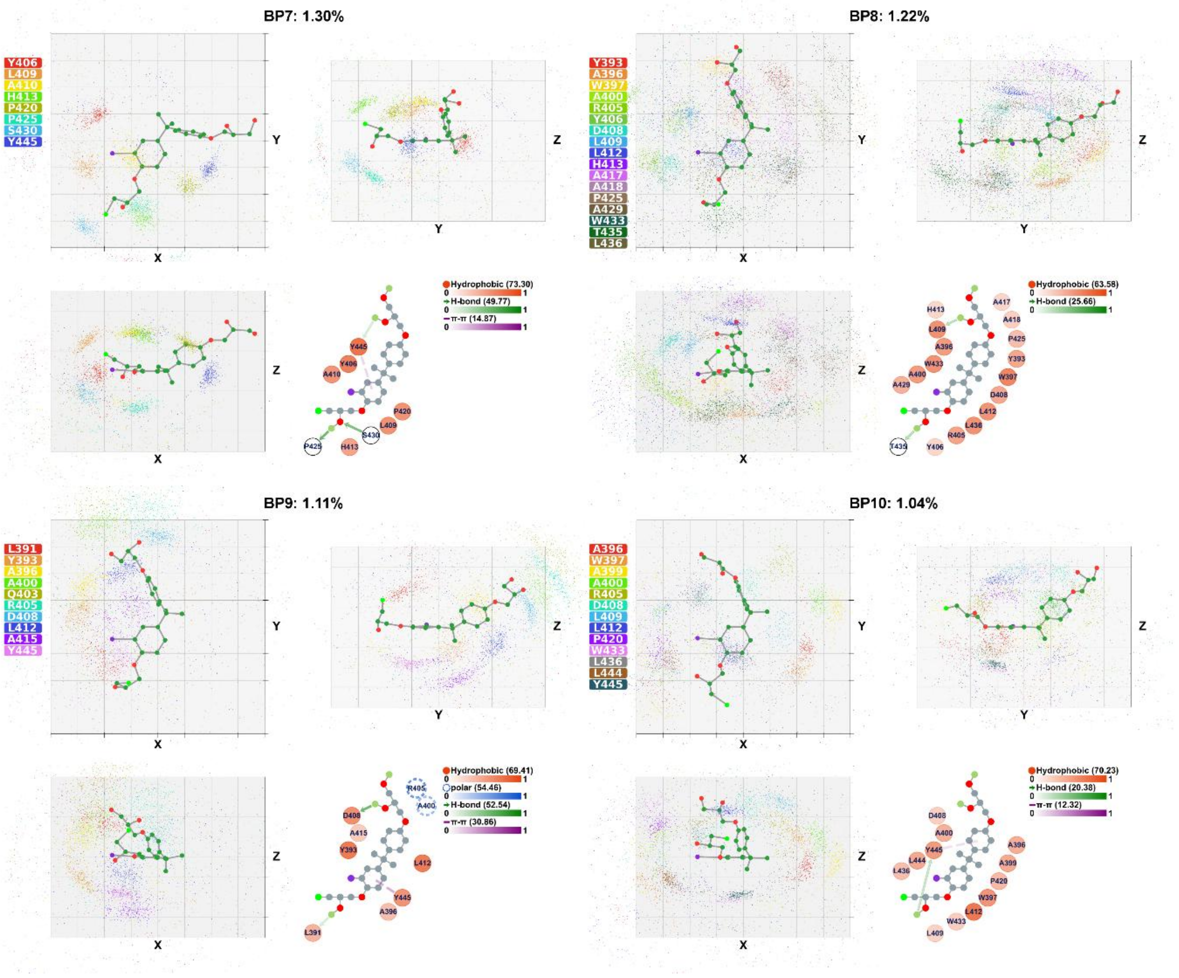


**Figure S19**. Binding patterns 7-13 that are formed by AR-Tau5 and iodoEPI-002. For the orientation plot, each 3D binding pattern is presented through three views, where the compound is shown in ball-and-stick and residues are shown in colored geometric centers. The grid in the background has a spacing length of 2.5 Å. For the interaction plot, the hydrophobic, polar, π-π, and hydrogen bond interactions are shown as red circles, blue segmented circles, purple dashed lines, and green arrows (pointing from donor to acceptor), respectively. The number of each interaction refers to its maximum probability.

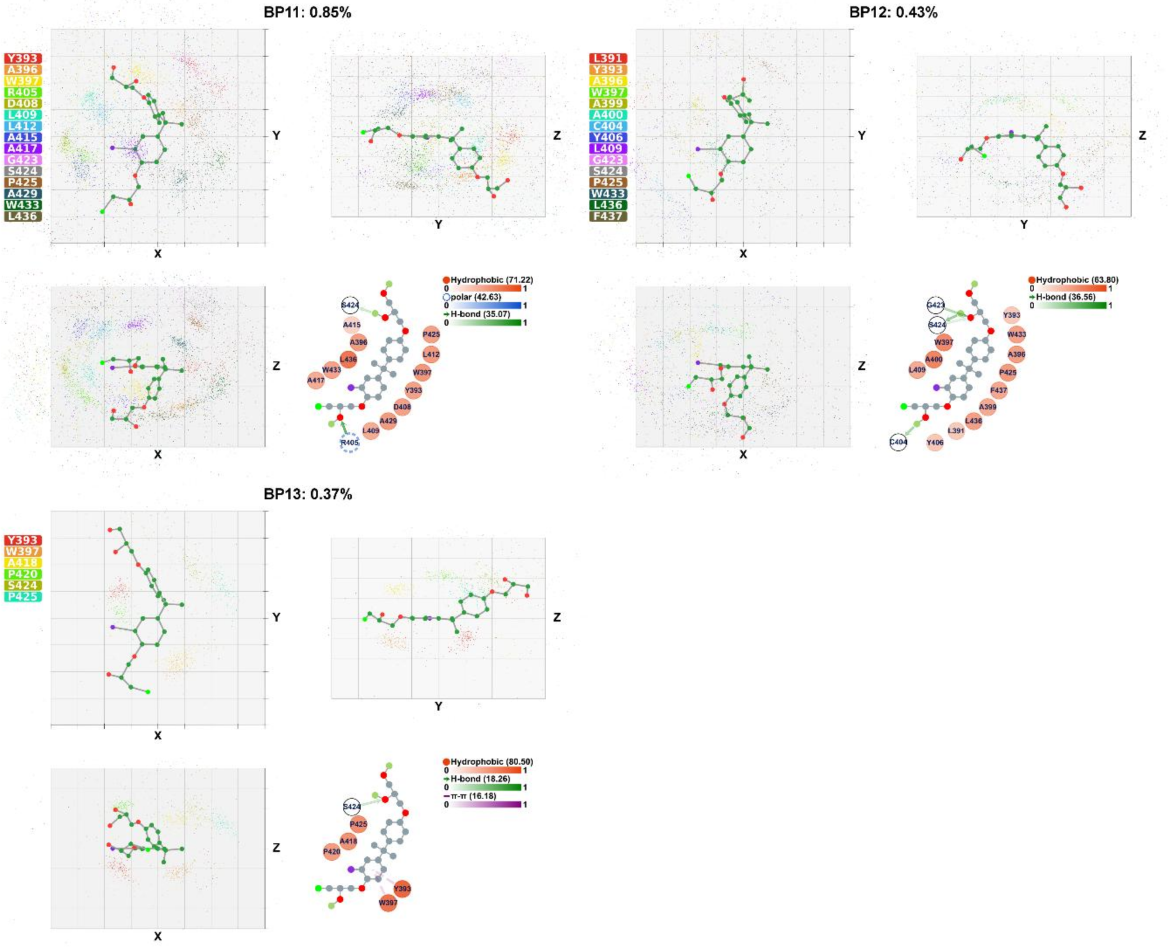


**Figure S20**. Binding patterns 7-13 that are formed by AR-Tau5 and iodoEPI-002. For the orientation plot, each 3D binding pattern is presented through three views, where the compound is shown in ball-and-stick and residues are shown in colored geometric centers. The grid in the background has a spacing length of 2.5 Å. For the interaction plot, the hydrophobic, polar, π-π, and hydrogen bond interactions are shown as red circles, blue segmented circles, purple dashed lines, and green arrows (pointing from donor to acceptor), respectively. The number of each interaction refers to its maximum probability.

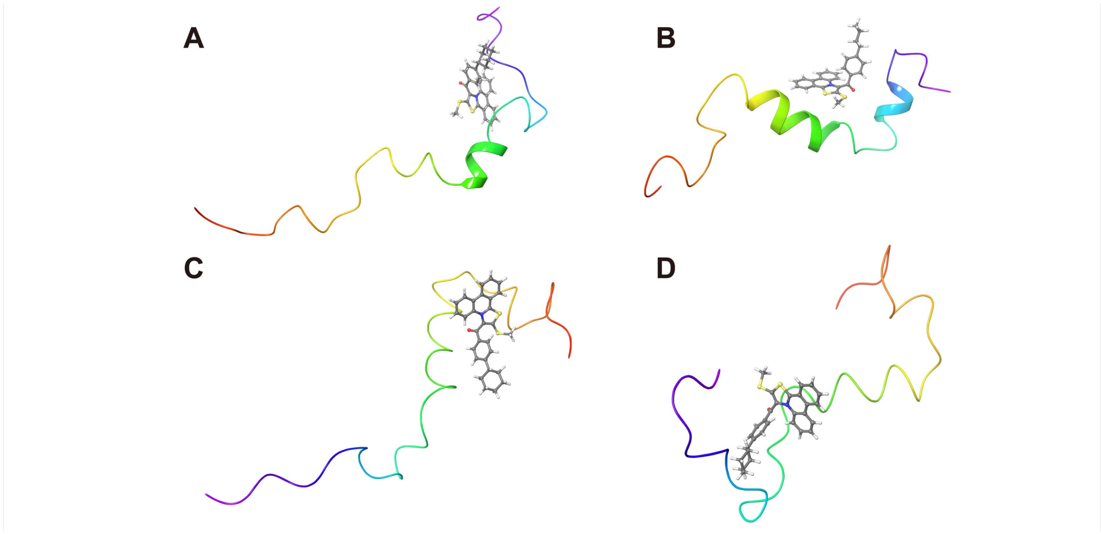


**Figure S21**. Initial structures for p53TAD1-1050-6 MD simulations. p53TAD1 is shown in the rainbow-colored cartoon with the N-terminus in blue and the C-terminus in red. Compound 1050-6 is shown in ball-and-stick.

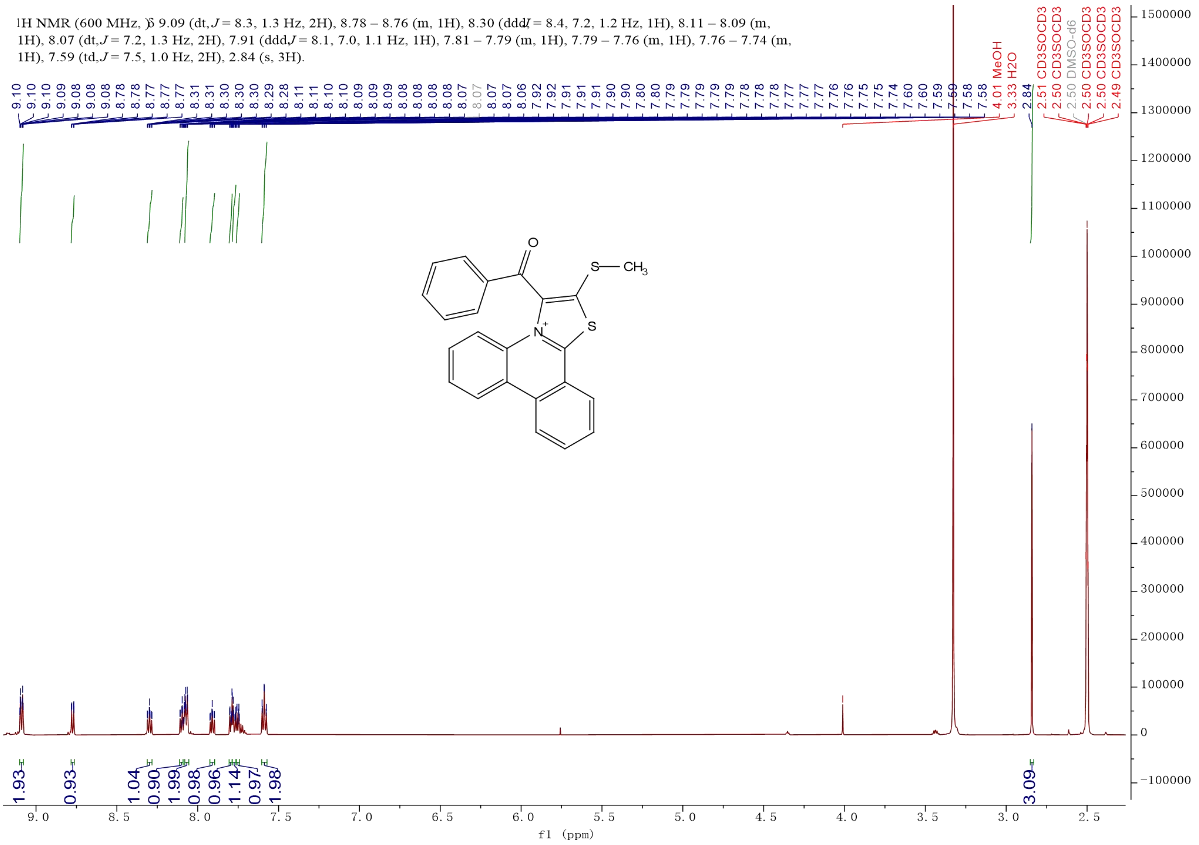


**Figure S22**. ¹H NMR of compound 1050.

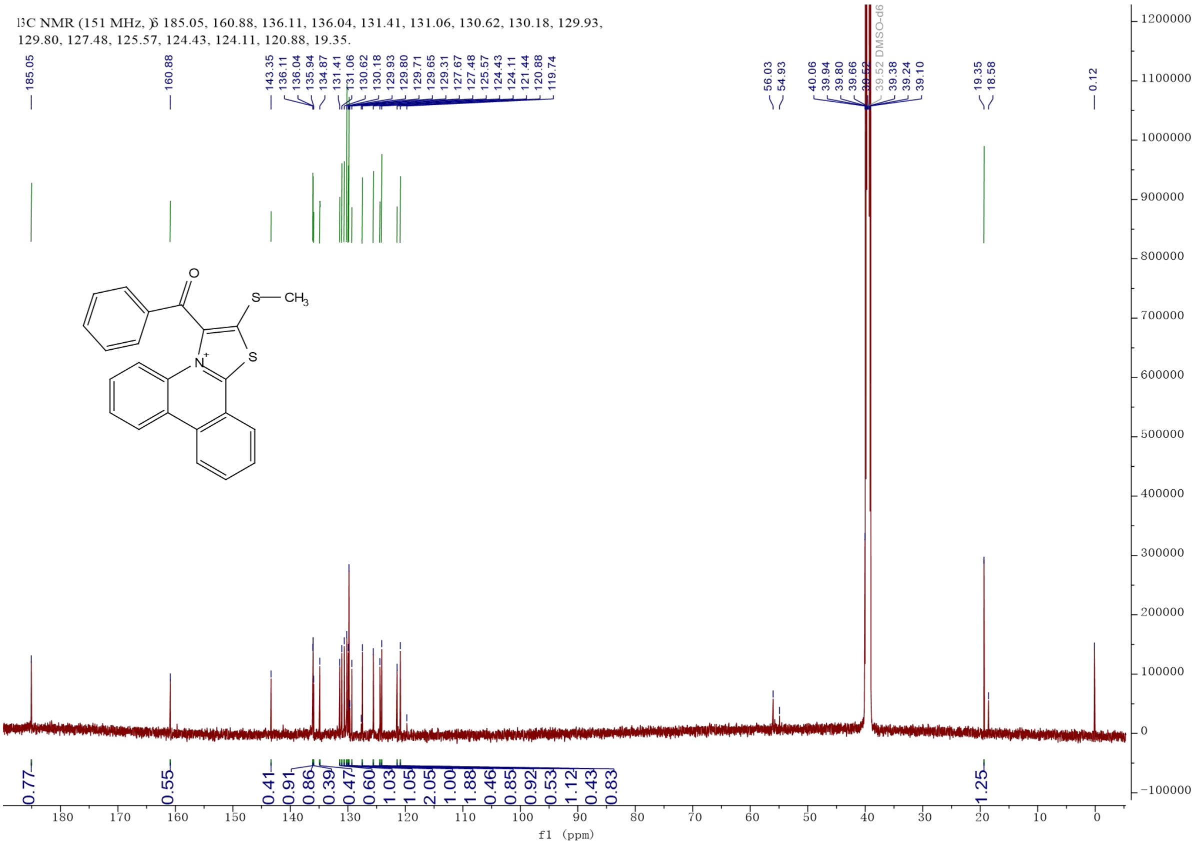


**Figure S23**. ¹³C NMR of compound 1050.

### Peking University Mass Spectrometry Sample Analysis Report

**Analysis Info**

| | | | |
|---|---|---|---|
| Analysis Name | FTMS-25010074_Pos_20250120_000002.d | Acquisition Date | 1/20/2025 9:38:02 AM |
| Sample | 1050 | Instrument | Bruker Solarix XR FTMS |
| Comment | | Operator | Peking University |

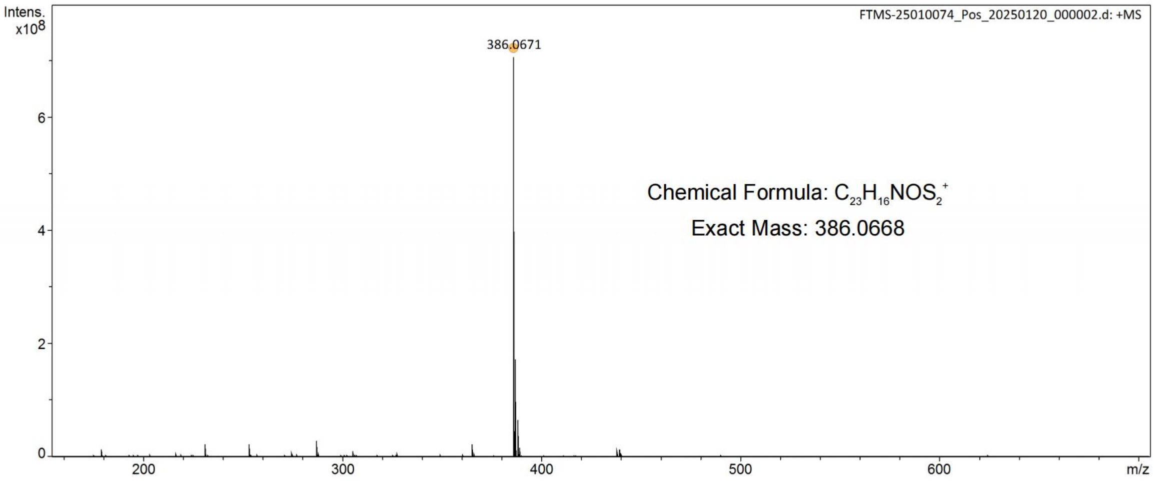


| Meas. m/z | # | Ion Formula | Score | m/z | err [ppm] | Mean err [ppm] | mSigma | rdb | e⁻ Conf | N-Rule |
|---|---|---|---|---|---|---|---|---|---|---|
| 386.0671 | 1 | C23H16NOS2 | 100.00 | 386.0668 | -0.7 | -1.4 | 14.0 | 21.0 | even | ok |

**Figure S24**. High-resolution mass spectrometry spectrum of compound 1050.

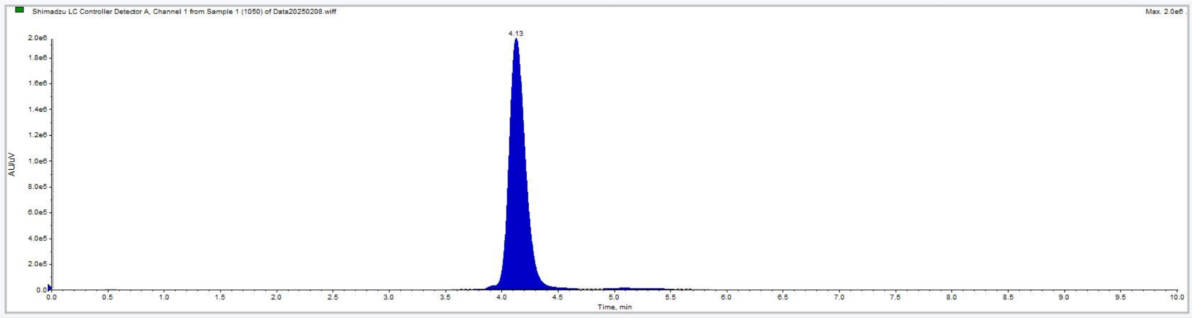


Shimadzu LC Controller Detector A, Channel 1 from Sample 1 (1050) of Data20250208.wiff

Data List Peak List

| | Time (min) | Area (counts) | % Area | Height | % Height | Width (min) | Baseline Type |
|---|---|---|---|---|---|---|---|
| 1 | 0.2358 | 428.5000 | 2.1391e-3 | 100.3840 | 5.0562e-3 | 0.1383 | Base to Base |
| 2 | 0.5177 | 6.4815e4 | 0.3236 | 9930.4217 | 0.5002 | 0.8933 | Base to Base |
| 3 | 1.4792 | 4078.5000 | 0.0204 | 317.2780 | 0.0160 | 0.3650 | Base to Base |
| 4 | 2.7408 | 2193.1000 | 0.0109 | 202.7118 | 0.0102 | 0.4033 | Base to Base |
| 5 | 3.2133 | 1.0852e4 | 0.0542 | 1278.9497 | 0.0644 | 0.3317 | Base to Base |
| 6 | 3.6225 | 1.5741e4 | 0.0786 | 2036.2373 | 0.1026 | 0.2633 | Base to Base |
| 7 | 4.1285 | 1.9516e7 | 97.4231 | 1.9477e6 | 98.1001 | 0.9267 | Valley |
| 8 | 4.8525 | 1.0147e5 | 0.5065 | 7665.8608 | 0.3861 | 0.2283 | Valley |
| 9 | 5.0908 | 1.9162e5 | 0.9566 | 8543.4592 | 0.4303 | 0.6117 | Valley |
| 10 | 5.6630 | 3.1460e4 | 0.1570 | 3618.3536 | 0.1823 | 0.3733 | Base to Base |
| 11 | 6.8150 | 5.6862e4 | 0.2839 | 1673.5840 | 0.0843 | 1.3300 | Base to Base |
| 12 | 7.9075 | 1.5202e4 | 0.0759 | 480.6597 | 0.0242 | 0.9617 | Base to Base |
| 13 | 9.4058 | 8998.7462 | 0.0449 | 481.4500 | 0.0242 | 0.5600 | Valley |
| 14 | 9.4192 | 1.1576e4 | 0.0578 | 487.6248 | 0.0246 | 0.5650 | Valley |
| 15 | 9.9908 | 900.8500 | 4.4971e-3 | 902.8750 | 0.0455 | 0.0167 | Base to Base |

**Figure S25**. HPLC spectrum of compound 1050.

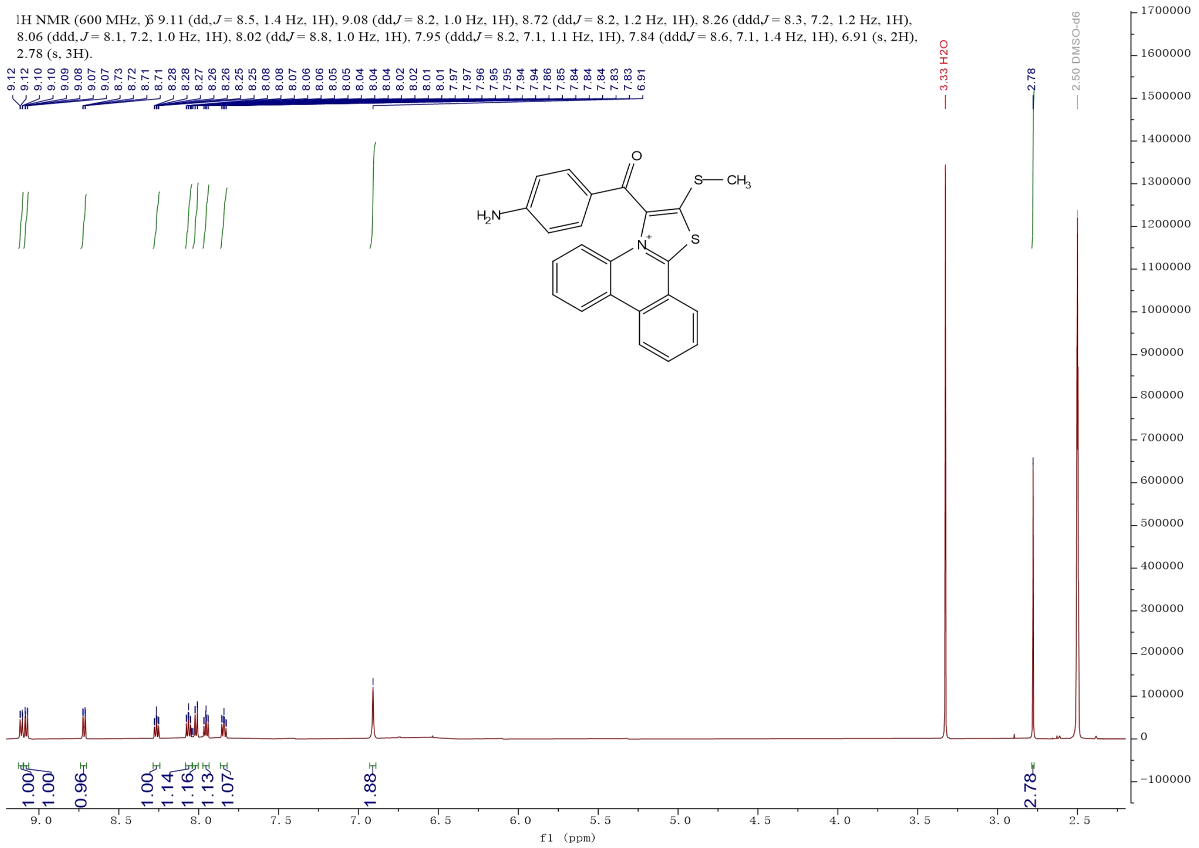


**Figure S26**. $^{1}H$ NMR of compound 1050-1.

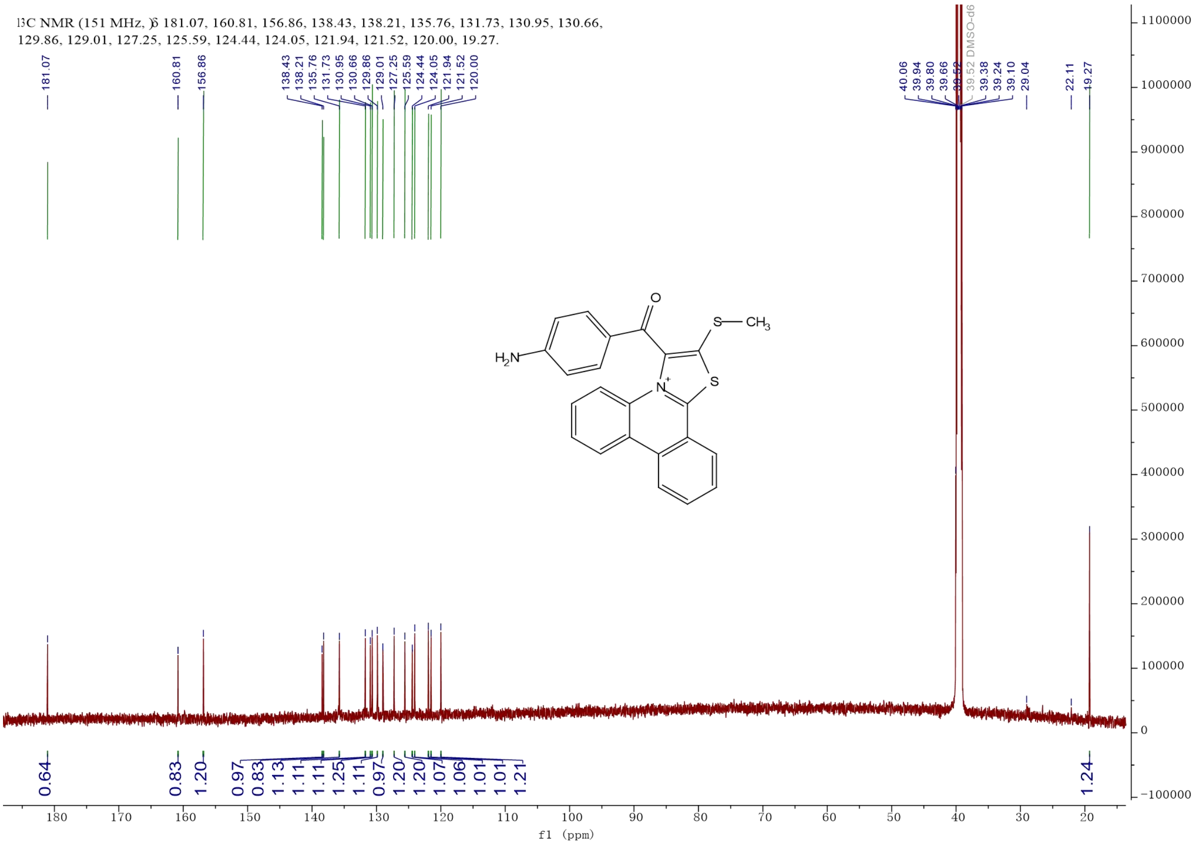


**Figure S27**. $^{13}C$ NMR of compound 1050-1.

### Peking University Mass Spectrometry Sample Analysis Report

**Analysis Info**

| | | | |
|---|---|---|---|
| Analysis Name | FTMS-25010074_Pos_20250120_000005.d | Acquisition Date | 1/20/2025 9:46:51 AM |
| Sample | 1050-1 | Instrument | Bruker Solarix XR FTMS |
| Comment | | Operator | Peking University |

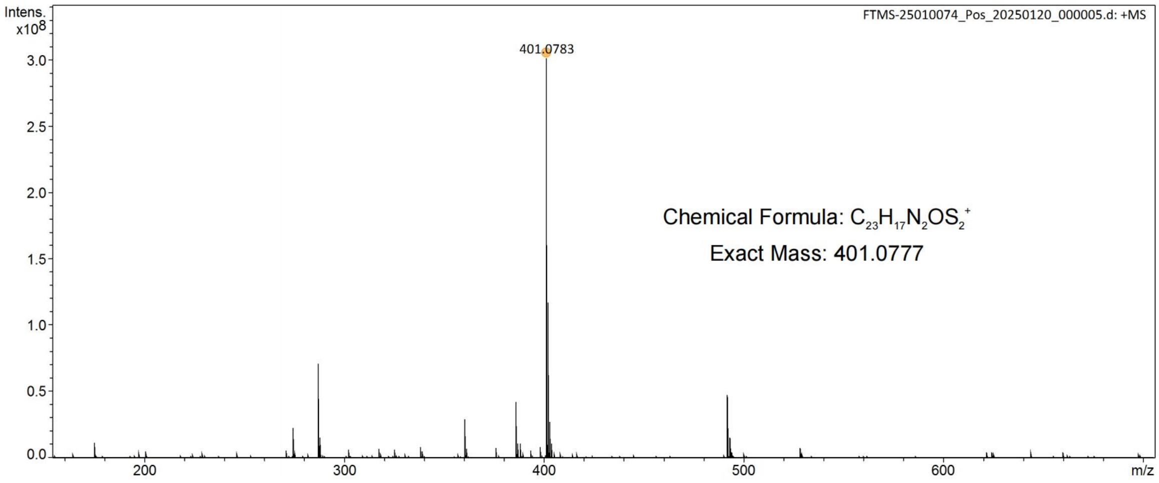


| Meas. m/z | # | Ion Formula | Score | m/z | err [ppm] | Mean err [ppm] | mSigma | rdb | e⁻ Conf | N-Rule |
|---|---|---|---|---|---|---|---|---|---|---|
| 401.0783 | 1 | C23H17N2OS2 | 100.00 | 401.0777 | -1.5 | -1.8 | 18.8 | 21.0 | even | ok |

**Figure S28**. High-resolution mass spectrometry spectrum of compound 1050-1.

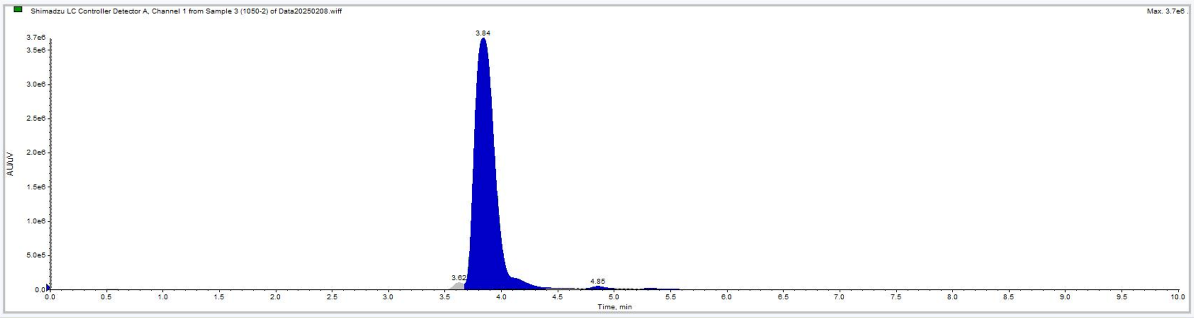


Shimadzu LC Controller Detector A, Channel 1 from Sample 3 (1050-2) of Data20250208.wiff

Data List Peak List

| | Time (min) | Area (counts) | % Area | Height | % Height | Width (min) | Baseline Type |
|---|---|---|---|---|---|---|---|
| 1 | 0.2163 | 1.0532e4 | 0.0224 | 1132.1371 | 0.0293 | 0.4467 | Base to Base |
| 2 | 0.5472 | 1.4193e5 | 0.3025 | 1.8880e4 | 0.4888 | 0.5750 | Base to Base |
| 3 | 1.4442 | 9672.8000 | 0.0206 | 626.2675 | 0.0162 | 0.5967 | Base to Base |
| 4 | 2.0992 | 2758.3000 | 5.8783e-3 | 368.8068 | 9.5489e-3 | 0.2750 | Base to Base |
| 5 | 2.6942 | 5314.4500 | 0.0113 | 481.6667 | 0.0125 | 0.4200 | Base to Base |
| 6 | 3.2608 | 745.0000 | 1.5877e-3 | 172.0194 | 4.4538e-3 | 0.1183 | Base to Base |
| 7 | 3.4108 | 2.3642e4 | 0.0504 | 3718.2731 | 0.0963 | 0.1983 | Base to Base |
| 8 | 3.6249 | 5.6340e5 | 1.2007 | 9.7827e4 | 2.5329 | 0.1633 | Valley |
| 9 | 3.8396 | 4.5621e7 | 97.2248 | 3.6735e6 | 95.1110 | 0.9650 | Valley |
| 10 | 4.7272 | 2.6285e4 | 0.0560 | 6101.5085 | 0.1580 | 0.1167 | Valley |
| 11 | 4.8533 | 2.7833e5 | 0.5931 | 3.4508e4 | 0.8934 | 0.2517 | Valley |
| 12 | 5.0556 | 6.1492e4 | 0.1310 | 6972.2513 | 0.1805 | 0.2233 | Valley |
| 13 | 5.3142 | 9.9603e4 | 0.2123 | 1.3180e4 | 0.3412 | 0.3533 | Base to Base |
| 14 | 7.0065 | 4.3943e4 | 0.0936 | 3924.3914 | 0.1016 | 0.8833 | Base to Base |
| 15 | 7.9725 | 7295.5000 | 0.0155 | 258.0557 | 6.6814e-3 | 0.5533 | Base to Base |
| 16 | 9.4175 | 2.7280e4 | 0.0581 | 676.6196 | 0.0175 | 1.2317 | Base to Base |

**Figure S29**. HPLC spectrum of compound 1050-1.

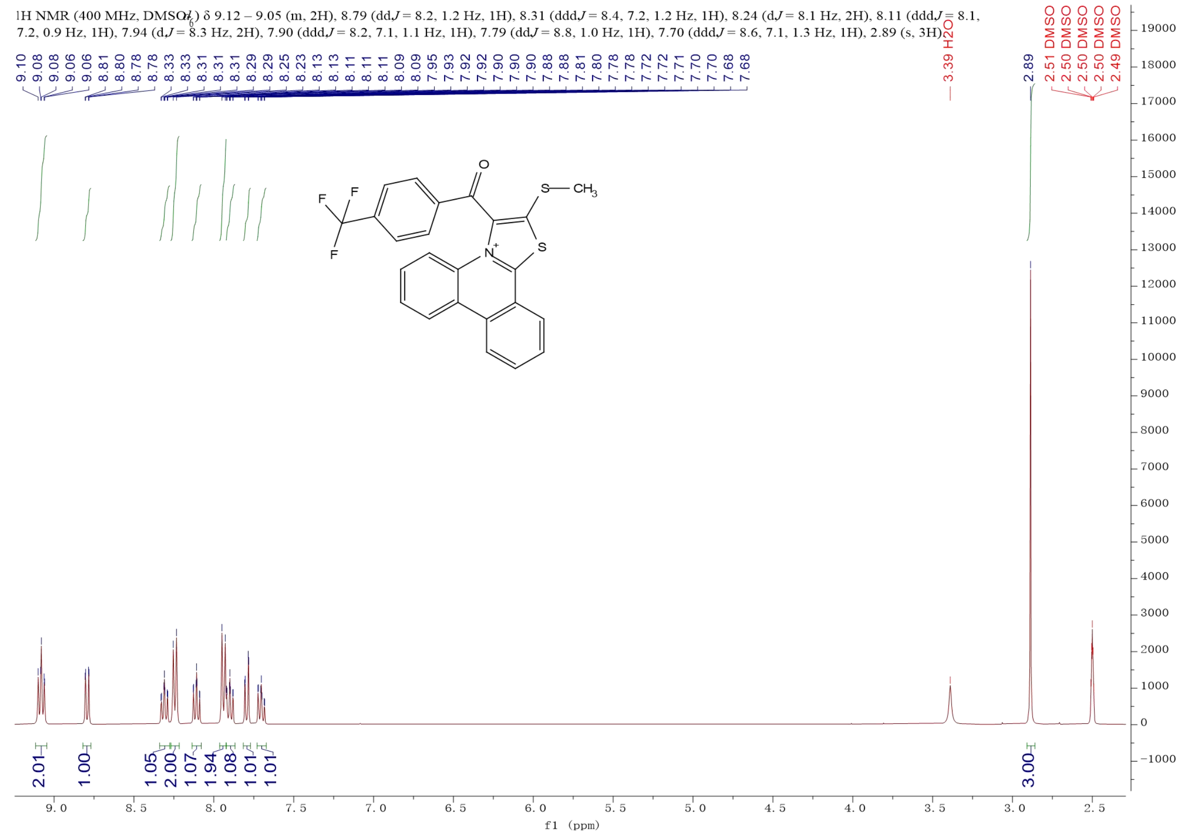


**Figure S30**. $^{1}$H NMR of compound 1050-2.

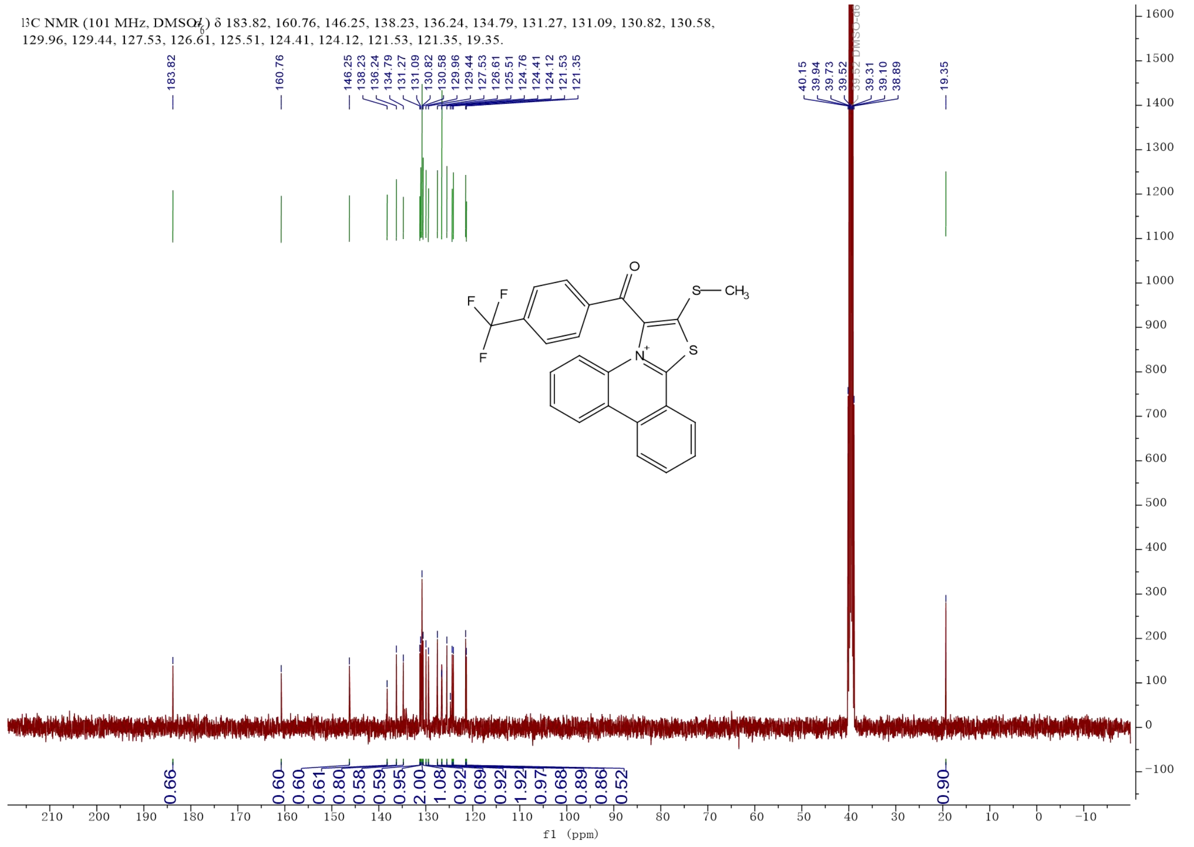


**Figure S31**. $^{13}$C NMR of compound 1050-2.

### Peking University Mass Spectrometry Sample Analysis Report

**Analysis Info**

| | | | |
|---|---|---|---|
| Analysis Name | FTMS-25010074_Pos_20250120_000007.d | Acquisition Date | 1/20/2025 9:51:40 AM |
| Sample | 1050-2 | Instrument | Bruker Solarix XR FTMS |
| Comment | | Operator | Peking University |

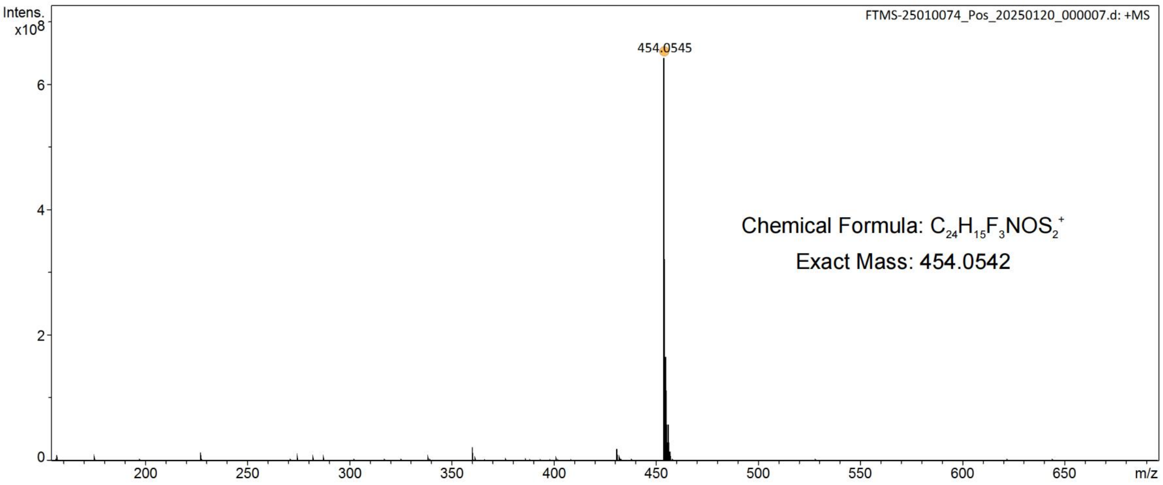


| Meas. m/z | # | Ion Formula | Score | m/z | err [ppm] | Mean err [ppm] | mSigma | rdb | e⁻ Conf | N-Rule |
|---|---|---|---|---|---|---|---|---|---|---|
| 454.0545 | 1 | C24H15F3NOS2 | 100.00 | 454.0542 | -0.8 | -1.0 | 20.6 | 21.0 | even | ok |

**Figure S32**. High-resolution mass spectrometry spectrum of compound 1050-2.

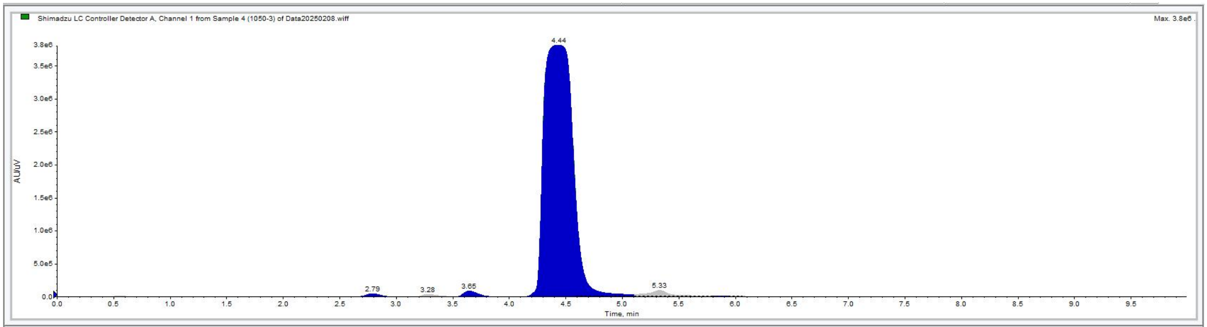


Shimadzu LC Controller Detector A, Channel 1 from Sample 4 (1050-3) of Data20250208.wiff

Data List Peak List

| | Time (min) | Area (counts) | % Area | Height | % Height | Width (min) | Baseline Type |
|---|---|---|---|---|---|---|---|
| 1 | 0.2200 | 1.2414e4 | 0.0173 | 1536.2880 | 0.0373 | 0.3067 | Base to Base |
| 2 | 0.5483 | 1.3307e5 | 0.1859 | 1.7871e4 | 0.4341 | 0.5200 | Base to Base |
| 3 | 1.3992 | 5990.6000 | 8.3703e-3 | 423.9897 | 0.0103 | 0.4250 | Base to Base |
| 4 | 2.3925 | 7359.0000 | 0.0103 | 894.9397 | 0.0217 | 0.3283 | Base to Base |
| 5 | 2.7891 | 4.4817e5 | 0.6262 | 5.0291e4 | 1.2215 | 0.6150 | Base to Base |
| 6 | 3.2816 | 3.3550e5 | 0.4688 | 3.7514e4 | 0.9112 | 0.3683 | Base to Base |
| 7 | 3.6463 | 7.3495e5 | 1.0269 | 8.6027e4 | 2.0896 | 0.3400 | Valley |
| 8 | 3.9436 | 1.8791e4 | 0.0263 | 2833.5612 | 0.0688 | 0.1983 | Valley |
| 9 | 4.4373 | 6.8450e7 | 95.6408 | 3.8120e6 | 92.5900 | 1.0383 | Valley |
| 10 | 5.3304 | 1.2068e6 | 1.6862 | 8.8133e4 | 2.1407 | 0.6883 | Valley |
| 11 | 5.8695 | 3.3745e4 | 0.0472 | 5780.7819 | 0.1404 | 0.1450 | Valley |
| 12 | 5.9785 | 5.2009e4 | 0.0727 | 5988.1702 | 0.1454 | 0.2367 | Valley |
| 13 | 6.2952 | 2.4181e4 | 0.0338 | 3115.3487 | 0.0757 | 0.3233 | Base to Base |
| 14 | 6.9758 | 7.5509e4 | 0.1055 | 3842.5364 | 0.0933 | 0.9717 | Base to Base |
| 15 | 8.0342 | 3802.2500 | 5.3126e-3 | 154.0857 | 3.7426e-3 | 0.4033 | Base to Base |
| 16 | 9.4275 | 2.7550e4 | 0.0385 | 666.0785 | 0.0162 | 1.2183 | Base to Base |

**Figure S33**. HPLC spectrum of compound 1050-2.

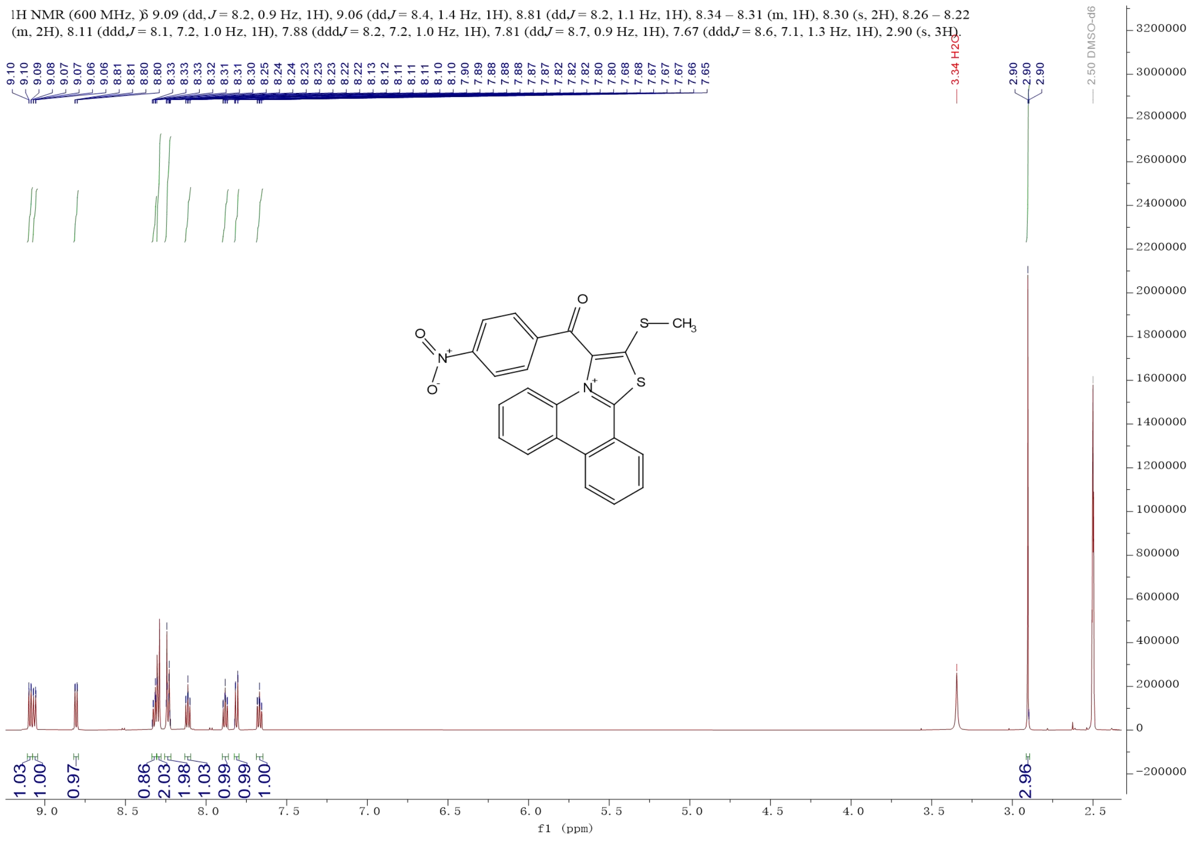


**Figure S34**. $^{1}$H NMR of compound 1050-3.

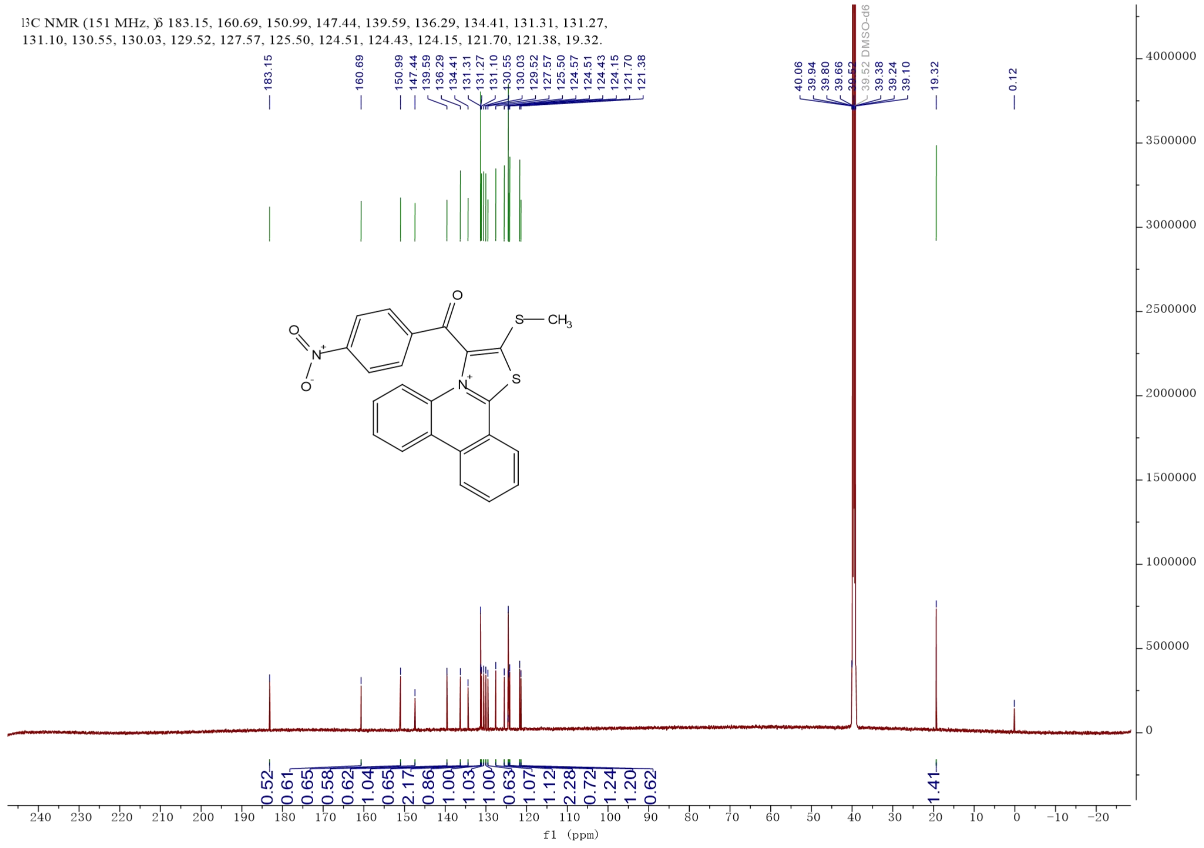


**Figure S35**. $^{13}$C NMR of compound 1050-3.

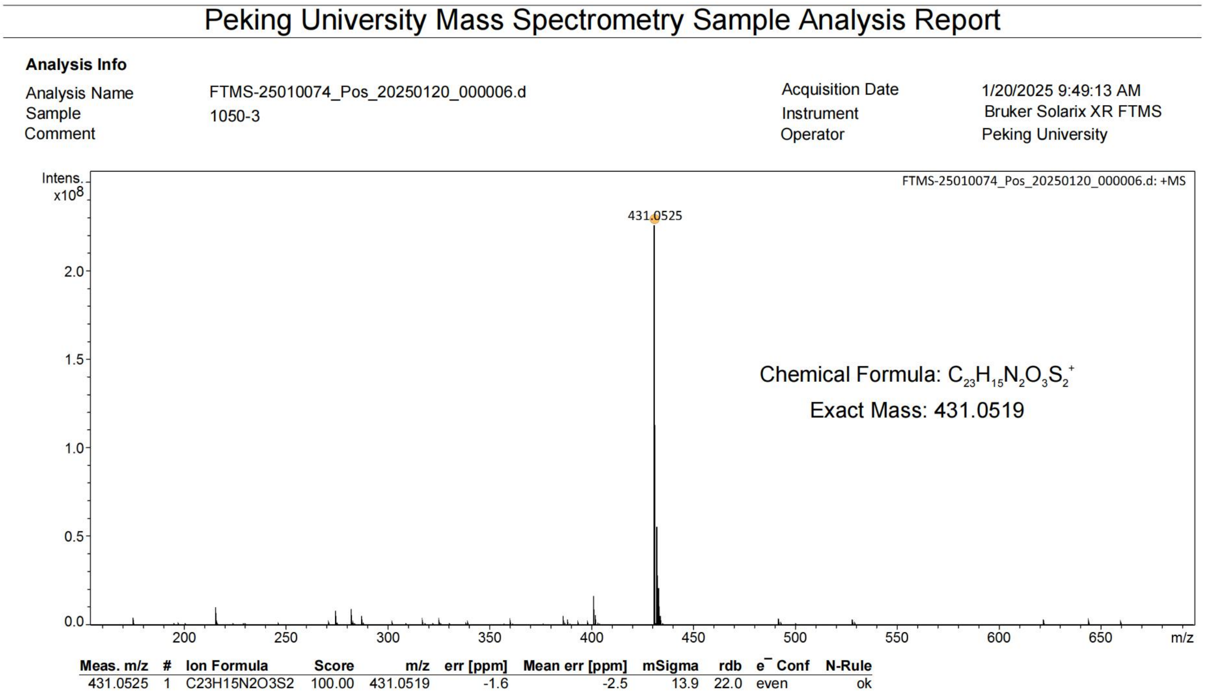
Peking University Mass Spectrometry Sample Analysis Report

**Analysis Info**

| | | | |
|---|---|---|---|
| Analysis Name | FTMS-25010074_Pos_20250120_000006.d | Acquisition Date | 1/20/2025 9:49:13 AM |
| Sample | 1050-3 | Instrument | Bruker Solarix XR FTMS |
| Comment | | Operator | Peking University |



| Meas. m/z | # | Ion Formula | Score | m/z | err [ppm] | Mean err [ppm] | mSigma | rdb | e⁻ Conf | N-Rule |
|---|---|---|---|---|---|---|---|---|---|---|
| 431.0525 | 1 | C23H15N2O3S2 | 100.00 | 431.0519 | -1.6 | -2.5 | 13.9 | 22.0 | even | ok |

**Figure S36**. High-resolution mass spectrometry spectrum of compound 1050-3.

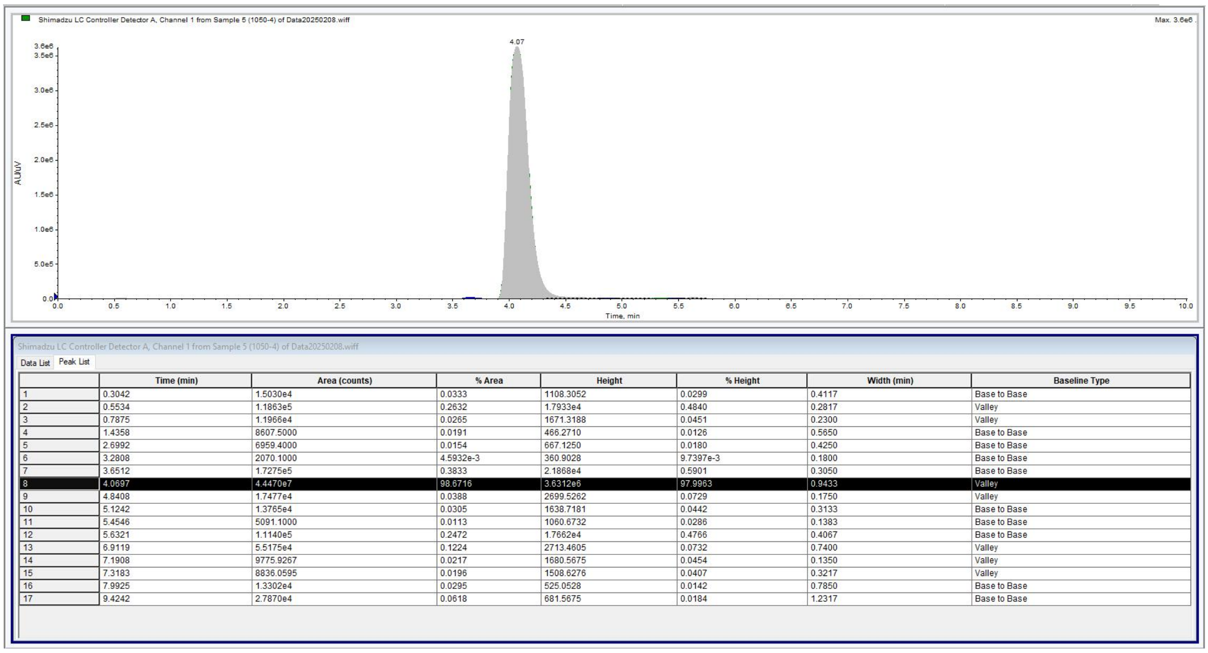


Shimadzu LC Controller Detector A, Channel 1 from Sample 5 (1050-4) of Data20250208.wiff

Data List Peak List

| | Time (min) | Area (counts) | % Area | Height | % Height | Width (min) | Baseline Type |
|---|---|---|---|---|---|---|---|
| 1 | 0.3042 | 1.5030e4 | 0.0333 | 1108.3052 | 0.0299 | 0.4117 | Base to Base |
| 2 | 0.5534 | 1.1863e5 | 0.2632 | 1.7933e4 | 0.4840 | 0.2817 | Valley |
| 3 | 0.7875 | 1.1966e4 | 0.0265 | 1671.3188 | 0.0451 | 0.2300 | Valley |
| 4 | 1.4358 | 8607.5000 | 0.0191 | 466.2710 | 0.0126 | 0.5650 | Base to Base |
| 5 | 2.6992 | 6959.4000 | 0.0154 | 667.1250 | 0.0180 | 0.4250 | Base to Base |
| 6 | 3.2808 | 2070.1000 | 4.5932e-3 | 360.9028 | 9.7397e-3 | 0.1800 | Base to Base |
| 7 | 3.6512 | 1.7275e5 | 0.3833 | 2.1868e4 | 0.5901 | 0.3050 | Base to Base |
| 8 | 4.0697 | 4.4470e7 | 98.6716 | 3.6312e6 | 97.9963 | 0.9433 | Valley |
| 9 | 4.8408 | 1.7477e4 | 0.0388 | 2699.5262 | 0.0729 | 0.1750 | Valley |
| 10 | 5.1242 | 1.3765e4 | 0.0305 | 1638.7181 | 0.0442 | 0.3133 | Base to Base |
| 11 | 5.4546 | 5091.1000 | 0.0113 | 1060.6732 | 0.0286 | 0.1383 | Base to Base |
| 12 | 5.6321 | 1.1140e5 | 0.2472 | 1.7662e4 | 0.4766 | 0.4067 | Base to Base |
| 13 | 6.9119 | 5.5175e4 | 0.1224 | 2713.4605 | 0.0732 | 0.7400 | Valley |
| 14 | 7.1908 | 9775.9267 | 0.0217 | 1680.5675 | 0.0454 | 0.1350 | Valley |
| 15 | 7.3183 | 8836.0595 | 0.0196 | 1508.6276 | 0.0407 | 0.3217 | Valley |
| 16 | 7.9925 | 1.3302e4 | 0.0295 | 525.0528 | 0.0142 | 0.7850 | Base to Base |
| 17 | 9.4242 | 2.7870e4 | 0.0618 | 681.5675 | 0.0184 | 1.2317 | Base to Base |

**Figure S37**. HPLC spectrum of compound 1050-3.

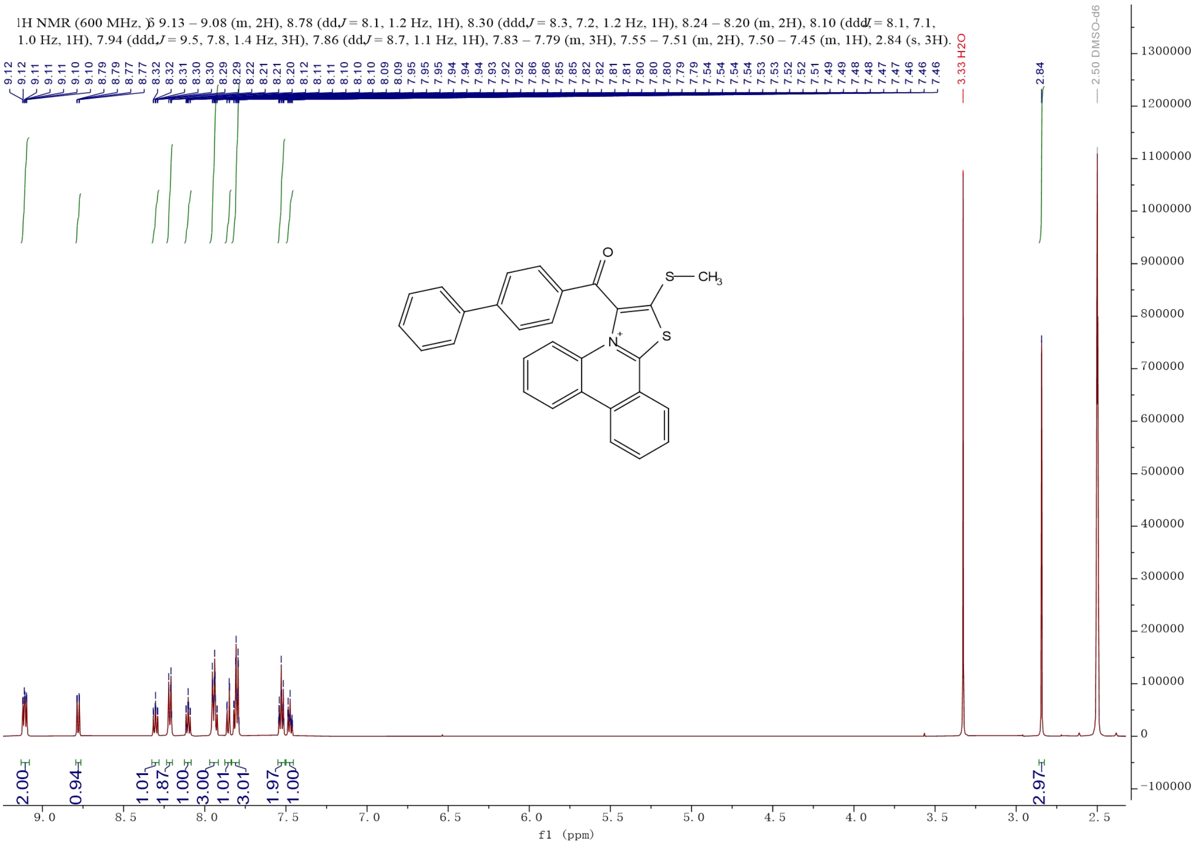


**Figure S38**. $^{1}$H NMR of compound 1050-4.

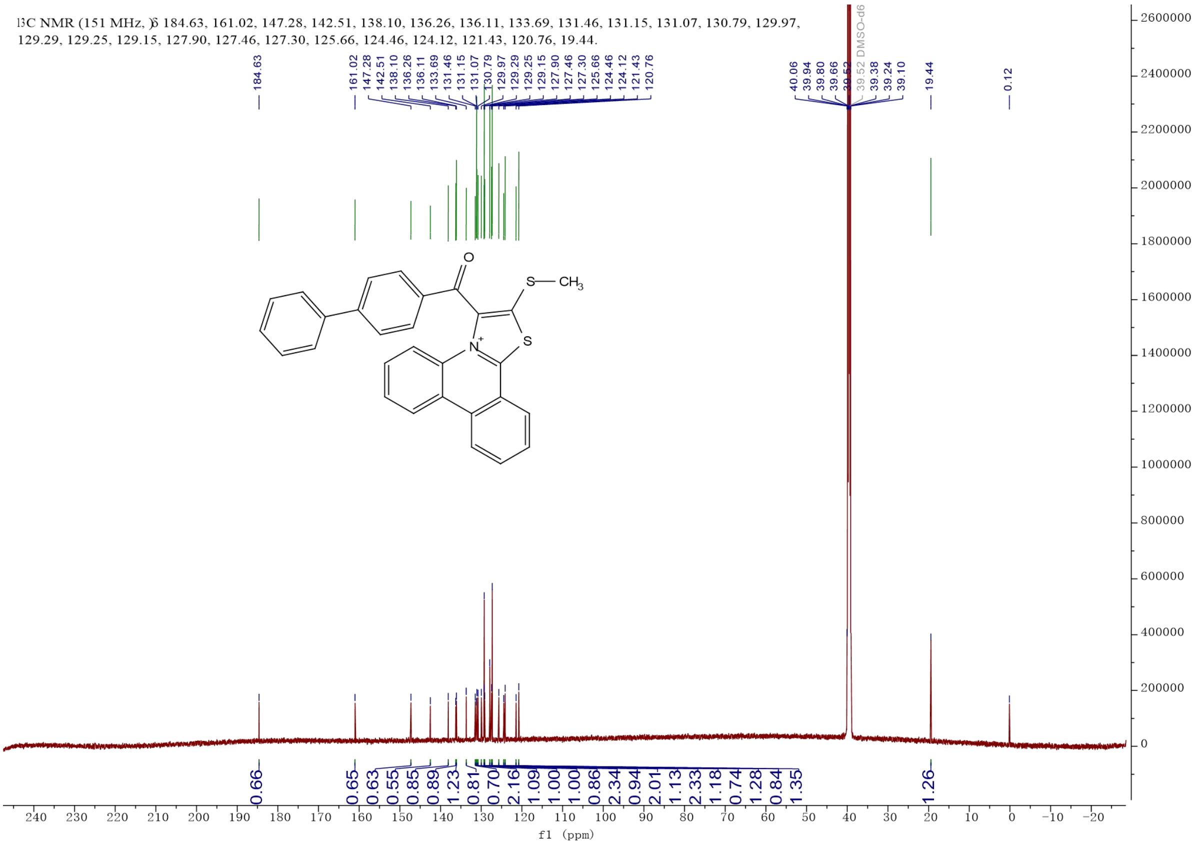


**Figure S39**. $^{13}$C NMR of compound 1050-4.

### Peking University Mass Spectrometry Sample Analysis Report

**Analysis Info**

| | | | |
|---|---|---|---|
| Analysis Name | FMTS-25010074_Pos_20250120_000003.d | Acquisition Date | 1/20/2025 11:10:51 AM |
| Sample | 1050-4 | Instrument | Bruker Solarix XR FTMS |
| Comment | | Operator | Peking University |

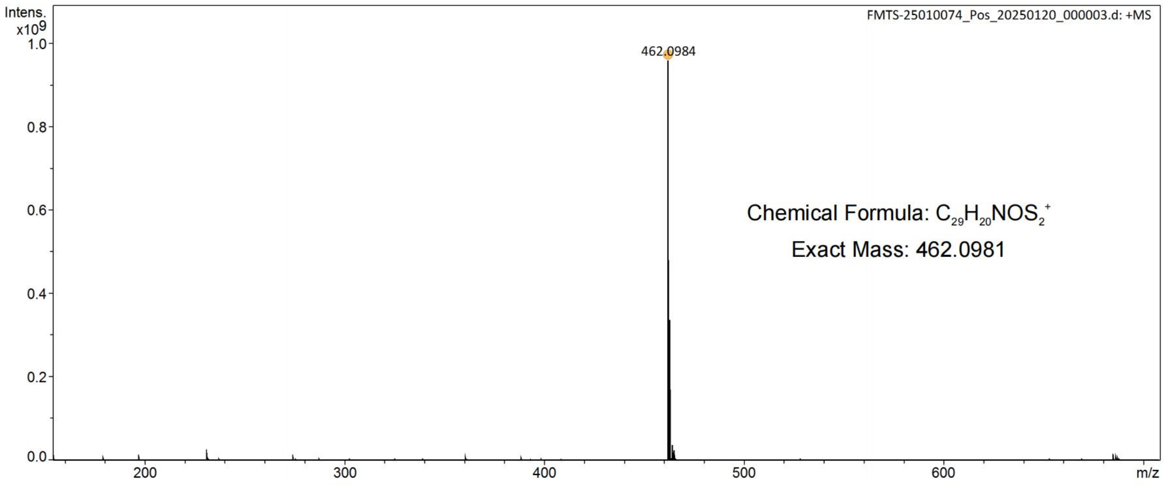


| Meas. m/z | # | Ion Formula | Score | m/z | err [ppm] | Mean err [ppm] | mSigma | rdb | e⁻ Conf | N-Rule |
|---|---|---|---|---|---|---|---|---|---|---|
| 462.0984 | 1 | C29H20NOS2 | 100.00 | 462.0981 | -0.8 | -0.7 | 23.7 | 25.0 | even | ok |

**Figure S40**. High-resolution mass spectrometry spectrum of compound 1050-4.

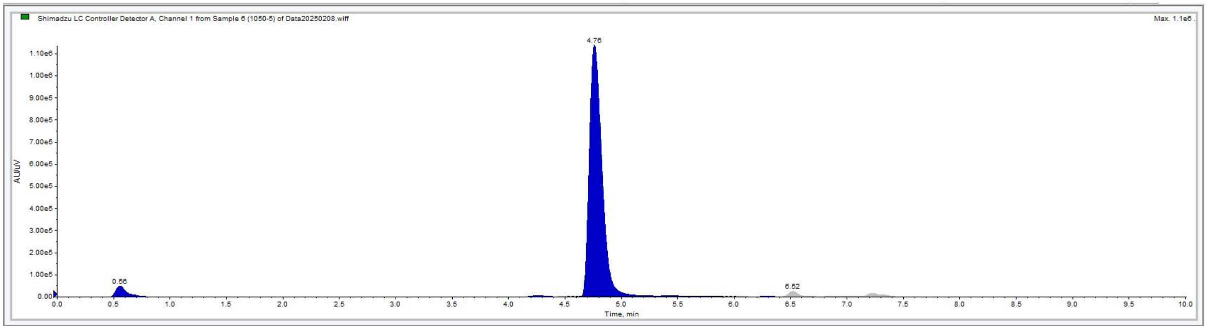


Shimadzu LC Controller Detector A, Channel 1 from Sample 6 (1050-5) of Data20250208.wiff

Data List Peak List

| | Time (min) | Area (counts) | % Area | Height | % Height | Width (min) | Baseline Type |
|---|---|---|---|---|---|---|---|
| 1 | 0.0742 | 2320.6500 | 0.0242 | 386.3138 | 0.0316 | 0.1633 | Base to Base |
| 2 | 0.2292 | 5332.1500 | 0.0556 | 756.8529 | 0.0618 | 0.2833 | Base to Base |
| 3 | 0.5564 | 3.5476e5 | 3.6978 | 4.9288e4 | 4.0264 | 0.5267 | Base to Base |
| 4 | 1.4142 | 8586.7000 | 0.0895 | 512.9296 | 0.0419 | 0.6183 | Base to Base |
| 5 | 3.6608 | 7193.8000 | 0.0750 | 866.2250 | 0.0708 | 0.2917 | Base to Base |
| 6 | 3.9608 | 2757.2000 | 0.0287 | 437.3852 | 0.0357 | 0.2050 | Base to Base |
| 7 | 4.2592 | 4.1098e4 | 0.4284 | 5555.0928 | 0.4538 | 0.3367 | Base to Base |
| 8 | 4.5083 | 1.3615e4 | 0.1419 | 2010.8750 | 0.1643 | 0.2133 | Base to Base |
| 9 | 4.7621 | 8.8002e6 | 91.7273 | 1.1373e6 | 92.9085 | 1.2567 | Base to Base |
| 10 | 5.9608 | 7808.8000 | 0.0814 | 860.7972 | 0.0703 | 0.2650 | Base to Base |
| 11 | 6.2992 | 4901.7000 | 0.0511 | 865.7090 | 0.0707 | 0.2183 | Base to Base |
| 12 | 6.5178 | 3.0842e5 | 3.2147 | 2.4232e4 | 1.9795 | 1.1800 | Base to Base |
| 13 | 7.9725 | 1.0025e4 | 0.1045 | 379.7736 | 0.0310 | 0.7067 | Base to Base |
| 14 | 9.4433 | 2.6849e4 | 0.2799 | 656.5615 | 0.0536 | 1.2200 | Base to Base |

**Figure S41**. HPLC spectrum of compound 1050-4.

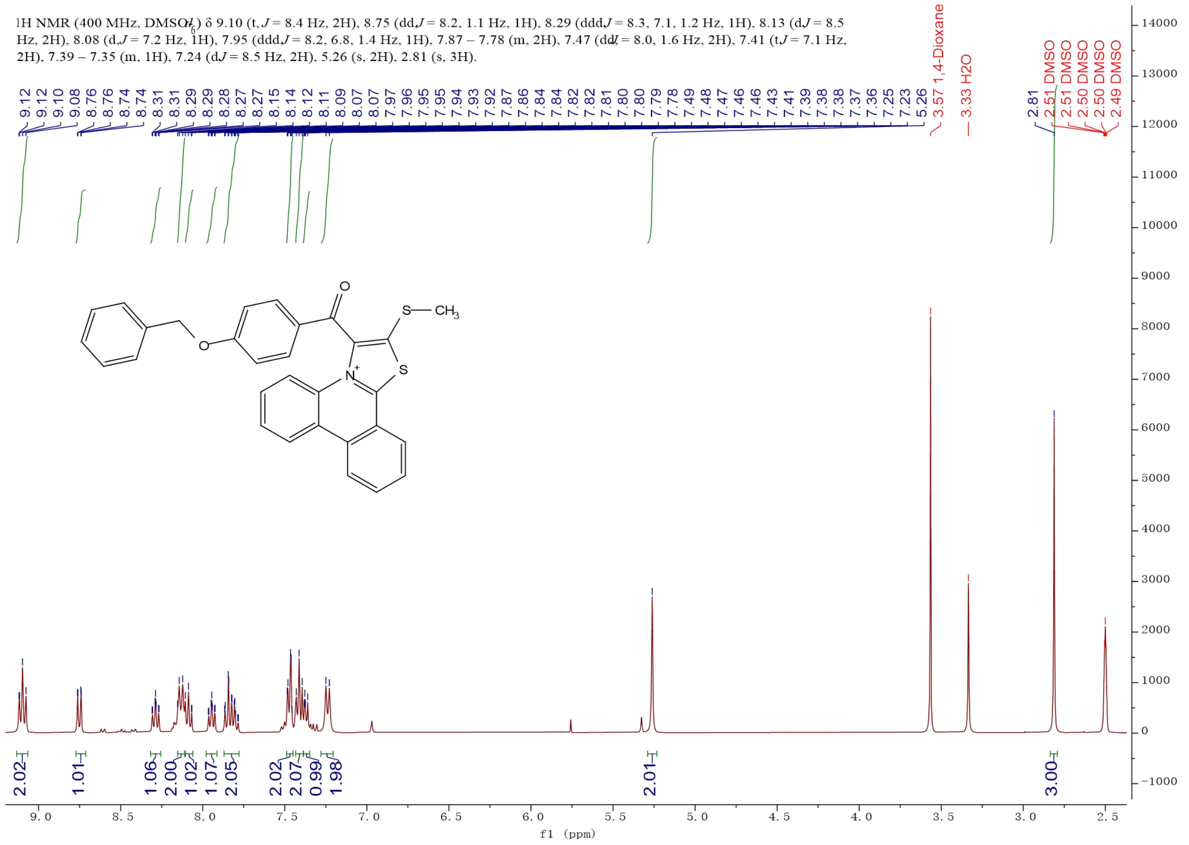


**Figure S42**. $^1$H NMR of compound 1050-5.

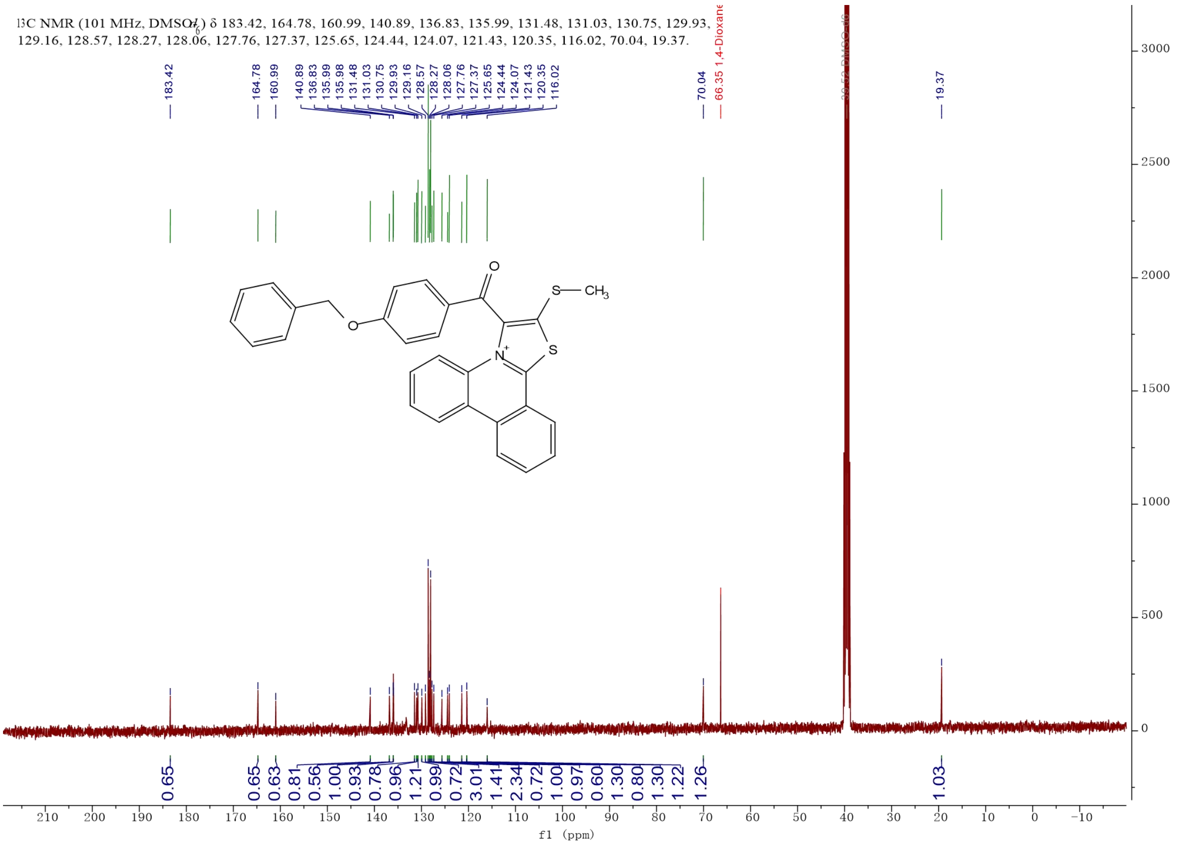


**Figure S43**. $^{13}$C NMR of compound 1050-5.

### Peking University Mass Spectrometry Sample Analysis Report

**Analysis Info**

| | | | |
|---|---|---|---|
| Analysis Name | FMTS-25010074_Pos_20250120_000004.d | Acquisition Date | 1/20/2025 11:13:14 AM |
| Sample | 1050-5 | Instrument | Bruker Solarix XR FTMS |
| Comment | | Operator | Peking University |

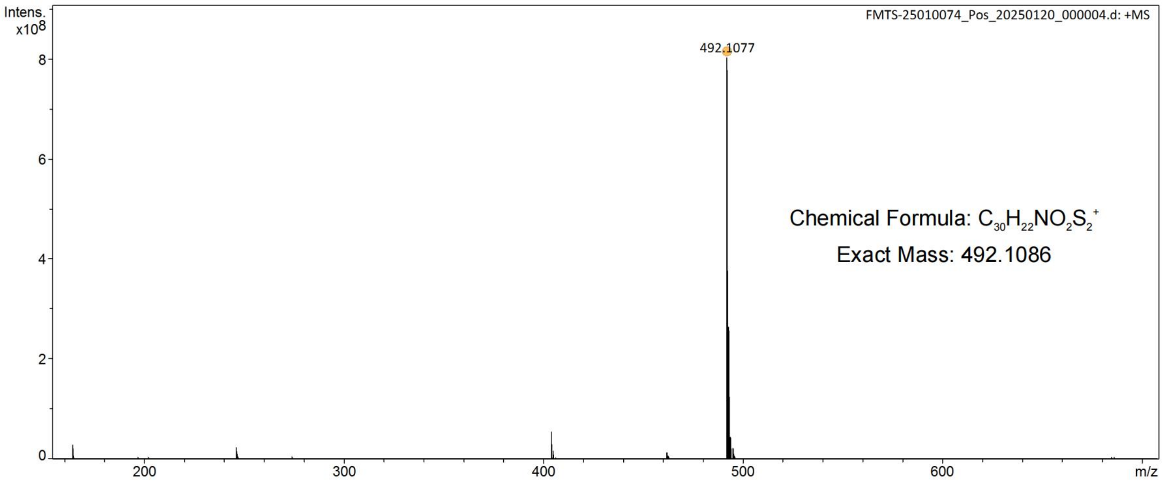


| Meas. m/z | # | Ion Formula | Score | m/z | err [ppm] | Mean err [ppm] | mSigma | rdb | e¯ Conf | N-Rule |
|---|---|---|---|---|---|---|---|---|---|---|
| 492.1077 | 1 | C30H22NO2S2 | 100.00 | 492.1086 | 2.0 | 2.5 | 49.3 | 25.0 | even | ok |

**Figure S44**. High-resolution mass spectrometry spectrum of compound 1050-5.

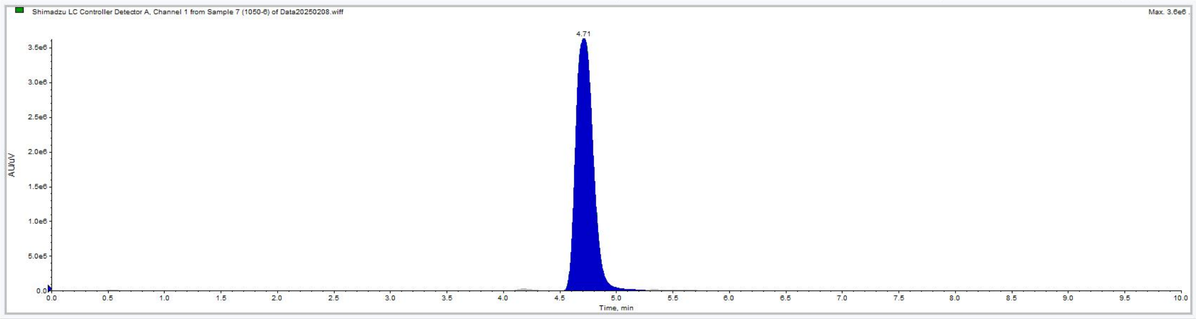


Shimadzu LC Controller Detector A, Channel 1 from Sample 7 (1050-6) of Data20250208.wiff

Data List | Peak List

| | Time (min) | Area (counts) | % Area | Height | % Height | Width (min) | Baseline Type |
|---|---|---|---|---|---|---|---|
| 1 | 0.2942 | 1.6135e4 | 0.0404 | 1344.0441 | 0.0362 | 0.3483 | Valley |
| 2 | 0.5301 | 1.4046e5 | 0.3515 | 1.8579e4 | 0.5007 | 0.5167 | Valley |
| 3 | 1.3942 | 9715.3000 | 0.0243 | 563.9010 | 0.0152 | 0.5283 | Base to Base |
| 4 | 3.2558 | 1985.7500 | 4.9690e-3 | 259.6031 | 6.9961e-3 | 0.2667 | Base to Base |
| 5 | 3.7531 | 2.7388e4 | 0.0685 | 2909.7575 | 0.0784 | 0.4067 | Base to Base |
| 6 | 4.1757 | 2.6440e5 | 0.6616 | 2.9140e4 | 0.7853 | 0.3833 | Base to Base |
| 7 | 4.7127 | 3.9013e7 | 97.6240 | 3.6334e6 | 97.9181 | 0.8250 | Valley |
| 8 | 5.3345 | 3.8868e5 | 0.9726 | 1.8014e4 | 0.4855 | 0.9550 | Valley |
| 9 | 6.9521 | 5.7958e4 | 0.1450 | 5241.1149 | 0.1412 | 1.0300 | Base to Base |
| 10 | 8.0775 | 1.5572e4 | 0.0390 | 555.6970 | 0.0150 | 0.8800 | Base to Base |
| 11 | 9.4025 | 2.7222e4 | 0.0681 | 646.7950 | 0.0174 | 1.2350 | Base to Base |

**Figure S45**. HPLC spectrum of compound 1050-5.

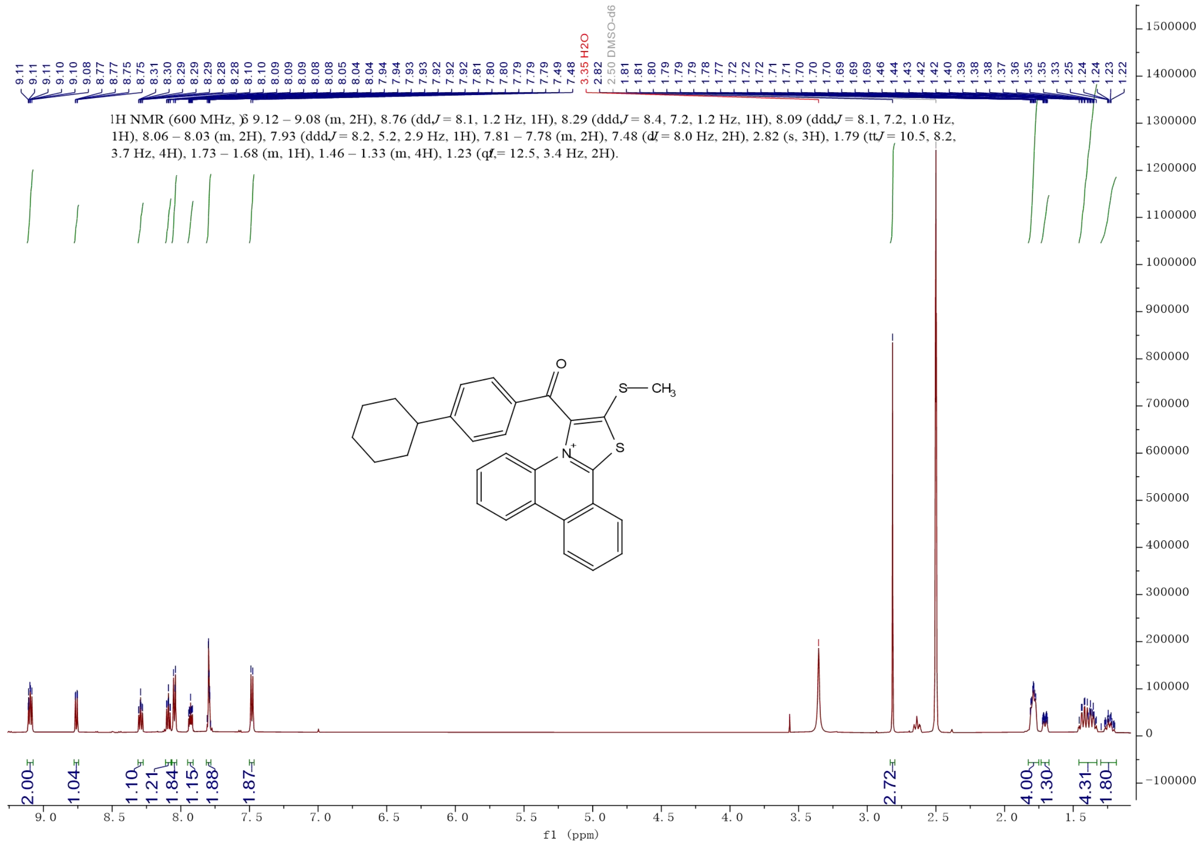


**Figure S46**. $^{1}$H NMR of compound 1050-6.

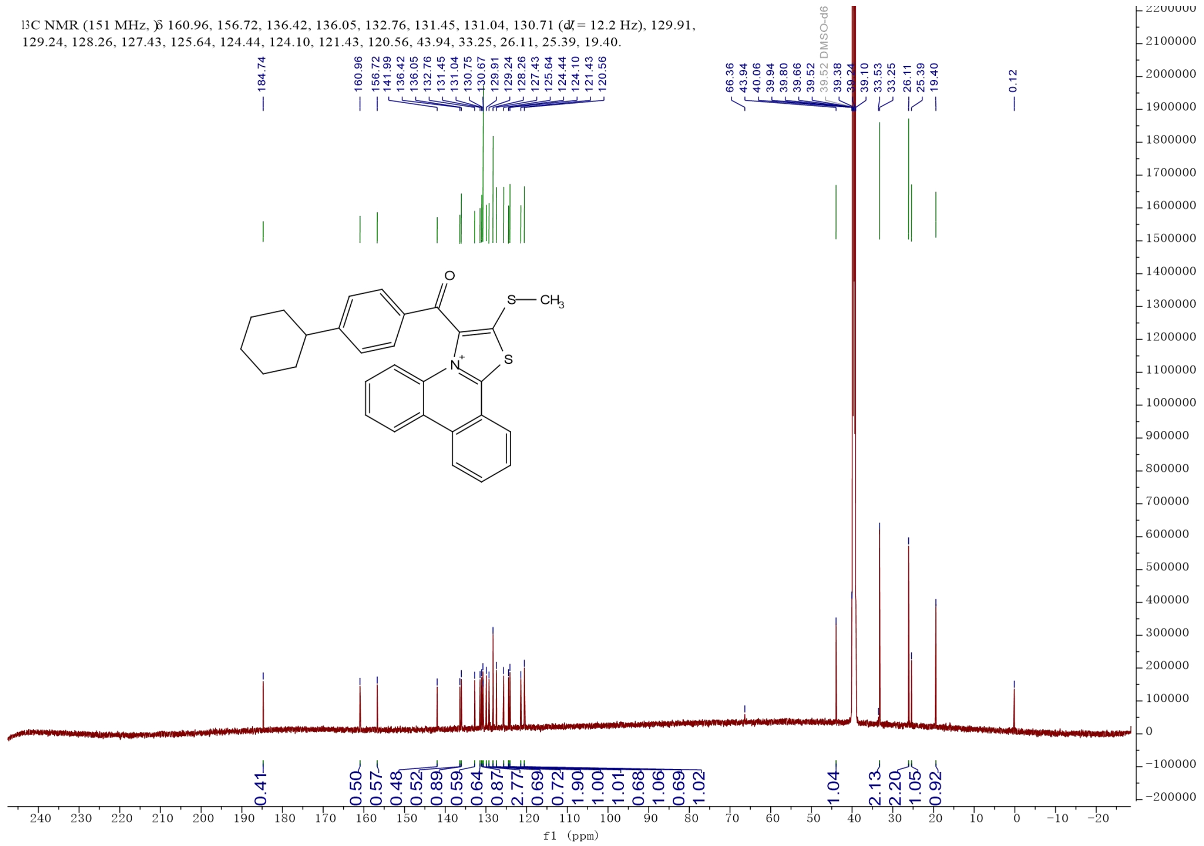


**Figure S47**. $^{13}$C NMR of compound 1050-6.

### Peking University Mass Spectrometry Sample Analysis Report

**Analysis Info**

| | | | |
|---|---|---|---|
| Analysis Name | FMTS-25010074_Pos_20250120_000006.d | Acquisition Date | 1/20/2025 11:16:59 AM |
| Sample | 1050-6 | Instrument | Bruker Solarix XR FTMS |
| Comment | | Operator | Peking University |

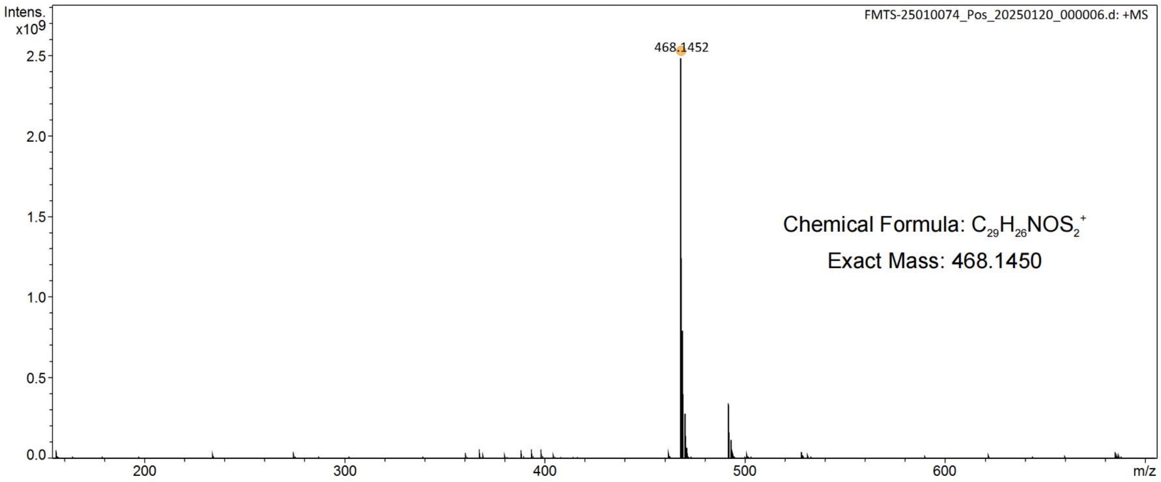


| Meas. m/z | # | Ion Formula | Score | m/z | err [ppm] | Mean err [ppm] | mSigma | rdb | e⁻ Conf | N-Rule |
|---|---|---|---|---|---|---|---|---|---|---|
| 468.1452 | 1 | C29H26NOS2 | 100.00 | 468.1450 | -0.3 | -7.7 | 25.8 | 22.0 | even | ok |

**Figure S48**. High-resolution mass spectrometry spectrum of compound 1050-6.

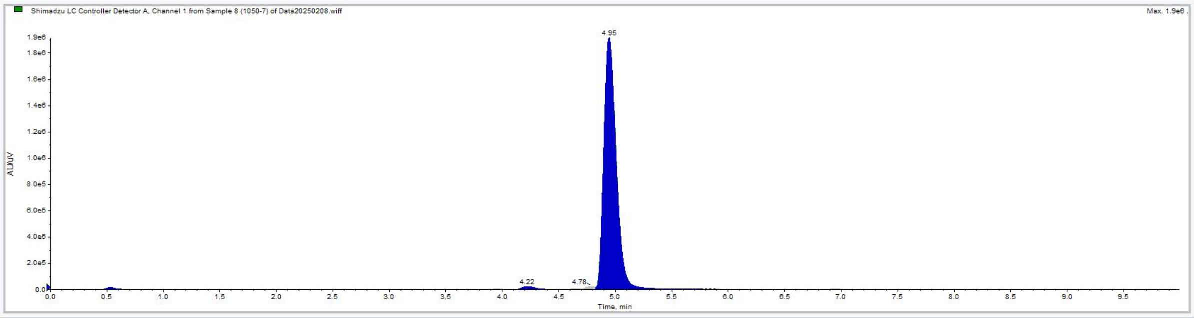


Shimadzu LC Controller Detector A, Channel 1 from Sample 8 (1050-7) of Data20250208.wiff

Data List Peak List

| | Time (min) | Area (counts) | % Area | Height | % Height | Width (min) | Baseline Type |
|---|---|---|---|---|---|---|---|
| 1 | 0.0700 | 1488.7000 | 9.8377e-3 | 301.3373 | 0.0151 | 0.1383 | Base to Base |
| 2 | 0.1958 | 5034.5000 | 0.0333 | 787.4664 | 0.0396 | 0.2917 | Base to Base |
| 3 | 0.5313 | 1.2883e5 | 0.8513 | 1.7127e4 | 0.8609 | 0.5317 | Base to Base |
| 4 | 1.3858 | 7971.1000 | 0.0527 | 442.8667 | 0.0223 | 0.6517 | Base to Base |
| 5 | 2.3675 | 3800.8500 | 0.0251 | 436.4737 | 0.0219 | 0.3800 | Base to Base |
| 6 | 2.9342 | 1.9987e4 | 0.1321 | 2383.4306 | 0.1198 | 0.3600 | Base to Base |
| 7 | 3.2975 | 9136.1500 | 0.0604 | 481.3310 | 0.0242 | 0.5867 | Base to Base |
| 8 | 3.9592 | 2.4140e4 | 0.1595 | 3381.1068 | 0.1700 | 0.2517 | Base to Base |
| 9 | 4.2238 | 1.6594e5 | 1.0966 | 2.2181e4 | 1.1149 | 0.3633 | Base to Base |
| 10 | 4.7834 | 1.1841e5 | 0.7825 | 1.9276e4 | 0.9689 | 0.2600 | Valley |
| 11 | 4.9451 | 1.4534e7 | 96.0411 | 1.9137e6 | 96.1916 | 0.9283 | Valley |
| 12 | 5.8558 | 1.0383e4 | 0.0686 | 1203.1693 | 0.0605 | 0.2633 | Base to Base |
| 13 | 6.1658 | 3825.2500 | 0.0253 | 769.8545 | 0.0387 | 0.1833 | Base to Base |
| 14 | 6.6858 | 7011.9651 | 0.0463 | 1039.0313 | 0.0522 | 0.2017 | Valley |
| 15 | 6.8258 | 9156.1498 | 0.0605 | 1673.0154 | 0.0841 | 0.1117 | Valley |
| 16 | 6.9417 | 4.3287e4 | 0.2860 | 3088.9468 | 0.1553 | 0.6967 | Valley |
| 17 | 7.9708 | 1.4848e4 | 0.0981 | 556.6466 | 0.0280 | 0.8100 | Base to Base |
| 18 | 9.3892 | 2.5843e4 | 0.1708 | 637.1841 | 0.0320 | 1.2117 | Base to Base |

**Figure S49**. HPLC spectrum of compound 1050-6.

**Table S1**. Spatial stability of binding residues in each binding pattern of the p53TAD1-1050 complex. The interaction type and interacting ligand group of each residue were selected by the maximum binding probability across all interactions the residue engagedSpatial stability of binding residues in each binding pattern of the p53TAD1-1050 complex. The interaction type and interacting ligand group of each residue were selected by the maximum binding probability across all interactions the residue engaged in.

| binding pattern | interaction type | ligand group | residue | RMSF | $RMSF_{norm}$ |
|---|---|---|---|---|---|
| BP1 | hydro | g4 | 12 | 0.47 | 0.08 |
| BP1 | electro | g0 | 16 | 0.25 | 0.00 |
| BP1 | hydro | g5 | 19 | 0.60 | 0.13 |
| BP1 | hydro | g1 | 23 | 1.21 | 0.35 |
| BP1 | hydro | g1 | 35 | 1.19 | 0.34 |
| BP1 | hydro | g2 | 36 | 1.83 | 0.58 |
| BP1 | hydro | g4 | 38 | 2.05 | 0.66 |
| BP1 | hydro | g4 | 39 | 2.99 | 1.00 |
| BP2 | hydro | g4 | 16 | 0.52 | 0.00 |
| BP2 | hydro | g4 | 17 | 0.92 | 0.46 |
| BP2 | hydro | g2 | 19 | 0.89 | 0.42 |
| BP2 | hydro | g2 | 23 | 0.67 | 0.17 |
| BP2 | hydro | g1 | 35 | 0.92 | 0.46 |
| BP2 | hydro | g1 | 36 | 1.40 | 1.00 |
| BP3 | hydro | g1 | 23 | 1.50 | 0.59 |
| BP3 | hydro | g1 | 25 | 2.28 | 1.00 |
| BP3 | hydro | g5 | 29 | 1.70 | 0.70 |
| BP3 | electro | g1 | 30 | 0.98 | 0.32 |
| BP3 | electro | g0 | 31 | 0.35 | 0.00 |
| BP3 | hydro | g4 | 32 | 0.53 | 0.09 |
| BP3 | hydro | g3 | 34 | 0.63 | 0.15 |
| BP3 | hydro | g2 | 35 | 1.07 | 0.37 |
| BP4 | hydro | g1 | 14 | 4.72 | 1.00 |
| BP4 | hydro | g1 | 18 | 3.06 | 0.61 |
| BP4 | hydro | g1 | 19 | 3.80 | 0.78 |
| BP4 | hydro | g5 | 22 | 2.05 | 0.38 |
| BP4 | hydro | g5 | 23 | 1.39 | 0.22 |
| BP4 | hydro | g4 | 32 | 1.23 | 0.19 |
| BP4 | electro | g0 | 33 | 0.43 | 0.00 |
| BP4 | hydro | g3 | 34 | 0.55 | 0.03 |
| BP4 | hydro | g1 | 36 | 1.99 | 0.36 |
| BP5 | hydro | g1 | 7 | 3.39 | 1.00 |
| BP5 | hydro | g1 | 8 | 3.11 | 0.91 |
| BP5 | hydro | g1 | 12 | 0.65 | 0.11 |
| BP5 | electro | g0 | 16 | 0.31 | 0.00 |
| BP5 | hydro | g4 | 19 | 0.90 | 0.19 |
| BP5 | hydro | g5 | 38 | 1.12 | 0.26 |
| BP6 | hydro | g1 | 23 | 0.80 | 1.00 |
| BP6 | electro | g0 | 24 | 0.49 | 0.34 |
| BP6 | hydro | g5 | 25 | 0.41 | 0.18 |
| BP6 | electro | g0 | 26 | 0.33 | 0.00 |
| BP6 | hydro | g4 | 27 | 0.78 | 0.97 |

| | | | | | |
|---|---|---|---|---|---|
| BP6 | hydro | g2 | 28 | 0.70 | 0.78 |
| BP7 | hydro | g4 | 7 | 2.59 | 1.00 |
| BP7 | hydro | g4 | 8 | 2.16 | 0.80 |
| BP7 | hydro | g1 | 12 | 1.10 | 0.29 |
| BP7 | hydro | g1 | 13 | 1.30 | 0.38 |
| BP7 | hydro | g1 | 19 | 0.90 | 0.19 |
| BP7 | hydro | g5 | 22 | 0.50 | 0.00 |
| BP7 | hydro | g5 | 23 | 1.08 | 0.28 |
| BP7 | hydro | g5 | 26 | 0.53 | 0.01 |
| BP7 | hydro | g5 | 28 | 0.70 | 0.10 |
| BP7 | hydro | g4 | 29 | 0.61 | 0.05 |
| BP7 | hydro | g4 | 32 | 1.96 | 0.70 |

**Table S2**. Spatial stability of binding residues in each binding pattern of the p53TAD1-1050-6 complex. The interaction type and interacting ligand group of each residue were selected by the maximum binding probability across all interactions the residue engaged in.

| binding pattern | interaction type | ligand group | residue | RMSF | $RMSF_{norm}$ |
|---|---|---|---|---|---|
| BP1 | hydro | g4 | 4 | 7.59 | 0.82 |
| BP1 | hydro | g6 | 19 | 9.06 | 1.00 |
| BP1 | hydro | g6 | 22 | 8.77 | 0.96 |
| BP1 | hydro | g6 | 23 | 7.22 | 0.77 |
| BP1 | hydro | g1 | 25 | 4.54 | 0.43 |
| BP1 | hydro | g5 | 30 | 1.42 | 0.04 |
| BP1 | electro | g0 | 31 | 1.10 | 0.00 |
| BP1 | hydro | g4 | 32 | 1.68 | 0.07 |
| BP1 | hydro | g2 | 34 | 1.72 | 0.08 |
| BP1 | hydro | g2 | 35 | 2.19 | 0.14 |
| BP2 | hydro | g4 | 4 | 7.26 | 1.00 |
| BP2 | hydro | g3 | 10 | 4.35 | 0.56 |
| BP2 | hydro | g2 | 12 | 2.80 | 0.32 |
| BP2 | hydro | g6 | 19 | 0.84 | 0.02 |
| BP2 | hydro | g6 | 23 | 0.71 | 0.00 |
| BP2 | hydro | g5 | 31 | 2.84 | 0.33 |
| BP2 | hydro | g1 | 32 | 1.39 | 0.10 |
| BP2 | hydro | g2 | 34 | 2.24 | 0.23 |
| BP3 | electro | g0 | 2 | 8.04 | 0.97 |
| BP3 | hydro | g6 | 4 | 8.25 | 1.00 |
| BP3 | electro | g0 | 7 | 4.65 | 0.53 |
| BP3 | hydro | g2 | 10 | 3.65 | 0.41 |
| BP3 | hydro | g6 | 12 | 5.20 | 0.61 |
| BP3 | hydro | g6 | 13 | 5.77 | 0.68 |
| BP3 | hydro | g6 | 14 | 5.69 | 0.67 |
| BP3 | hydro | g6 | 23 | 6.68 | 0.80 |
| BP3 | hydro | g1 | 25 | 4.53 | 0.52 |
| BP3 | Hbond | g1 | 30 | 0.79 | 0.04 |
| BP3 | electro | g0 | 31 | 0.51 | 0.00 |
| BP3 | hydro | g5 | 32 | 1.21 | 0.09 |

| | | | | | |
|---|---|---|---|---|---|
| BP3 | hydro | g4 | 34 | 1.16 | 0.08 |
| BP3 | hydro | g2 | 35 | 2.43 | 0.25 |
| BP4 | hydro | g4 | 3 | 3.43 | 0.72 |
| BP4 | hydro | g4 | 4 | 3.23 | 0.67 |
| BP4 | hydro | g2 | 8 | 1.74 | 0.32 |
| BP4 | hydro | g4 | 13 | 0.48 | 0.02 |
| BP4 | hydro | g4 | 16 | 0.38 | 0.00 |
| BP4 | hydro | g4 | 17 | 0.55 | 0.04 |
| BP4 | hydro | g1 | 23 | 1.11 | 0.17 |
| BP4 | hydro | g6 | 26 | 1.60 | 0.29 |
| BP4 | hydro | g2 | 27 | 3.64 | 0.77 |
| BP4 | hydro | g6 | 31 | 4.31 | 0.93 |
| BP4 | hydro | g6 | 32 | 4.62 | 1.00 |
| BP5 | hydro | g4 | 1 | 1.73 | 0.53 |
| BP5 | hydro | g5 | 2 | 1.04 | 0.20 |
| BP5 | hydro | g4 | 3 | 1.47 | 0.41 |
| BP5 | hydro | g1 | 4 | 1.47 | 0.41 |
| BP5 | hydro | g2 | 8 | 0.61 | 0.00 |
| BP5 | hydro | g2 | 18 | 1.00 | 0.18 |
| BP5 | hydro | g2 | 19 | 1.01 | 0.19 |
| BP5 | hydro | g6 | 21 | 2.72 | 1.00 |
| BP5 | hydro | g2 | 22 | 0.83 | 0.10 |
| BP5 | hydro | g6 | 25 | 1.89 | 0.60 |
| BP5 | hydro | g6 | 31 | 1.73 | 0.53 |
| BP6 | hydro | g4 | 14 | 1.07 | 1.00 |
| BP6 | electro | g0 | 15 | 0.43 | 0.00 |
| BP6 | hydro | g1 | 17 | 0.51 | 0.12 |
| BP6 | hydro | g6 | 19 | 0.70 | 0.42 |
| BP7 | hydro | g1 | 12 | 2.10 | 0.31 |
| BP7 | hydro | g5 | 13 | 2.69 | 0.43 |
| BP7 | hydro | g6 | 14 | 2.46 | 0.39 |
| BP7 | electro | g0 | 17 | 5.42 | 1.00 |
| BP7 | hydro | g6 | 19 | 3.14 | 0.53 |
| BP7 | hydro | g2 | 25 | 0.88 | 0.06 |
| BP7 | hydro | g2 | 30 | 0.80 | 0.04 |
| BP7 | electro | g0 | 31 | 1.24 | 0.13 |
| BP7 | hydro | g1 | 32 | 0.60 | 0.00 |
| BP7 | hydro | g5 | 34 | 0.90 | 0.06 |
| BP7 | hydro | g4 | 35 | 1.72 | 0.23 |
| BP8 | electro | g0 | 3 | 6.93 | 1.00 |
| BP8 | hydro | g4 | 4 | 5.71 | 0.81 |
| BP8 | hydro | g4 | 11 | 2.46 | 0.32 |
| BP8 | hydro | g4 | 12 | 1.60 | 0.18 |
| BP8 | hydro | g4 | 32 | 2.27 | 0.29 |
| BP8 | hydro | g5 | 34 | 0.55 | 0.02 |
| BP8 | Hbond | g1 | 35 | 0.41 | 0.00 |
| BP8 | hydro | g1 | 36 | 0.88 | 0.07 |
| BP9 | hydro | g2 | 19 | 3.27 | 1.00 |
| BP9 | hydro | g2 | 21 | 0.69 | 0.08 |
| BP9 | hydro | g4 | 22 | 1.32 | 0.31 |

| | | | | | |
|---|---|---|---|---|---|
| BP9 | hydro | g6 | 24 | 0.67 | 0.07 |
| BP9 | hydro | g1 | 25 | 0.46 | 0.00 |
| BP9 | hydro | g5 | 27 | 0.78 | 0.11 |
| BP10 | hydro | g4 | 2 | 0.72 | 0.12 |
| BP10 | electro | g0 | 4 | 0.89 | 0.20 |
| BP10 | hydro | g2 | 8 | 1.28 | 0.38 |
| BP10 | hydro | g2 | 18 | 1.13 | 0.31 |
| BP10 | hydro | g6 | 19 | 0.46 | 0.00 |
| BP10 | hydro | g1 | 22 | 0.49 | 0.01 |
| BP10 | hydro | g6 | 23 | 0.70 | 0.11 |
| BP10 | Hbond | g1 | 30 | 1.55 | 0.50 |
| BP10 | hydro | g5 | 32 | 1.63 | 0.54 |
| BP10 | hydro | g4 | 34 | 2.62 | 1.00 |

Table S3. The parameters used for extracting binding patterns of AR-Tau5 and EPI-series ligands from the trajectories generated using REST2. A frame is defined as a contacting frame if it has at least four residues contacting the ligand. For AR-Tau5 systems, *L*=15 means that each window represents 1.2 ns.

| **System** | **Contacting frames** | **Total frames** | **Contact probability (%)** | ***L*** | ***minPts*** | $S_{core}$ | $S_{query}$ |
|---|---|---|---|---|---|---|---|
| BADGE | 18999 | 52988 | 35.9 | - | - | - | - |
| EPI-002 | 17355 | 49507 | 35.1 | 15 | 20 | 7 | 0.7 |
| EPI-7170 | 33858 | 55157 | 61.4 | - | - | - | - |
| iodoEPI-002 | 40780 | 65491 | 62.3 | 15 | 60 | 7 | 0.7 |
